# The small molecule ML233 is a direct inhibitor of tyrosinase function

**DOI:** 10.1101/2025.02.16.638443

**Authors:** Romain Menard, Aissette Baanannou, Caroline Halluin, Dexter Morse, Sadie Kuhn, Joel H. Graber, James Strickland, Romain Madelaine

## Abstract

Melanogenesis is the biological process regulating the synthesis of melanin pigments in melanocytes. Defective melanogenesis is associated with numerous human skin diseases, including, but not limited to, albinism, vitiligo, melasma, and hypo- and hyperpigmentation disorders. Tyrosinase is the rate-limiting enzyme controlling melanogenesis, and hence tremendous efforts have been made to identify potent and safe inhibitors of tyrosinase function. However, despite decades of research, currently there is no effective treatment that inhibits melanogenesis or tyrosinase activity with no adverse side effects. In this study, we report characterization of the ML233 chemical as a potent inhibitor of tyrosinase activity *in vivo* and *in vitro*. We demonstrate that ML233 reduces melanin production in the zebrafish model with no observable significant toxic side effects, and in murine melanoma cells. We also predict that these effects are mediated through direct tyrosinase-ML233 interaction, i.e., binding of the ML233 molecule to the active site of the protein to inhibit its function. Together, our results reveal that ML233 plays roles in both healthy and pathological skin cells via inhibition of melanin production. ML233-mediated tyrosinase inhibition is a potentially safe and effective approach to alleviate the symptoms of melanocyte-associated diseases and thereby substantially improve human health.

## INTRODUCTION

Skin pigmentation has evolved to play different roles across evolution. It is involved in protection against UV rays, attraction for mating, and camouflage via background adaptation. A conserved feature of skin coloring is the production of pigments from specialized cells termed melanocytes^1,2^. Melanocytes contain cellular organelles termed melanosomes that produce melanin, a natural skin pigment responsible for coloring of the skin, eyes, and hair in humans. Defects in melanocyte function or melanin production can lead to skin-pigmentation disorders, including vitiligo, albinism, and melanoma^3–6^.

In fish, amphibians, reptiles, and other vertebrates, eumelanin, a black pigment, is produced by a specific type of melanocytes termed melanophores. Because the genetic pathways regulating melanocyte and melanophore function are highly conserved across vertebrates, model organisms including zebrafish are biologically relevant and valuable models for uncovering the molecular mechanisms regulating melanocyte function and underlying melanin-related diseases in humans^7–12^. During embryonic vertebrate development, expression of the transcription factor SOX10 defines a neural crest population that gives rise to the melanocytes^13^. A downstream target of SOX10, the MITF gene (microphthalmia-associated transcription factor), promotes melanocyte differentiation^14,15^. In addition, MITF controls the expression of genes essential for melanin synthesis^16^, including tyrosinase (TYR), tyrosinase-related proteins (TYRP), and dopachrome tautomerase (DCT).

The biological process regulating melanin synthesis, termed melanogenesis, is achieved through a complex series of enzymatic and chemical reactions^17–19^. The rate-limiting enzyme involved in this process is TYR, as the other reactions occur spontaneously. During melanin synthesis, TYR first hydroxylates the amino acid tyrosine to L-DOPA (3,4-dihydroxyphenylalanine) and then oxidates L-DOPA to produce dopaquinone, which in turn is converted to DOPA (via oxidation again by TYR) and to dopachrome (via auto-oxidation). Finally, reaction products from dopachrome or dopaquinone are used in two different metabolic pathways to produce two types of melanin: the black–brown eumelanin and the yellow–red pheomelanin.

The amount of melanin in pigment cells is highly important for pigment-cell homeostasis. Melanin protects the skin from UV exposure-generated DNA damage and reactive oxygen species production^20,21^. Little or no melanin production is associated with albinism in humans^4^, an inherited skin condition that leads to pale skin that is highly sensitive to sun exposure and is associated with a high risk of skin cancer. Vitiligo syndrome, a pale skin condition, is characterized by the loss of melanocytes^3,6^. The genetic basis of vitiligo is still poorly understood, but recent evidence suggests an autoimmune component in the disease etiology. For example, it has been proposed that in vitiligo T lymphocytes against melanocyte antigens could lead to pigment-cell death^22^, and that autoimmune antibodies inhibiting melanin-concentrating hormone-receptor activity could lead to overexpression of α-melanin-stimulating hormone (α-MSH) and thereby affect pigment-cell survival ^23–25^. Together, the evidence above highlights how reducing or increasing levels of melanogenesis can both affect melanocyte homeostasis and underlie human skin diseases.

Although melanoma is not the most common cause of skin cancer, it is the most dangerous, as it is highly metastatic. Melanoma is caused by melanocytes with high proliferative and invasive properties, and it is currently believed that these properties result from genetic and epigenetic modifications that restore a multipotent neural crest-like state in differentiated melanocytes^26–28^. It has been suggested that inhibition of TYR activity and/or melanogenesis could be a promising therapeutic strategy to sensitize melanoma to chemotherapeutic treatments^29,30^.

Over the past 20 years, many melanogenesis inhibitors have been identified and tested, including, but not limited to, hydroquinone, kojic acid, azelaic acid, and arbutin^18,31–34^. Melanogenesis inhibitors, in particular, direct TYR inhibitors, are highly relevant both therapeutically and cosmetically, for their ability to maintain pigment-cell homeostasis and health or to induce skin-lightening. Unfortunately, one of the widely used skin-lightening agents that inhibit TYR activity, hydroquinone, causes skin irritation and has been suggested to be mutagenic to melanocytes^35–37^. Alternative skin-lightening products that inhibit TYR such as arbutin demonstrated relatively high efficiency *in vivo*, but because arbutin can be converted to hydroquinone by skin bacteria or the intestinal microbiome^38,39^, it may carry carcinogenic risk. In sum, there is a clear need to identify new anti-melanogenesis molecules with skin-lightening or anti-cancer properties that can be tested for efficiency and safety.

In the hunt to discover new melanogenesis inhibitors, the zebrafish is a recognized model organism because it alleviates the need for mammalian study during the early stages of drug efficiency and toxicity testing ^40^. The zebrafish provides countless advantages for drug screening and/or testing, including a large number of individuals, drug delivery via solubilization of the drug in the water of the fish, high penetration of the compound through the mouth, gills, and skin, and direct visualization of the pigmentation phenotype in embryos or larvae. In addition, zebrafish larvae harbor all internal organs present in adults, such as heart, kidney, and liver, indicating that we can also assess the organ-specific toxicity of a candidate molecule *in vivo*. Finally, because the molecular mechanisms of melanocyte differentiation and melanogenesis are highly conserved between mammals and zebrafish^8–10^, the molecules identified in zebrafish to be potent inhibitors of melanogenesis are strong candidates to be active in mammalian melanocytes as well ^41^.

In this study we used zebrafish embryos and larvae to analyze the effect of a small chemical molecule, ML233, in the control of skin pigmentation, melanogenesis, and TYR activity. ML233 was characterized in 2011 as a small chemical agonist of the apelin receptor (a.k.a. APJ, APLNR)^42^. ML233 has a potent binding activity on APJ in cell culture^42^. ML233 has also been validated *in vivo* in the zebrafish model, where loss- and gain-of-function of apelin receptor genes (*aplnr*a and *aplnrb*) demonstrated APLNR-dependent ML233 function in zebrafish embryos^43^. In the present study, we provide *in vivo* genetic evidence that ML233 also acts as an inhibitor of melanogenesis independently of the apelin signaling pathway. Importantly, we show that ML233 does not affect melanocyte development and that its effect on melanogenesis is reversible. In addition, *in vitro* experiments and computational modelling reveal that the mechanism of melanogenesis inhibition by ML233 is, at least partially, through direct inhibition of TYR activity and function. Finally, using murine and human melanoma cell lines, we demonstrate that ML233-dependent inhibition of melanogenesis is conserved in mammals, indicating that this small chemical molecule has therapeutic and cosmetic potential in humans.

## RESULTS

### The small chemical molecule ML233 has no impact on zebrafish embryo survival

ML233 is a small chemical molecule (C_19_H_21_NO_4_S) with a molecular weight of 359.44 (Fig. 1A). To better characterize the impact of ML233 activity on tissue and organ homeostasis in zebrafish, we first assessed the toxicity of the molecule. We performed an acute toxicity text by treating zebrafish eggs at different concentrations of ML233 ranging from 1.25 to 200 µM, beginning at 4 hpf (hours post-fertilization) (Fig. S1A, B). We observed relatively low toxicity in zebrafish embryos with a survival rate of ∼80% at 2 dpf (days post-fertilization) and an ML233 concentration between 7.5 and 200 µM. During the survival analysis we observed a precipitate in the medium at concentrations over 30 µM, raising the issue that ML233 may have low solubility. Based on this observation, we next used ML233 at a maximum concentration of 20 µM. Following the guidelines of the OECD 236 toxicity test on zebrafish embryos, we next confirmed the previous observations and showed a 100% survival rate at 2 dpf with a concentration of 20 µM (Fig. 1B, C). We then compared the efficiency and toxicity of ML233 with that of two broadly use skin-lightening agents, kojic acid and arbutin, determining that 10 and 200 µM of either kojic acid or arbutin has little effect on skin pigmentation (Fig. 1D and Fig. S1C). At a concentration of 400 µM, both kojic acid and arbutin have an incomplete impact on the reduction of skin pigmentation, while harboring significant side effects on eye size, process of hatching and embryo survival (Fig. 1D and Fig. S1C-F). In contrast, ML233 has no negative impact on survival or hatching at a concentration sufficient to reduce skin pigmentation, and shows only a slight, but significant, reduction of eye axial length at this concentration (Fig. S1C-F). The above results suggest that, of the three molecules tested, ML233 is the most potent and tolerable inhibitor of melanogenesis in zebrafish embryos. Finally, to calculate the LD50 of the ML233 molecule, we extended the survival analysis until 4 dpf, but we observed no increase in lethality during these two additional days (Fig. 1C). Together, these data indicate that use of ML233 at a concentration of 20 µM or lower has no significant toxic side effects on zebrafish during development.

**Figure 1.**
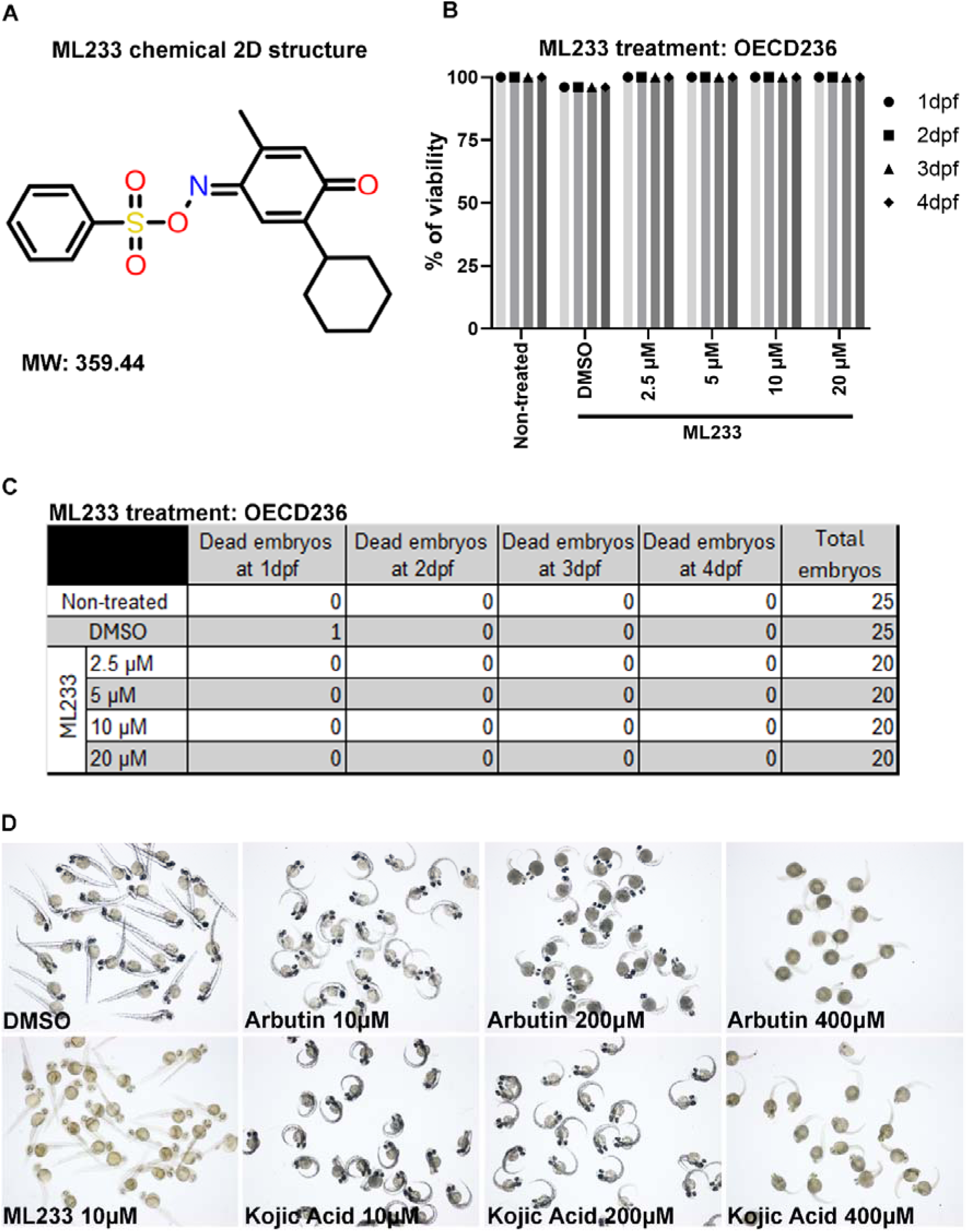
The ML233 molecule exhibits no detectable toxicity on zebrafish embryos. **(A)** 2D representative structure of the ML233 chemical (molecular weight: 359.44). **(B)** Zebrafish embryo survival (% of viability) was quantified in non-treated, control (DMSO) and ML233-treated embryos by OECD236 test to assess acute toxicity in fish embryos (n≥20). **(C)** Table showing the number of embryos used for the OECD236 test in (B) and the survival rate every day between 1 and 4 dpf. **(D)** Skin pigmentation of DMSO-, ML233-, arbutin-, or -kojic acid-treated embryos at 48 hpf after treatment between 4 and 48 hpf.

### ML233 inhibits melanin production and reduces skin pigmentation *in vivo*

During our testing of ML233 toxicity in zebrafish, we observed a striking reduction in skin pigmentation in embryos treated with 10 µM ML233 compared to embryos under DMSO-only treatment conditions (Fig. 1D), where both treatments began at 4 hpf. The extent of the reduced pigmentation phenotype correlates positively with increasing ML233 concentrations (Fig. 2A, B), suggesting a dose-dependent effect. *In vitro* absorbance measurement at 490nm revealed a significant reduction of melanin quantity at 2 dpf in embryos under ML233 treatment conditions (Fig. 2B), indicating potent inhibition of skin pigmentation. At ML233 concentrations as low as 0.5 µM, we observe a significant reduction of melanin quantity, i.e., ∼50% reduction. At 5 µM and higher concentrations of ML233, inhibition of melanogenesis is over 80%, a reduction similar to that observed in embryos at 2 dpf under treatment conditions using 200 µM of 1-phenyl 2-thiourea (PTU), a melanogenesis inhibitor widely used to prevent pigmentation of zebrafish embryos and larvae during development. Together, these results indicate that the ML233 small chemical molecule reduces skin pigmentation *in vivo*, and that the efficiency of melanogenesis inhibition is dependent on the chemical concentration.

**Figure 2.**
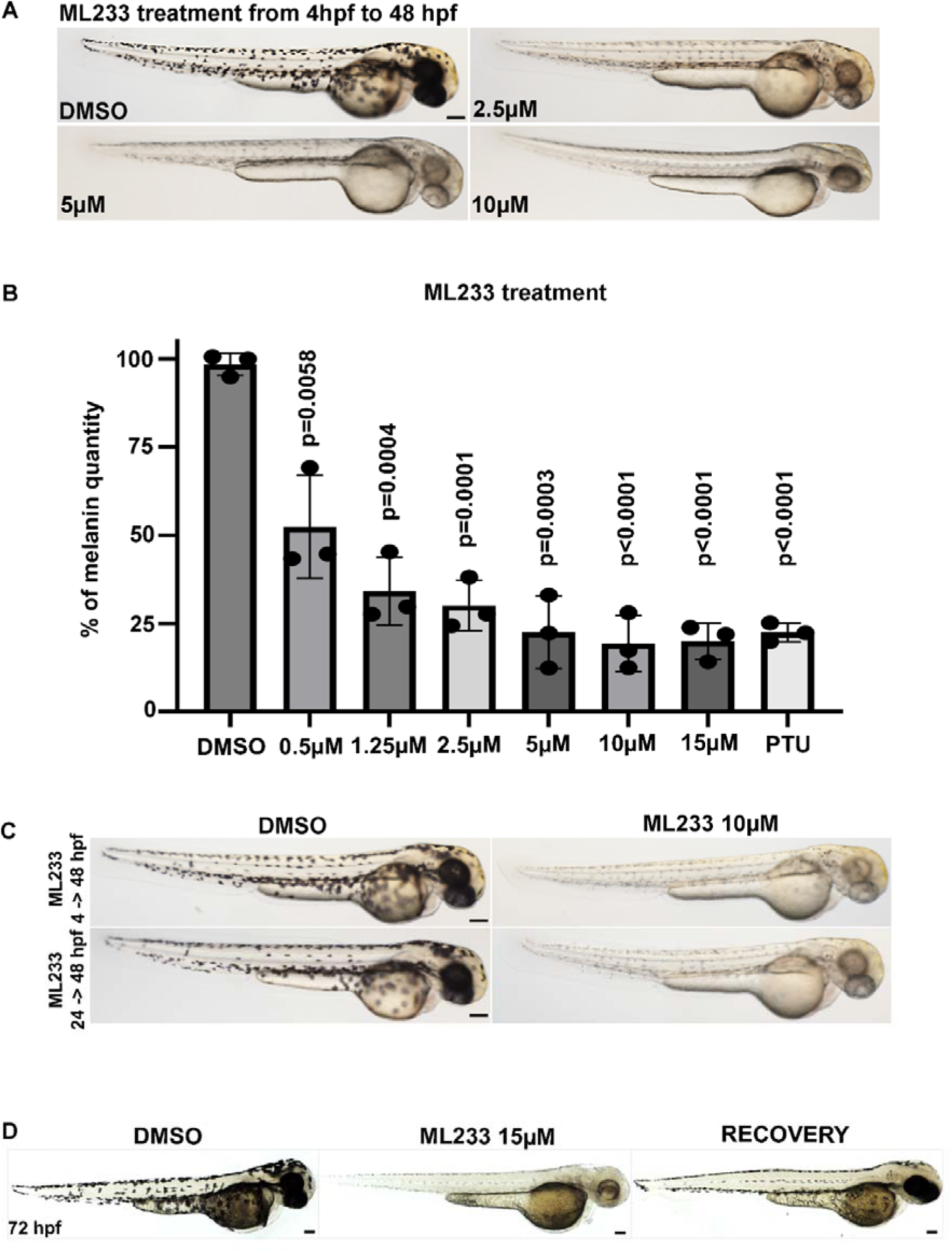
ML233 inhibits melanogenesis, and this effect is reversible. **(A)** Skin pigmentation of DMSO- or ML233-treated embryos at 48 hpf after treatment between 4 and 48 hpf. **(B)** Melanin was quantified in control or ML233-treated embryos (n=3 biological replicates with 3 technical replicates; ≥40 embryos/biological replicate). Significance is determined by t-test, two-tailed, unpaired. **(C)** Skin pigmentation of DMSO- or ML233-treated embryos at 48 hpf after treatment between 4 and 48 hpf (top) or 24 and 48 hpf (bottom). **(D)** Skin pigmentation of DMSO, ML233-treated embryos (treatment between 24 and 72 hpf) and ML233-treated embryos after recovery (treatment between 24 and 48 hpf) at 72 hpf. Scale bars: 100 μm. Error bars represent s.d.

### Reduction of skin pigmentation via ML233-induced melanogenesis inhibition is reversible

The above results led us to investigate the mechanism of ML233-dependent skin-pigmentation reduction. Because ML233 treatment at an early stage of development could potentially affect melanocyte formation in zebrafish embryos, we tested the effect of ML233 treatment starting not only at 4 hpf but also at 24 hpf, when melanocytes are specified^10,11^. Treatment with concentrations ranging from 2.5 to 15 µM between 24 and 48 hpf reveals that ML233 inhibits melanogenesis by skin melanocytes similarly to treatment with the same concentrations beginning at 4 hpf (Fig. 2C and Fig. S1G). The above results support the hypothesis that inhibition of skin pigmentation is mediated through inhibition of melanogenesis.

To rule out a potential negative effect of ML233 treatment on melanocyte survival, we next tested the reversibility of ML233 treatment. Zebrafish embryos were treated with 15 µM ML233 between 24 and 48 hpf. At 48 hpf, the water with chemical was removed, and embryos were washed in chemical-free medium and kept in chemical-free medium until 72 hpf. At 72 hpf, we observed increased melanogenesis by melanocytes in embryos treated with ML233 between 24 and 48 hpf, compared to embryos treated with ML233 until 72 hpf (Fig. 2D). Strikingly, eye pigmentation in embryos after 24 hours of recovery following ML233 treatment strongly resembles that of control DMSO-treated embryos. These observations suggest that melanocyte death is not driving the reduced pigmentation phenotype observed after ML233 treatment. This result also indicates that ML233 regulates skin pigmentation via melanogenesis inhibition rather than inducing developmental defects in melanocytes.

To assess whether ML233 can regulate melanin production directly in melanocytes, we tested its activity in highly pigmented embryos. We demonstrated that embryos treated with ML233 at 48 hpf exhibit reduced skin pigmentation at 54 and 72 hpf (Fig. S2A, B), after 6 and 24 hours of ML233 treatment, respectively. This result suggests a rapid and potent effect of ML233 on the regulation of melanin production and/or quantity by melanocytes. Together, these experiments support the hypothesis that ML233 affects the capacity of differentiated pigment cells to produce melanin. Therefore, we propose that ML233 can be classified as a small chemical inhibitor of melanogenesis *in vivo* with a reversible lightening effect on skin pigmentation.

### ML233 inhibits melanogenesis independently of the apelin signaling pathway

Because ML233 was originally described as a small chemical agonist of the apelin signaling pathway^42,43^, we wanted to determine whether ML233-mediated inhibition of melanogenesis is dependent of apelin-receptor activation in zebrafish. The zebrafish genome contains two apelin receptor genes that result from ancestral genome duplication^44^, *aplnra* and *aplnrb*, and also contains one *aplnr2* gene. To knock out (KO) these genes and assess their functions in the ML233-dependent effect on skin pigmentation, we designed and validated three gRNAs to target these loci (one gRNA for each gene) and induce indels using CRISPR/Cas-9. We first generated *aplnra*-*aplnrb* double-KO embryos by injecting the corresponding two gRNAs and the Cas-9 protein in one-cell-stage zebrafish eggs, and then treated the embryos with ML233 at 15 µM until 3 dpf. We did not notice any increase in pigmentation in the double-KO embryos, and the overall pigmentation of these embryos at 3 dpf is comparable to that of control embryos (Fig. S2C and Fig. S3A), suggesting that apelin-receptor-dependent pathways are not involved in homeostatic control of skin pigmentation. Genotyping and indel analysis of individual embryos injected with the *aplnr* gRNAs confirmed the introduction of a high percentage of indels in the *aplnra* and *aplnrb* loci and the absence of pigmentation defects in these embryos (Fig. S3B). Importantly, we also did not observe any rescue of the ML233-reduced skin-pigmentation phenotype with KO of the *apelin receptor a* and *b* genes (Fig. S2C and Fig. S3A). We also targeted the *aplnr2* gene alone or in combination with *aplnra* and *aplnrb* double-KO and demonstrated that none of these genetic conditions leads to rescue of skin pigmentation after ML233 treatment (Fig. S3C, D). Together, the above results suggest that the effect of ML233 on melanogenesis inhibition is independent of apelin receptor activity. Finally, to confirm the above results and demonstrate that the apelin signaling pathway is not involved in control of skin pigmentation and melanocyte function, we performed an apelin peptide gain- and loss-of-function analysis. We generated a transgenic line, Tg(*ubi:Lox-stop-Lox-apelin*), allowing temporal control of apelin peptide expression. Using this new transgenic line in combination with the previously described Tg(*ubi:creERT2*)^45^, we expressed apelin ubiquitously at 24 hpf by addition of hydroxy-tamoxifen (4-OHT) at 10 µM. We then monitored the pigmentation of these embryos during the next two days and observed no reduction in skin pigmentation compared to control embryos (Fig. S2D, top and middle panels). We also established a new apelin CRISPR/Cas-9 mutant harboring a 26-bp insertion that results in a premature stop codon at position 18 of the amino-acid sequence (Fig. S3E). Supporting the above observation regarding the loss of apelin receptor function, loss of apelin peptide function has no effect on skin pigmentation at 3 dpf (Fig. S2D, bottom panel). Together, our study of apelin receptor loss-of-function, apelin overexpression, and apelin mutant phenotypes strongly suggests that ML233-mediated inhibition of melanogenesis *in vivo* is independent of apelin-receptor activation and signal transduction.

### ML233 inhibits tyrosinase activity *in vivo* and *in vitro*

To assess whether ML233 treatment and the reduced skin-pigmentation phenotype correlates with TYR dysfunction, we determined the enzymatic activity of zebrafish tyrosinase under different conditions. We prepared cellular extracts containing TYR protein from control or ML233-treated zebrafish embryos and quantified the capacity of the protein to convert L-DOPA to melanin, with absorbance at 475nm. We observe significantly reduced L-DOPA conversion at 2 dpf after ML233 treatment for 24 hours compared to DMSO-only-treated embryos (Fig. 3A), suggesting inhibition of TYR protein activity. This experiment also demonstrates that ML233 concentrations as low as 0.5 µM have a robust capacity to inhibit TYR function by ∼60%. At a concentration of 10 µMm, ML233 reduces TYR enzymatic activity by ∼80%, an inhibition similar to a treatment with PTU at 200 µM (Fig. 3A). These observations correlate with the reduced pigmentation phenotype and the strong reduction of melanogenesis observed after ML233 treatment (Fig. 2A, B). Together, these results indicate that ML233 acts as a potent inhibitor of TYR enzymatic activity *in vivo*.

**Figure 3.**
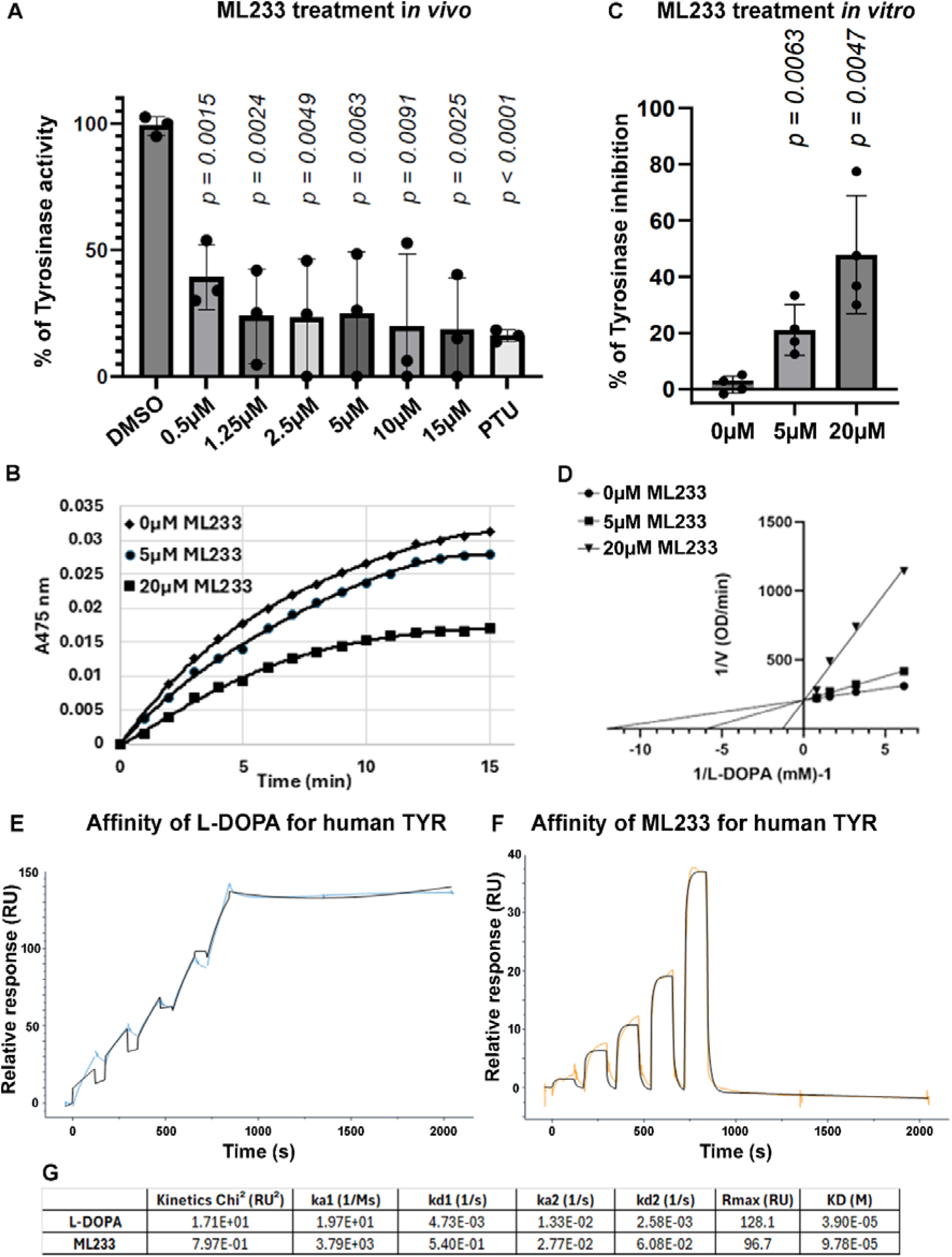
ML233 inhibits TYR protein activity *in vivo* and *in vitro*. **(A)** Tyrosinase activity was quantified in cellular extracts of DMSO- or ML233-treated embryos for 24 hours (n=3 biological replicates with 3 technical replicates; ≥40 embryos/biological replicate). Significance is determined by t-test, two-tailed, unpaired. Error bars represent s.d. **(B-D)** Kinetic assays of L-DOPA conversion by Tyrosinase in presence or absence of ML233 at 5 µM or 20 µM over a 15 min time course analysis (B). Tyrosinase activity was quantified *in vitro* in DMSO- or ML233-treated samples (n=4). Significance is determined by t-test, two-tailed, unpaired. Error bars represent s.d. (C) Lineweaver-Burk plot indicates that after 1 min, ML233 acts with a competitive inhibition mode on the Tyrosinase protein (D). **(E, F)** Two-state reaction model SPR analysis to determine the binding affinity of L-DOPA (C) and ML233 (D) with human TYR. Two-fold dilution series of 5 doses with a high concentration of 500 µM for L-DOPA and 50 µM for ML233 were used. **(G)** Table showing the constants of association and dissociation calculated for L-DOPA and ML233 interaction with human TYR. Significance is determined by t-test, two-tailed, unpaired. Error bars represent s.d.

We next aimed to determine *in vitro* whether this small inhibitor of melanogenesis acts directly on TYR activity. We assessed the activity of mushroom TYR, via measurement of its capacity to convert L-DOPA to melanin *in vitro*. Kinetic analysis over 15 min of competition between L-DOPA and ML233 to inhibit mushroom TYR suggests that ML233 reduces melanogenesis through a mechanism of competitive inhibition (Fig. 3B-D). Quantification of ML233-mediated melanogenesis inhibition after 1 min of the time course analysis revealed a significant inhibition of melanogenesis by ∼20% or ∼50% in samples treated with 5 µM or 20 µM of ML233 respectively compared to DMSO-only-treated samples (Fig. 3C). Finally, to determine the type of interaction between TYR and L-DOPA or ML233, we performed surface plasmon resonance (SPR) analysis with purified human recombinant TYR. We used a two-state reaction model with 5 different concentrations (2-fold serial dilution) with 500 µM and 50 µM as the highest concentrations for L-DOPA and ML233, respectively. SPR analysis suggests that L-DOPA and ML233 can bind human TYR with different modes of action (Fig. 3E-G), slow on/off for L-DOPA (ka1 [1/Ms] = 1.97e+1 and KD [M] = 3.90e+5) and fast on/off for ML233 (ka1 [1/Ms] = 3.79e+3 and KD [M] = 9.78+5,) respectively, indicating that ML233 is quicker to bind human TYR but that L-DOPA is more difficult to dissociate. Together, these *in vitro* experiments confirm our *in vivo* observations regarding ML233-mediated inhibition of TYR activity and suggest that this small chemical inhibitor of melanogenesis acts directly on the TYR protein to regulate its function.

### Tyrosinase gene expression is not abolished by ML233 treatment

To better understand the molecular action of ML233 on skin pigmentation and melanogenesis, we analyzed the expression of genes involved in melanogenesis, including tyrosinase (*tyr*), dopachrome tautomerase (*dct*), and microphthalmia-associated transcription factor (*mitfa*), the latter which is known to control melanocyte formation and TYR expression. Global quantification of *tyr*, *dct*, and *mitfa* mRNA expression by RT-qPCR in zebrafish embryos at 2 dpf indicates a mild (∼15% at 1.25-5 µM) to moderate (∼40% at 5-20 µM) reduction of gene expression after 1 day of ML233 treatment (Fig. 4A). Significantly decreased *tyr* expression is observed at ML233 concentrations of 5 µM and above, with a maximum reduction of ∼40% at 20 µM of ML2333 compared to control DMSO-treated embryos. Expression of the transcription factor *mitfa*, master regulator of melanocyte formation and melanogenesis, is not significantly dysregulated by ML233 treatment at any concentration. In addition, we showed by *in situ* hybridization that the *mitfa* and *tyr* genes are still expressed in melanocytes at 2 dpf after 1 day of 15 µM ML233 treatment (Fig. 4B). Similarly to PTU-treated embryos, *tyr* mRNA is highly detectable in ML233-treated embryos in a pattern resembling that of melanocyte skin cell organization. These results support our above observations indicating that melanocytes are still present in the skin after ML233 treatment despite the lack of melanin production. Based on these gene-expression results, we hypothesized that while reduced expression of *tyr* and/or *mitfa* mRNA could partially underlie the reduced skin-pigmentation phenotype, other factors likely contribute to the inhibition of skin pigmentation observed after ML233 treatment.

**Figure 4.**
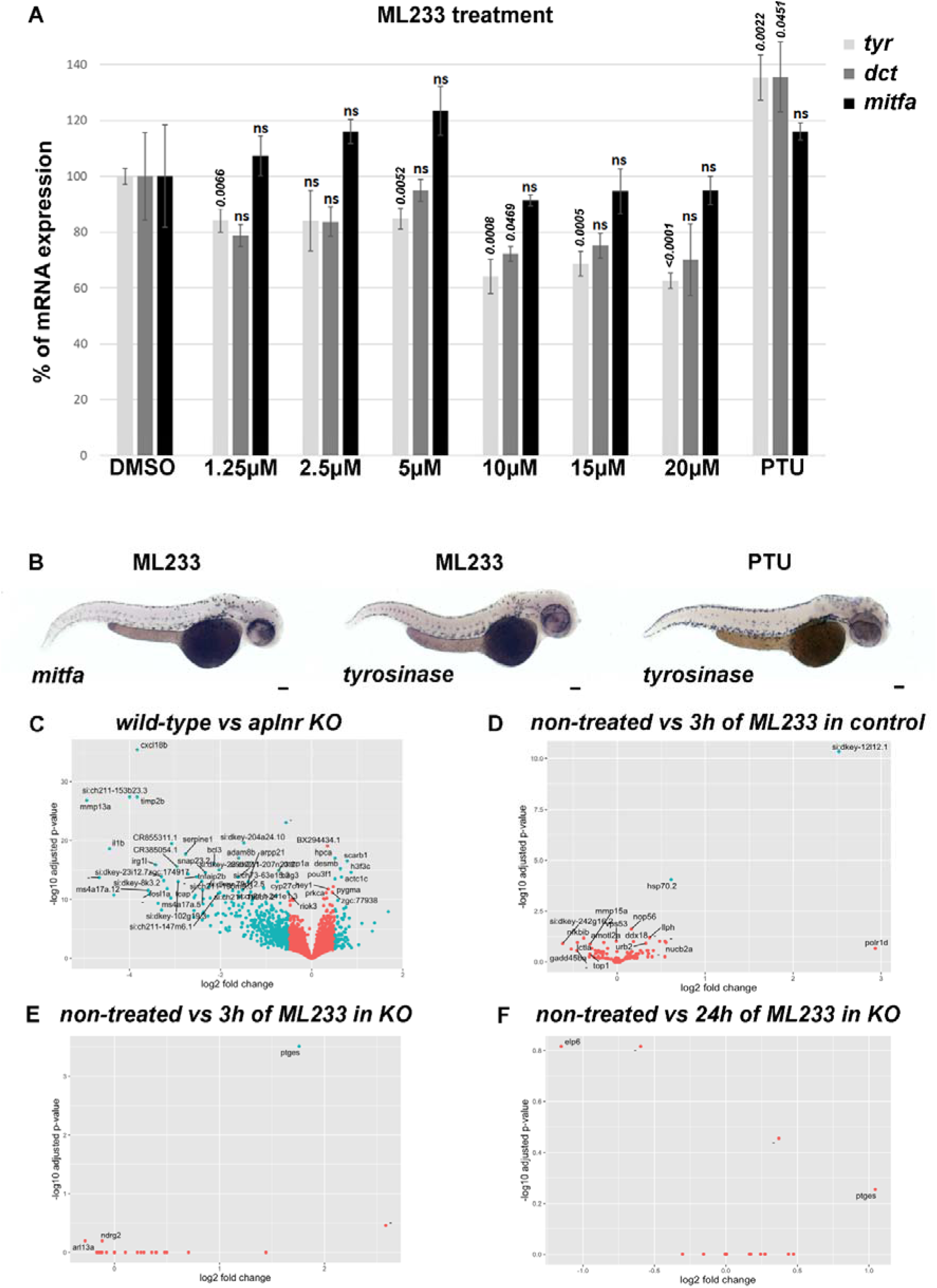
ML233-dependent regulation of TYR function is not mediated at the transcriptional level. **(A)** Analysis of *tyr*, *dct*, and *mitfa* mRNA expression by RT-qPCR in DMSO-, ML233-, or PTU-treated (between 24 and 48 hpf) zebrafish embryos at 48 hpf (n=3 biological replicates with 3 technical replicates; ≥40 embryos/biological replicate). Significance is determined by t-test, two-tailed, unpaired. **(B)** Analysis of *mitfa* and *tyr* mRNA expression by *in situ* hybridization in ML233- or PTU-treated (between 24 and 48 hpf) embryos at 48 hpf. **(C-F)** Graphical representation of bulk RNAseq analysis (n=4; ≥20 embryos/sample) and variation in gene-expression profiles in WT vs. apelin-receptor KO, both untreated (C), WT after 3 hours of ML233 treatment (D); apelin-receptor KO after 3 hours of ML233 treatment (E); and apelin-receptor KO after 24 hours of ML233 treatment (F). Scale bars: 100 μm. Error bars represent s.d.

Because ML233 is an agonist of the apelin receptors^42,43^, we decided to investigate ML233-driven differential expression in an apelin-dependent and - independent manner. We treated zebrafish embryos with 0.5 µM of ML233 (a concentration showed to significantly reduce melanin content and TYR protein activity, Fig. 2B and Fig. 3A) for 3 hours or 24 hours in a wild-type (WT) or apelin-receptor KO genetic background. We observed a clear separation in gene expression between the WT and KO backgrounds, both untreated (Fig 4C). However, minimal expression changes were identified for either the WT or KO genetic background treated with ML233 (Fig. 4D-F), indicating that genotype differences (WT vs KO) are the strongest determinant underlying change in gene-expression profiles. Gene Set Enrichment Analysis (GSEA) indicated that ML233 treatment at this concentration has no significant impact on biological processes such as, but not limited to, inflammation, stress response, or apoptosis (Table S1 and Table S2). GO and KEGG analyses of ML233 impact in the KO also support the above observation (Table S3-6), further indicating ML233 as a potentially tolerable pharmaceutical chemical. Altogether, these data indicate that the ML233 molecule has little impact on the transcriptome profile at a concentration sufficient to inhibit pigmentation. This analysis also supports the above hypothesis suggesting that dysregulation of *tyr* gene expression is not the main mechanism leading to reduced skin pigmentation after ML233 treatment, as its expression is not significantly affected in the transcriptomic analysis.

### TYR protein expression is not affected by ML233 treatment

Because *tyrosinase* mRNA expression is not dramatically reduced after ML233 treatment *in vivo* and because *in vitro* data suggest that ML233 can directly inhibit TYR protein function, we hypothesized that a reduction in skin pigmentation could also reflect either a reduction of TYR protein expression and/or inhibition of TYR protein enzymatic activity. To study these two possibilities, we used an antibody raised against the human TYR protein. We tested the capacity of the antibody to recognize TYR in two mammalian cell lines (Fig. 5A and Fig. S4A, C): B16F10 (murine melanoma) and A375 (human melanoma). Using the validated antibody, we analyzed the level of tyrosinase protein expression by western blot in control DMSO- and 5 µM ML233-treated cells (Fig. 5B and Fig. S4B, D, E). We observed no significant reduction of TYR protein expression normalized with beta-actin protein expression (Fig. 5C and Fig. S4E). Importantly, in the same cells and at the same concentration of ML233 treatment, we observed a strong reduction in pigmentation (Fig. 5D). This observation indicates that inhibition of melanin production by ML233 is not dependent on TYR protein degradation, suggesting that a different molecular mechanism underlies the phenotype.

**Figure 5.**
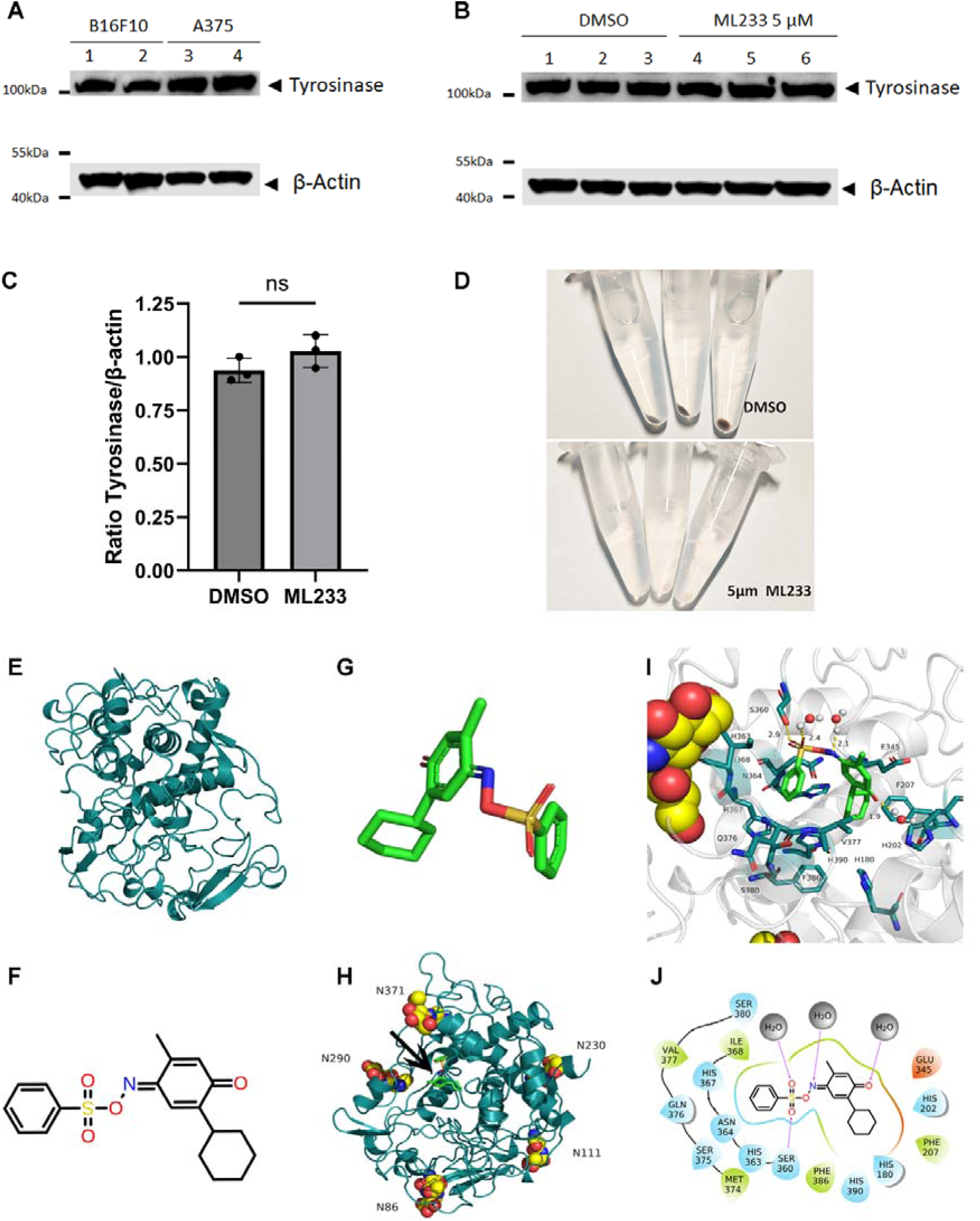
ML233-dependent regulation of TYR function is not mediated through protein degradation and is computationally predicted to take place via direct inhibition of the TYR active site. **(A)** Expression of tyrosinase protein was analyzed by western blot in murine (B16F10) or human (A375) melanoma cells (n=3). **(B)** Expression of tyrosinase protein was analyzed by western blot in murine (B16F10) melanoma cells after DMSO or ML233 treatment (n=3). **(C)** Tyrosinase protein expression was quantified and normalized by expression of the beta-actin protein after DMSO or ML233 treatment. Significance is determined by t-test, two-tailed, unpaired. Error bars represent s.d. **(D)** Representative pictures of melanin expression in murine (B16F10) melanoma cells after DMSO or ML233 treatment. **(E)** 3D structure of human TYR protein. **(F)** 2D representative structure of the ML233 chemical. **(G)** 3D representative structure of the ML233 chemical. **(H)** 3D structure of human TYR protein with asparagine N-glycosylation sites and ML233 binding site (black arrow). **(I)** 3D representation of TYR and ML233 interaction. **(J)** 2D representation of TYR and ML233 interaction. Hydrogen bonds are represented by purple arrows, and amino-acid colors indicate different properties: green for hydrophobic amino acids, cyan for polar amino acids, and red for acidic negatively charged amino acids. Water molecules are represented by gray spheres.

### Computational prediction of ML233-dependent direct inhibition of tyrosinase protein function

We next studied the structural relationship between ML233 and the TYR protein to better understand the potential mechanisms underlying ML233-dependent inhibition of TYR activity. The protein structure of human TYR was predicted by AlphaFold2 (Fig. 5E). The two-dimensional (2D) structure of ML233 is shown in Figure 5F, and the derivative 3D structure used to establish the predictive conformation is presented in Figure 5G. Five potential asparagine N-glycosylation sites were identified in TYR (Fig. 5H): N86, N111, N230, N290, and N371. Interestingly, both N86 and N371 were previously described to be important for full enzymatic activity of TYR^46^. Finally, molecular docking experiments were conducted to predict the optimal binding site of ML233 within the TYR protein (Fig. 5G-J). This analysis indicates that ML233 could bind the ligand-binding pocket of the human TYR protein.

We also investigated the molecular dynamics and binding energy between the TYR protein and the small chemical inhibitor. Dynamic simulation showed that the overall protein structure is relatively stable and that the binding site of the small molecule did not change during the simulation (Fig. S5A and Video S1). We also determined whether the hydrophobic and hydrophilic areas changed following ML233 binding during the binding simulation. The simulation shows only a subtle change in the distribution of these two areas (Fig. S5B), indicating that the small molecule binding has a small effect on the hydrophilic and hydrophobic domains. We also quantified ML233-TYR binding stability, using Root Mean Square Deviation (RMSD) analysis (Fig. S5C). The RMSD value remained lower than 2 Å throughout the simulation, maintaining an average level of 1.54 Å. An RMSD value below 2 Å suggests no drastic structural change, predicting that the TYR-ML233 interaction and the protein structure are relatively stable. We determined that the amino acids showing the greatest amount of time of contact with ML233 during the dynamic simulation are His180, Glu345, Ser360, His367, Ile368, and Val377 (Fig. S5D), suggesting that these are the most important amino acids for binding of ML233 to the TYR protein.

The binding free energy between ML233 and TYR was also analyzed (Fig. S5E), revealing a binding free energy of −9.87kcal/mol and a dissociation constant of 58.18 nmol, the latter of which was calculated as Kd = ΔG/RT. This binding affinity is primarily the result of the van der Waals action (ΔE_vdw_), which is −34.13kcal/mol, together with the almost negligible electrostatic effect (ΔE_ele_), which is only - 1.62kcal/mol. The free energy of polar solvents (ΔE_GB_) is 6.14 kcal/mol, and the free energy of non-polar solvents (ΔE_surf_) is −4.59 kcal/mol, indicating that the total free energy (ΔGsolv) is 1.55kcal/mol less energy than would be lost in desolvation. Because we observed no drastic change in protein structure during the dynamic simulation, we hypothesized that the potential entropy loss (-TΔS=24.33kcal/mol) of the system is due to subtle structural adjustments and oscillations of the small molecule itself at the binding site.

A potential mode of interaction between ML233 and the TYR protein is represented in Figure 5J. The sulfoxide group of the ML233 molecule includes oxygen atoms that can form a 2.9Å hydrogen bond with Ser360. Another oxygen atom in the sulfide group and the unsaturated N atom form 2.4Å and 2.1Å hydrogen bonds, respectively, with water molecules. The carbonyl oxygen atom on the molecular benzene ring also forms a 1.9Å hydrogen bond with a water molecule. In addition, the benzene ring of the molecule forms hydrophobic interactions with Ile368 and Val377, and forms polar van der Waals interactions with His367, Ser380, Gln376, and other polar amino acids. Finally, the cyclohexane molecule has hydrophobic interactions with the hydrophobic amino acids Phe207 and Phe386, and van der Waals contacts with the polar amino acids His180, His202, and His390.

Histidines His180, His202, His367, and His390 on the human TYR protein have been proposed to regulate TYR activity by binding copper ions^47–49^. The above computational analysis suggests that a direct interaction between ML233 and TYR in the functional site of the protein could occupy copper-ion binding sites and potentially inhibit TYR enzymatic activity. This predicted mechanism of inhibition supports *in vivo* and *in vitro* data regarding the ML233-mediated inhibition of TYR activity.

### ML233 reduces pigmentation in a mammalian melanoma cell line

Our *in vivo*, *in vitro*, and computational studies above support our hypothesis that ML233 is a direct inhibitor of TYR protein expression and function. To determine how this effect is conserved in higher vertebrate species, we tested ML233 activity in the B16F10 murine melanoma cell line. We first analyzed B16F10-cell proliferation after ML233 treatment to determine the IC50. We observed a 50% reduction in the number of cells at concentrations of ML233 between 5 and 10 µM (Fig. 6A), using ATP production as an indirect readout of cell viability. Comparison of the effect of ML233 with that of cisplatin treatment (Fig. 6A, B) suggests a potent inhibition of melanoma-cell proliferation by ML233. We next asked whether ML233 can inhibit pigmentation of B16F20 cells at low ML233 concentrations (without effects on cell proliferation). We quantified melanin production in B16F10 cells after 24 hours of treatment with 0.625 µM, 1.25 µM, 2.5 µM, or 5 µM of ML233. Observation of cell pellets after centrifugation confirms that the ability of ML233 to inhibit melanin production is conserved in mammals (Fig. 6C). Finally, absorbance quantification of melanin confirmed this observation, and revealed that concentrations of ML233 as low as 0.625 µM significantly reduce melanoma-cell pigmentation (Fig. 6D, E). Importantly, concentrations of ML233 between 0.625 and 5 µM have the capacity to reduce melanin production without affecting B16F10-cell survival. Together, these results confirm our *in vivo* data and validate ML233 as a potent inhibitor of melanogenesis in mammalian cells.

**Figure 6.**
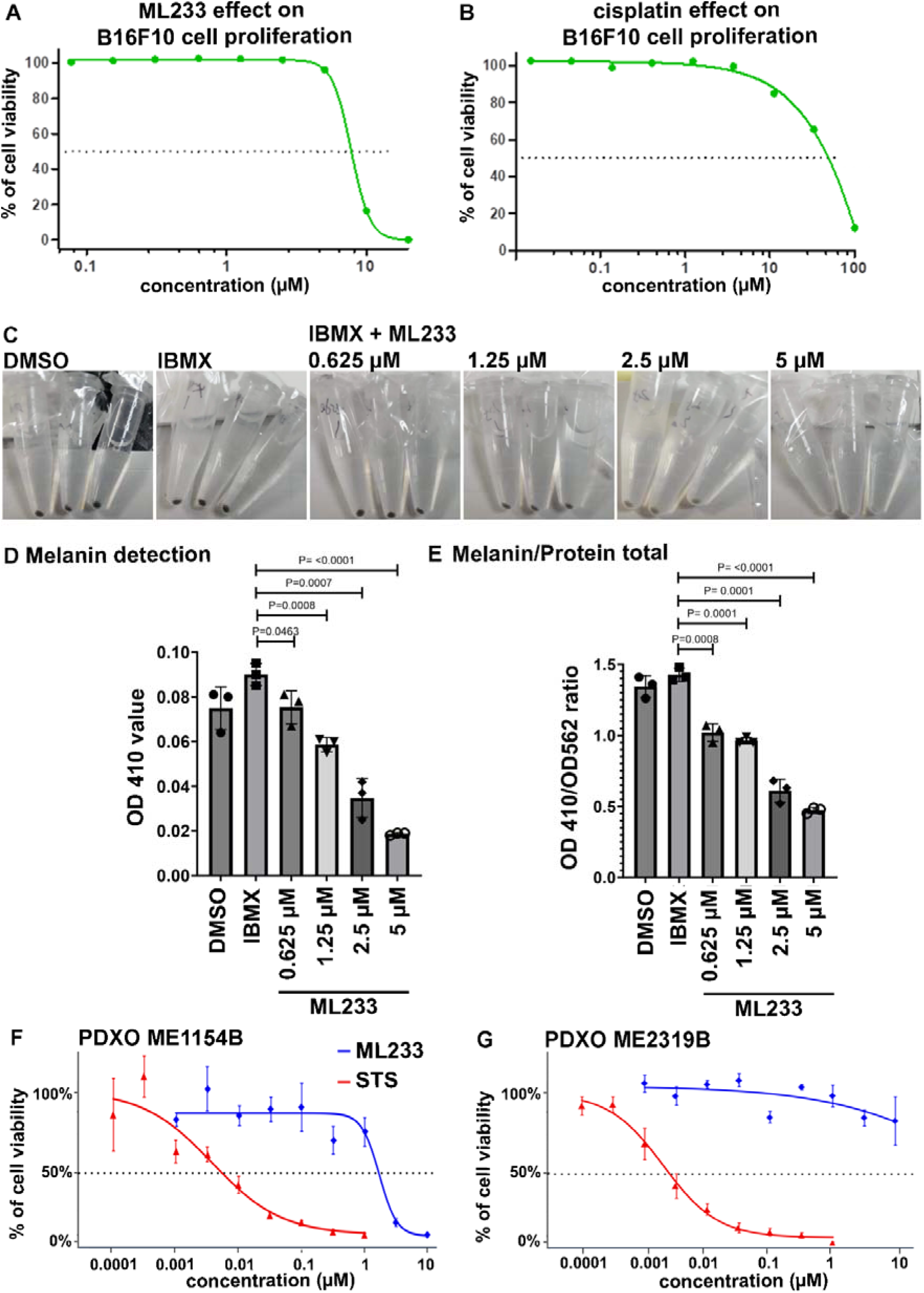
ML233 inhibits melanogenesis and melanoma proliferation in mammals. (A, **B)** Effect of ML233 (A) or cisplatin (B, positive control) treatment on B16F10 murine melanoma cell proliferation (n=3). **(C)** Representative pictures of melanin expression in murine (B16F10) melanoma cells after treatment with DMSO, IBMX alone, or IBMX plus ML233. **(D, E)** Melanin expression was quantified (D) and normalized (E) after treatment with DMSO, IBMX alone, or IBMX plus ML233 (n=3). Significance is determined by t-test, two-tailed, unpaired. (**F, G)** Cell proliferation in PDXO cellular models was quantified in ME1154B (F) or ME2319B (G) human melanoma lines after staurosporine (STS, positive control) or ML233 treatment (n=3). Error bars represent s.d.

### ML233 inhibits proliferation of PDX-derived melanoma organoids

Because ML233 affects B16F10 murine melanoma-cell proliferation, we next asked whether it could also reduce the proliferative potential of human metastatic melanoma. We used patient-derived xenograft organoids (PDXOs), which are 3D *in vitro* models generated from patient tumors. We tested the effect of ML233 treatment (between 0.001 and 10 µM) in two human melanoma cell lines collected from metastasis: ME1154B and ME2319B (Crown Bioscience). Interestingly, we observed that these two melanoma lines differ in their sensitivity to both ML233 and the control molecule, staurosporine (STS), in 3D organoids (Fig. 6F, G). ML233 inhibited ME1154B viability/proliferation (IC50 = 1.65 µM) but did not affect ME2319B viability at any concentration tested. However, ME2319B is more sensitive than ME1154B to STS treatment (IC50 = 0.0025 µM; IC50 = 0.0054 µM, respectively). Together, these data indicate that ML233 has the potential to inhibit proliferation of human metastatic melanoma in PDXO *in vitro* models, and that this capacity is dependent on the particular melanoma subtype.

## DISCUSSION

Inhibition of TYR activity is a potent way to reduce melanogenesis and skin pigmentation. The use of tyrosinase inhibitors as skin-lightening agents is of great interest for treatment of pigmentation disorders such as vitiligo syndrome (to lighten the color of healthy skin so that it matches the color of the disease-impacted skin), and hypo- or hyper-pigmentation-associated diseases. In this study, we identified an effective skin-lightening agent, ML233. We demonstrated that ML233 reduces skin pigmentation without affecting melanocyte development and without significant toxicity in zebrafish embryos, and that this effect is reversible. Our SPR and molecular docking studies indicate a potential direct interaction and competitive inhibition between ML233 and the human TYR protein activity. While these *in vitro* and *in silico* approaches yield different outputs regarding the affinity and stability of the ML233-human TYR interaction, they both support the chemical-protein interaction. The western blot analysis indicates that TYR protein expression is not affected by ML233 treatment at a dose of 5 µM, a dose that leads to inhibition of melanogenesis in B16F10 cells. However, our results also show that increased concentrations of ML233 have a moderate but statistically significant impact on *tyr* gene expression in zebrafish. Further study will be needed to determine whether this effect results from ML233-mediated regulation of *tyr* mRNA expression or a regulatory control between TYR protein activity and its own gene expression. Our gene and protein expression analysis suggests that ML233 regulates TYR function primarily independently of the control of mRNA and protein expression; understanding the effect of different doses of ML233 on gene regulation and protein expression and function will be an important future area of investigation. Together, our combination of *in vivo*, *in vitro*, and *in silico* analysis suggests that ML233 inhibits TYR activity at the protein level via a direct interaction, possibly with the TYR active site, preventing copper ions from binding and activating TYR. ML233 and efficient derivatives could be new attractive compounds in the hunt to alleviate discomfort associated with skin-pigmentation disorders.

For over two decades, melanogenesis inhibitors have been at the center of therapeutic research to treat skin pigmentation disorders. Many such inhibitors have been tested, including hydroquinone, kojic acid, azelaic acid, and arbutin ^18,31–34^. Unfortunately, despite their relative efficiency, none have been widely accepted to treat skin pigmentation^35–37^, mainly because of safety issues regarding an increased risk of developing skin cancer after long-term use. Supporting the concern regarding use of currently available TYR inhibitor agents is the recent report of a 55-year-old woman showing potentially irreversible skin damage (exogenous ochronosis) after use of a hydroquinone-containing skin-lightening cream to treat melasma^50^. Current efforts to treat skin-pigmentation disorders such as vitiligo focus on use of the recently FDA-approved JAK1/2 inhibitors to antagonize activation of the JAK/STAT signaling pathway^51–54^. This strategy limits the death of melanocytes by apoptosis, leading to a variable range of skin re-pigmentation in treated patients. The combination of JAK1/2 inhibitors with phototherapy has been suggested to improve the efficiency of re-pigmentation^51^, but the study was limited to two patients. The long-term effects of JAK1/2 inhibitors are currently not well appreciated because they have been developed and tested only recently. While these inhibitors are considered relatively safe for human health and are quite effective in skin lightening, recent studies suggest that they could underlie undesirable side effects including, but not limited to, infection, embolism, and thrombosis^53–57^. Also, some patients cease to respond to JAK inhibitors after a length of time of treatment, although this outcome differs among individuals. Together, the above information highlights the need for alternative, safe, and effective skin-lightening agents to treat, for example, resistant vitiligo not responding to JAK/STAT inhibitors, and hyperpigmentation disorders such as post-inflammatory pigmentation or melasma^57^.

Importantly, our *in vitro* studies revealed significantly reduced proliferation of B16F10 and PDXO melanoma lines after ML233 treatment. This observation supports the hypothesis that ML233 could be used as a new potent small chemical that couples inhibition of both cell proliferation and melanogenesis in melanoma cells. However, we found that one of the two PDXO melanoma cell lines tested did not respond to ML233 treatment, but the factors contributing to this resistance are not known as the biological mechanisms underlying the ML233-dependent effect on proliferation are not understood. Because the dose of ML233 required to reduce pigmentation is lower than that required to inhibit melanoma-cell proliferation, we hypothesized that reduced cell proliferation is independent of the regulation of melanogenesis. Testing this hypothesis is a potentially valuable future avenue for better understanding the mechanisms through which ML233 reduces melanoma-cell proliferation. For melanoma-cancer subtypes that respond poorly to current treatment strategies, the use of ML233 in synergy with current widely used therapeutic strategies, such as immuno- and targeted-therapy, could potentially increase treatment efficacy. Future work will shed light on the efficiency and safety of ML233 as a therapeutic agent to reduce melanoma-cell proliferation.

While our work establishes a safer toxicity profile for ML233 compared to the widely used kojic acid and arbutin, it is necessary to assess the potential side effects of ML233 in mammalian models. Systemic delivery of ML233 could potentially impact pigmentation in other organs such as the retina and kidney or reduce neuromelanin expression in the central nervous system. Future pre-clinical studies in mammals will enable identification of the potential side effects of topical and systemic use of ML233 for therapeutic purposes. Demonstration that uses of ML233 can produce the desired outcomes without affecting other organs will be an important milestone.

As discussed above, ML233 is also an apelin-receptor agonist. Here we have demonstrated that reduced melanogenesis after ML233 treatment is independent of apelin-receptor function. We also showed that most of the gene-expression variation observed after ML233 treatment in wild-type embryos is not observed in apelin-receptor KO embryos post-ML233 treatment, suggesting that ML233 regulates skin pigmentation in an apelin-receptor-independent manner by interacting with the TYR protein. These results were corroborated by our analysis of mutant zebrafish with KO of the *aplnra*, *aplnrb*, and *aplnr2* genes indicating that none of the knockouts led to skin-pigmentation rescue after ML233 treatment, and by the Daniocell single-cell database^58^, which shows that expression of these genes is not detected in melanocytes. This notion is also supported by our *in vitro* study of TYR activity after ML233 treatment, our SPR analysis, and the molecular docking prediction. Because the PDXO melanoma lines we tested do not express APJ (apelin receptor in mammals), and because low doses of ML233 in B16F10 cells inhibit melanogenesis without affecting cell proliferation, we hypothesized that reduced melanoma proliferation following ML233 treatment is not dependent on the apelin signaling pathway or TYR inhibition. However, the molecular mechanism underlying this effect still needs to be identified. Together, our data suggest that ML233 can regulate melanogenesis and melanoma-cell proliferation without apelin-receptor activation. Future studies are needed to determine the capacity of the ML233 molecule to inhibit TYR protein activity and/or melanoma-cell proliferation but not activate the apelin pathway. Such molecules could have significant potential for cosmetic or therapeutic applications.

## ONLINE METHODS

### Study design and statistical analysis

Research subjects (embryos, cell cultures, organoids…) and treatments applied are defined in the figures and/or figure legends. Experimental analyses were performed with a minimum of 3 biological replicates and 3 technical replicates for melanin quantification *in vivo*, tyrosinase activity *in vivo*, and gene expression analysis by qPCR (with a minimum of 20 zebrafish embryos per biological replicate). Regarding statistical power, we calculated the effect size with Cohen’s D for both melanin quantification and tyrosinase activity measurements and found that all measured differences had D-values that exceeded an absolute value of 4, which qualifies as a very large effect. For experiments using zebrafish embryos as one biological replicate such as survival and hatching analysis, a minimum of 20 embryos was used. The number of biological replicates and embryos used for each specific experiment as well as the statistical test used to calculate significance is indicated in the figure legends. For statistical analysis, p-values are indicated in the figures. Comparisons were conducted to determine significant differences between control and experimental conditions. The significance threshold was established at 5%. Processing and method analysis of the bulk RNAseq dataset is described below in the Methods section.

### Fish lines and developmental conditions

Embryos were raised and staged according to standard protocols^59^. Experiments were performed in accordance with animal care guidelines. Our IACUC animal protocol (AUP#22-06) is approved by the MDI Biological Laboratory. Tg(*ubi:creERT2*)^45^ was used to express the apelin peptide ubiquitously. For imaging analysis, embryos were fixed overnight at 4°C in 4% paraformaldehyde/1xPBS, after which they were dehydrated through an ethanol series and stored at −20°C until use.

### CRISPR/Cas-9 generation of zebrafish mutants

To generate *tyrosinase*, *apelin*, *apelin receptor a*, and *apelin receptor b* mutants, we took advantage of a previously described genome-editing method^60^. Specifically, we used tracrRNA and crRNA (IDT) to form functional gRNA duplexes targeting the ORF (*apln*: 5’-GAATGTGAAGATCTTGACGC-3’, *aplnra*: 5’-TGGGTGTGACTACTCGGAGT-3’, *aplnrb*: 5’-CTTGCAGAGTGCCACGCCAA-3’ and *aplnr2* 5’-GTGGCAACGGTTCAAAGCAG-3’). Co-injection of these gRNAs with the Cas-9 protein (IDT) resulted in indels in the *apln*, *aplnra*, *aplnrb*, or *aplnr2* gene coding sequences, identified by Sanger sequencing and analyzed with ICE software (Synthego) in F0 injected embryos. F0 carriers for mutations in the *apelin* locus in the germline were identified by sequencing the alleles on the clutches, and were then outcrossed to obtain F1 heterozygous mutants. Mutations (indels) leading to a premature stop codon were identified by ICE analysis (Synthego) of the Sanger-sequencing results. F2 *apelin* homozygous mutants were also identified by sequencing. The *apelin* mutant harbor a 26-bp insertion leading to a stop codon at position 18 of the peptide.

### Plasmid construction and transgenic line establishment

For generation of Tg(*ubi:Lox-stop-Lox-apelin*) zebrafish, a p5E entry vector containing the ubiquitin promoter and the LoxP cassette^45^ was used. The *apelin* ORF was amplified by PCR from zebrafish cDNA and cloned in the pME middle entry vector. The appropriate entry and middle entry clones were mixed with the SV40pA 3’ entry vector and recombined into the Tol2 transposon destination vector. To establish stable transgenic lines, plasmids were injected into one-cell-stage embryos with Tol2 transposase mRNA^61^.

### Pharmacological treatments

Stock solution of 1-phenyl 2-thiourea (PTU; Sigma) was prepared at 25X (0.075%). PTU 1X working solution was used to inhibit pigmentation in zebrafish embryos. Stock solutions of ML233 (Tocris), arbutin (Sigma), and kojic acid (Sigma) were prepared at 25 mM in 100% DMSO (Sigma). Working concentrations and times of treatments are specified in the Results section. A stock solution of hydroxytamoxifen (4-OHT) was prepared at 10 mM in 100% DMSO. For induction of apelin expression, zebrafish embryos were treated for 24 hours at a concentration of 10 µM.

### Acute toxicity studies

Preliminary acute toxicity tests were performed to determine the values of the lethal concentration (LC50). Twelve concentrations ranging from 0 µM to 200 µM were selected using a spacing factor equal or inferior to 2.5. Briefly, embryos were distributed in 6-well plates with 40 embryos per well filled with 5 mL of E3 medium containing or not containing ML233. Subsequently, observation of sub-lethal and lethal morphological endpoints was performed at 1 and 2 dpf under a Zeiss Axio Zoom V16 (16x zoom with a high numerical aperture of NA 0.25). Dead embryos were counted and discarded every 24h and the treatment medium was also changed daily. We noticed the presence of precipitation in the medium starting from 20 µM, which suggested that the maximum solubility was reached.

OECD236 toxicity tests of zebrafish embryos were conducted according to OECD guidelines for the testing of chemicals^62^. Zebrafish embryos were exposed to ML233 up to 4 dpf. Range of concentrations tested were from 0 µM to 20 µM with a spacing factor equal to 2. Briefly, embryos were distributed in 24-well plates with 1 embryo per well filled with 2 mL of E3 medium containing ML233 (n=20/condition). Subsequently, observation of sub-lethal and lethal morphological endpoints was performed at 1, 2, 3 and 4 dpf under a microscope (Zeiss Axio Zoom V16). Dead embryos were counted and discarded every 24h.

The effect of half-maximal inhibitory concentration (IC50) of ML233 on cell proliferation was determined in B16F10 murine melanoma cells. Cells were seeded in 96-well microplates and exposed to the experimental compound (ML233 or cisplatin control treatment) for 3 days. Drug-induced effects on the number of cells were quantified by measuring ATP using CellTiter-Glo® Luminescent Cell Viability Assay (Promega), which indirectly indicates the presence of proliferative and/or metabolically active cells.

### Gene-expression analysis

*In situ* hybridizations were performed as previously described^63^. The ORFs of *tyrosinase* (*tyr*) and *mitfa* were cloned in a pCS2+ vector using zebrafish cDNA. Antisense DIG-labelled probes were transcribed using T3 polymerase (Promega) and the linearized pCS2+ plasmid containing the ORF as a template. DIG probes were detected using anti-DIG antibody (Roche), and *in situs* were revealed using BCIP reagent (Roche).

For qPCR experiments, embryos were homogenized in Trizol solution (Invitrogen) and total RNA was purified using chloroform (Fischer) and then isopropanol (Fischer). RNA (1 μg) was reverse-transcribed using an oligo-dT primer and RevertAid kit (Thermo-Scientific). The resulting cDNA was used for qPCR experiments with the PowerTrack SYBR Green qPCR Master mix (Applied Biosystems). Gene-expression levels were normalized to expression of the housekeeping gene *Beta-actin*.

Total RNA samples for bulkRNAseq analysis were prepared with ∼50 embryos/sample. Each experimental condition was performed using biological quadruplicates to ensure reproducibility. Trizol (Invitrogen) was used to homogenize cellular extracts with the TissueLyzer (Qiagen). RNA purification was performed as described above. For bulkRNAseq analysis, library preparation and sequencing were carried out by NovoGene using the Illumina NGS platform. Raw FASTQ sequencing data underwent preprocessing to filter out adapters, poly-N tails, and low-quality reads. Indexing and alignment were performed using HISAT2 (v2.0.5)^64^ against the GRCz11 (Danio rerio) reference genome obtained from Ensembl. Mapped reads were assembled using StringTie (v1.3.3b)^65^, and quantification was performed using FeatureCounts (v1.5.0-p3)^66^. Differential expression analysis was conducted in the R environment using DeSeq2^67^, and Gene Set Enrichment Analysis (GSEA) was performed using clusterProfiler^68^. All visualizations were generated using ggplot2. RNAseq data are available via the GEO database and the accession number GSE268076.

### Western blot analysis

Lysate was prepared with 200uL Ripa lysis buffer (Beyotime) with proteinase inhibitor and phosphatase inhibitor (Roche) for 30min on ice, and collected by centrifugation at 17,000g for 15min at 4°C. The supernatant was transferred to another tube to test the protein concentration with BCA (Thermo Fisher). Protein electrophoresis was performed with 20μg/sample followed by semidry transfer using Trans-Blot® Turbo™ Transfer System (Biorad) onto a PVDF membrane (Biorad). Blocking with TBST solution containing 5% skimmed milk for 1,5h. Primary antibody was incubated with β-actin antibody (CST #4967) and tyrosinase monoclonal antibody T311 (Thermo Fischer# MA5-14177) at 4°C overnight, and the membrane was washed at room temperature with TBST, 10min per wash. Secondary antibody was incubated with anti-rabbit IgG, HRP-linked antibody (CST#7074), and anti-Mouse IgG H&L (bs-40296G-IRDye8) at room temperature for 1h, and the membrane was washed with TBST on a shaker at room temperature, 3 times, 10min each. ECL (Bio-Rad) fluorescent luminescence images were captured by Image Lab software using HRP-ECL luminescence.

### Melanin content quantification

For determination of melanin content *in vivo*, approximately 100 zebrafish embryos were sonicated in cold lysis buffer (20 mM sodium phosphate (pH 6.8), 1% Triton X-100, 1 mM PMSF, 1 mM EDTA) as previously described^69^. For the *in vitro* study, melanin content was analyzed in B16F10 murine melanoma cells. Cells were seeded in 6-well microplates and treated for 24 hours before lysate preparation. An aliquot of the lysate was used to determine the protein content with a Bradford assay kit (Thermo-Scientific). The lysate was then clarified by centrifugation and the melanin precipitate was resuspended with 1 mL of 1 N NaOH/20% DMSO at 95°C for 1 h. Spectrophotometric absorbance of intracellular melanin content was measured at 410 or 490 nm with SpectraMax ID3 (Molecular Devices). Absorbance quantification was normalized to the total protein content.

### Tyrosinase activity quantification

Evaluation of the effect of ML233 on tyrosinase activity in the zebrafish model system was performed according to the published method^69^ with minor modifications. Briefly, about 40 embryos was treated from 24h to 48h with ML233 and lysed by sonication (40 s total sonication [10s on; 30s off; 50% amplitude]) in 500µl of lysis buffer 20 mM sodium phosphate (pH 6.8), 1% Triton X-100, 1 mM PMSF, 1 mM EDTA) containing protease inhibitors cocktail (Roche). After quantification, 250 µg of total lysate protein was added to a reaction mixture containing 50 mM phosphate buffer (pH 6.8) and 2.5 mM L-DOPA (Sigma). After incubation at 25°C for 60 min, dopachrome formation was measured at 475 nm using the microplate reader SpectraMax iD3. Tyrosinase activity was expressed as the percentage change in comparison to that in the control group, which was considered as 100%. All conditions were tested in triplicate.

Investigation of the anti-tyrosinase activity of ML233 against mushroom tyrosinase (Sigma) was performed according to the published method^70^ with slight modifications. Each microplate well contained reaction mixture comprised of 80 μL of ML233 sample diluted in phosphate buffer (50 mM, pH 6.8), 40 μL of 2.5 mM L-DOPA (1 mg in 2 ml phosphate buffer 50 mM pH 6.8) and 40 μL mushroom tyrosinase (90 U/mL). For kinetic analysis, samples were incubated for 15s and the absorbance at 475 nm of each well was measured every minute at 25°C for 15 min, using the microplate reader SpectraMax iD3. The DMSO concentration in all wells was always equal to 0.16%. All the conditions were tested in quadruplicate. The percentage of TYR inhibition was calculated after 1 min by the following equation: Tyrosinase inhibition (%) = ((B – S)/B) × 100, where B is for the blank, and S is for the absorbance of sample. The characteristic of tyrosinase inhibition was represented at 1 min of the kinetic assay by Lineweaver−Burk double-reciprocal plot at specified concentrations of the substrates and inhibitors.

### SPR analysis

SPR studies were performed by BioDuro-Sundia using a Biacore 8K (Cytiva) with Series S Sensor Chip CM5 (Cytiva). The assays were performed at a temperature of 25°C. The target protein was human recombinant tyrosinase at 30 µg/ml (Origene), and the ligands were L-DOPA (Sigma) and ML233 (Tocris) with a contact time of 120s and a flow rate of 45 µl/min. Two-fold dilution series of 5 doses with a high concentration of 500 µM for L-DOPA and 50 µm for ML233 were used.

### Computational modeling

The protein structure of tyrosinase (TYR) was predicted by Alphafold 2^71^, and the N-glycosylation sites of the protein were analyzed using NetNGlyc1.0 tool^72^. Then UCSF Chimera^73^ was used to remove water molecules and irrelevant impurity atoms to retain protein structure only; AMBER14SB was used to calculate protein atomic charge; and the H++3 online tool was used to calculate and assign amino-acid PK values under neutral conditions (pH=7)^74^. The 3D structure of the ML233 molecule was generated by the open-source cheminformatic software package RDKit^75^, and then conformation sampling was conducted. The MMFF94 force field was used to optimize the configuration, and low energy conformation was the output. UCSF Chimera was used to assign AM1-BCC local charge^76^. Finally, AutoDock4.2 software^77^ was used to conduct molecular docking experiments; SiteMap software^78^ was used to predict the optimal binding site; and the predicted binding site was set as the docking center.

Molecular dynamics and binding stability of ML233 and glycosylated tyrosinase was analyzed using the open-source software package Gromacs5.1.5^79^. The simulation system was set as a closed environment; the temperature was set at 289.15K (normal temperature); the pH was set at 7; and the pressure was set at 1 atmosphere 1bar. The periodic boundary setting of the simulation system was centered on the protein, and the minimum distance between the protein edge and the box edge was set at 0.1nm. We used pdb2gmx tool to convert the receptor structure topology file into a file recognized by GROMACS. The force field parameter was AMBEff14SB^80^; the AmberTools tool^81^ was used to convert the ML233 molecule into a topology file recognized by GROMACS in itp format; and the GAFF force field was used to deal with ligand atoms^82^. TIP3P water molecules were added to simulate the water environment, and NaCl solvent was used to balance the system charge.

Molecular mechanics with generalized Born and surface area solvation (MM/GBSA)^83^ was used to calculate the binding energy (ΔG bind) between the TYR protein and ML233. The MMPBSA.py program^84^ integrated with AmberTools was used to calculate the binding energy according to the following formula: ΔGbind = ΔH -TΔS ≈ΔGsolv+ΔGGAS-TΔS (1); ΔGGAS =ΔEint+ΔEvdw + ΔEele (2); ΔGsolv = ΔEsurf +ΔEGB (3).

### Image acquisition

Images were acquired using a Zeiss Axio Zoom.V16 (ref: 435080-9031-000, Carl Zeiss Microscopy, Germany) equipped with a Zeiss Plan Z 1.0x/0.25 objective lens (ref: 435282-9100-000, Carl Zeiss Microscopy, Germany), a transillumination base 300 (ref: 435533-9500-000, Carl Zeiss Microscopy, Germany) equipped with a transillumination top 450 mot (ref: 435500-9000-000, Carl Zeiss Microscopy, Germany) in brightfield mode, and a mechanical stage 150*100 Mot (ref: 435465-9000-000, Carl Zeiss Microscopy, Germany) equipped with an insert plate S, glass 237×157×3 mm (ref:435465-9053-000 Carl Zeiss Microscopy, Germany).

A Zeiss Axiocam 506 color was used, in-color mode (ref: 426556-0000-000, Carl Zeiss Microscopy, Germany) controlled with Zen 3.1 (Carl Zeiss Microscopy, Germany), at zoom 63x, 5 ms exposure, binning 1, at resolution of 2752*2208 pixels, in 14 bit and saved in CZI format. Z-stack images were collected with a step size of 12 microns, using the Focus Motor 3, Central Profile Column 490 mm (ref: 435403-9000-000, Carl Zeiss Microscopy, Germany). Z-stacks were processed using the Zen module extended focus, with the parameters Wavelets, High Z-stack alignment.

Axial length of the eye was quantified using Fiji software after imaging of the larvae at 2 dpf using a Zeiss Axio Zoom V16.

The images underwent modifications using Fiji software. To ensure consistency in image processing, a macro was devised in collaboration with our team and the Light Microscopy Facility at MDIBL and is made available upon request.

## ACKNOWLEDGMENTS

Research reported in this manuscript was supported by an Institutional Development Award (IDeA) from the National Institute of General Medical Sciences of the National Institutes of Health under grant numbers P20GM103423, P20GM104318 and P20GM144265. Image collection. Processing and analysis for this manuscript was performed with the assistance of Dr. Frederic Bonnet and the MDI Biological Laboratory Light Microscopy Facility (RRID:SCR_019166), which is supported by an Institutional Development Award (IDeA) from the National Institute of General Medical Sciences of the National Institutes of Health under grant number P20GM103423“.

## Authors’ contribution

*Conceptualization*: Romain Menard, Aissette Baanannou, Joel, Graber, James Strickland, Romain Madelaine. *Investigation*: Romain Menard, Aissette Baanannou, Caroline Halluin, Dexter Morse, Sadie Kuhn, Romain Madelaine. *Data curation*: Romain Menard, Aissette Baanannou, Dexter Morse, Romain Madelaine. *Formal analysis*: Romain Menard, Aissette Baanannou, Dexter Morse. Funding acquisition: James Strickland, Romain Madelaine. *Supervision*: Romain Madelaine. *Writing – original draft:* Romain Menard, Aissette Baanannou, Romain Madelaine.

## Data availability

All data, reagents and genetic tools are available upon request.

## Competing interest

Romain Menard, Aissette Baanannou and Romain Madelaine hold a pending patent # PCT/US24/49268 for “MELANOGENESIS INHIBITION COMPOSITIONS AND METHODS OF USE THEREOF”.

## SUPPLEMENTARY FIGURES

**Figure S1.**
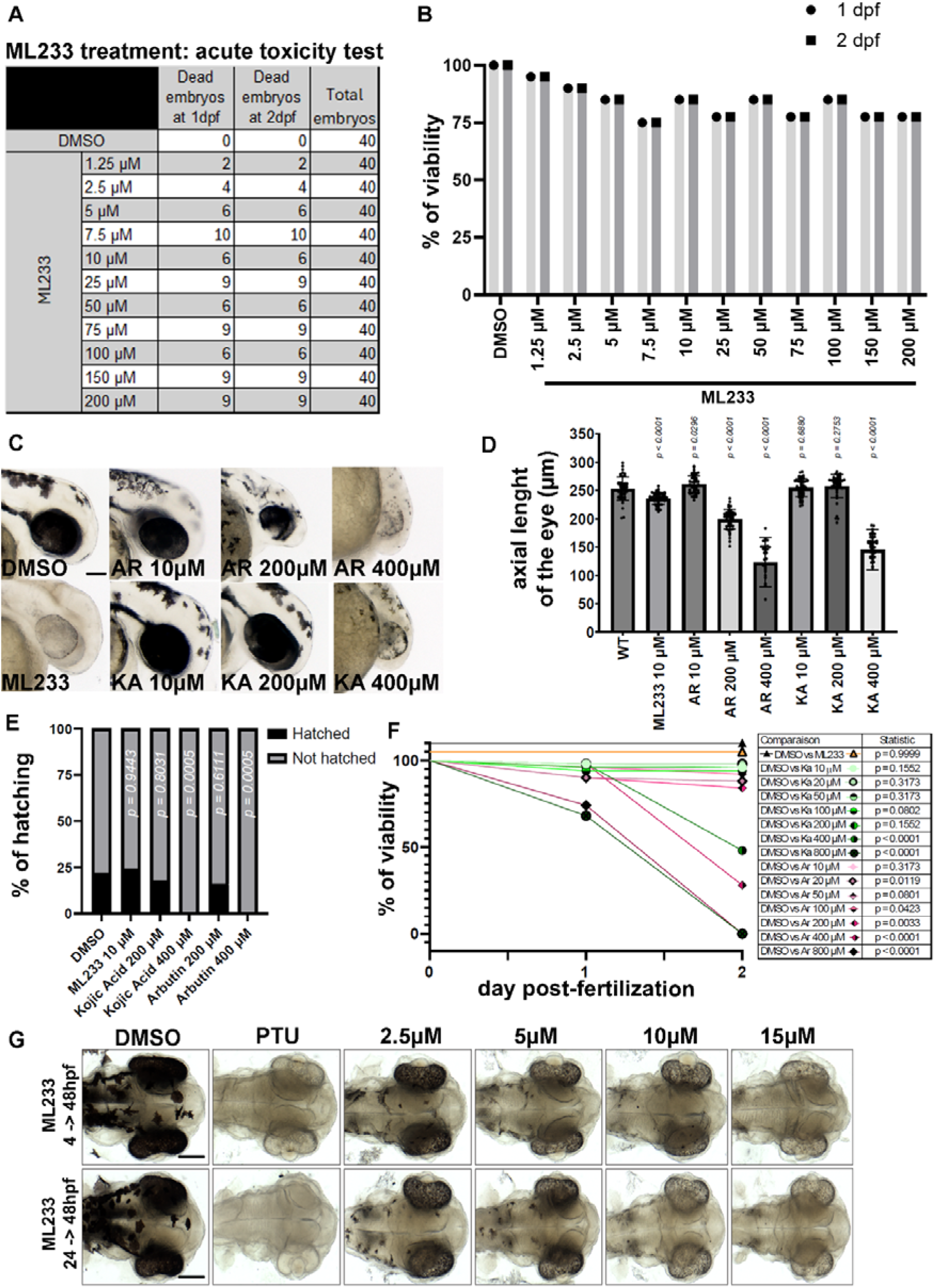
Dose-dependent effect of ML233 regulation of skin pigmentation. **(A)** Table showing the survival rate at 2 dpf for the different concentrations of ML233 tested. **(B)** Zebrafish embryo survival (% of viability) was quantified in control (DMSO) and ML233-treated embryos by acute toxicity test (n=40). **(C)** Pigmentation of the zebrafish eye after DMSO, ML233, kojic acid (KA), or arbutin (AR) treatment from 4 to 48 hpf**. (D)** Quantification of eye size (axial length in µm) following DMSO, ML233, kojic acid (KA), or arbutin (AR) treatment from 4 to 48 hpf (n≥30 eyes). t-test, two-tailed, unpaired. Error bars represent s.d. **(E)** Quantification of percentage (%) of hatching embryos after DMSO, ML233, kojic acid (KA), or arbutin (AR) treatment from 4 to 48 hpf (n=50). Fisher’s exact test. **(F)** Zebrafish embryo survival (% of viability) was quantified in DMSO-, ML233-, kojic acid (KA),- or arbutin (AR)-treated embryos by acute toxicity test until 2 dpf (n=50). Kaplan-Meier analysis. **(G)** Skin pigmentation of DMSO- or ML233-treated embryos at 48 hpf after treatment between 4 and 48 hpf (top) or 24 and 48 hpf (bottom). Dorsal view of the head. Scale bars: 100 μm.

**Figure S2.**
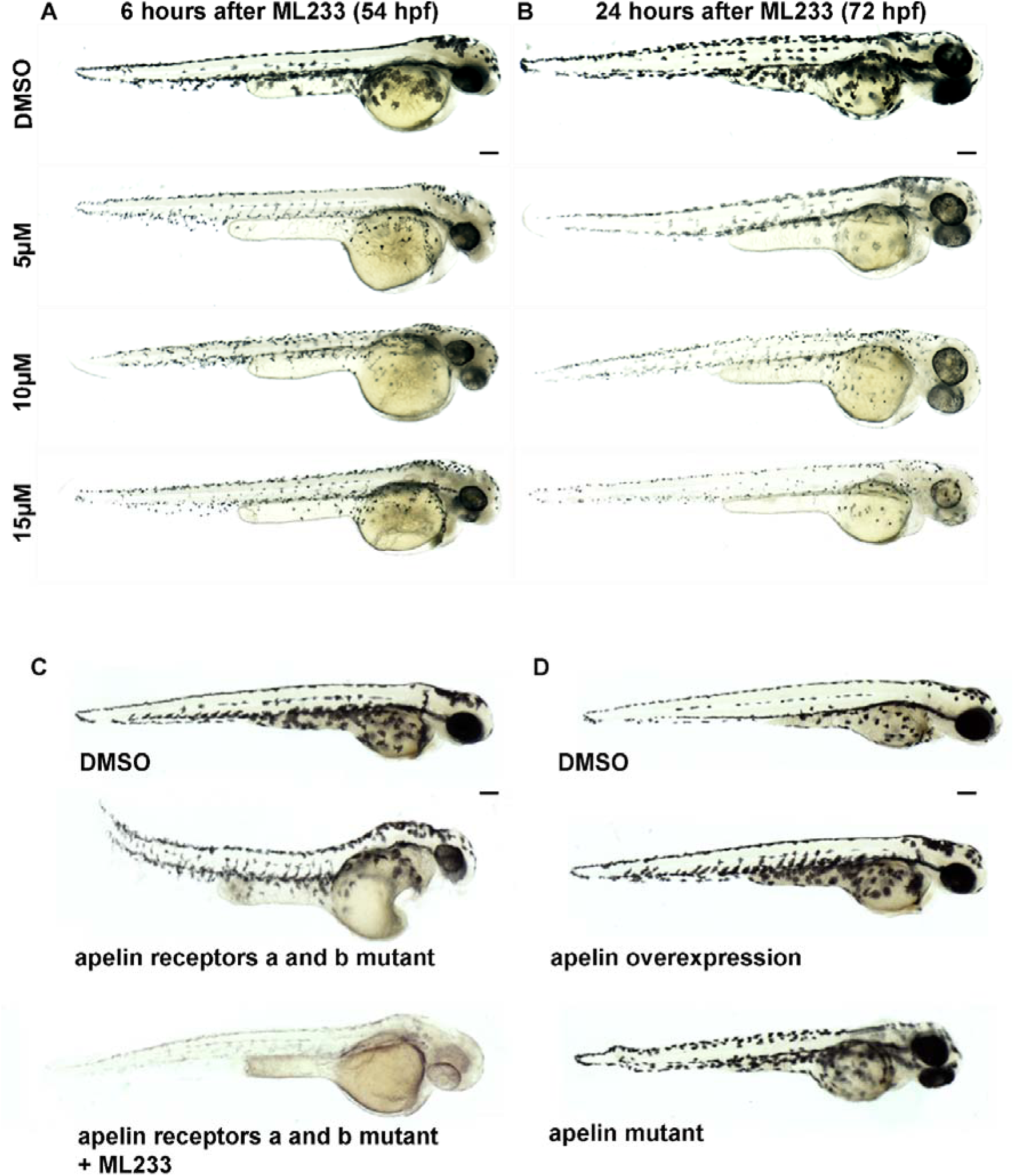
ML233-dependent inhibition of melanin production is independent of the apelin signaling pathway. **(A)** Skin pigmentation of DMSO- or ML233-treated embryos at 54 hpf after treatment between 48 and 54 hpf. **(B)** Skin pigmentation of DMSO- or ML233-treated embryos at 48 hpf after treatment between 48 and 72 hpf. **(C)** Skin pigmentation of DMSO, apelin receptor double-KO embryos, or ML233-treated apelin receptor double-KO embryos (between 24 and 72 hpf) at 72 hpf. **(D)** Skin pigmentation of DMSO-treated, apelin ubiquitous overexpression, or apelin mutant embryos at 72 hpf. Scale bars: 100 μm.

**Figure S3.**
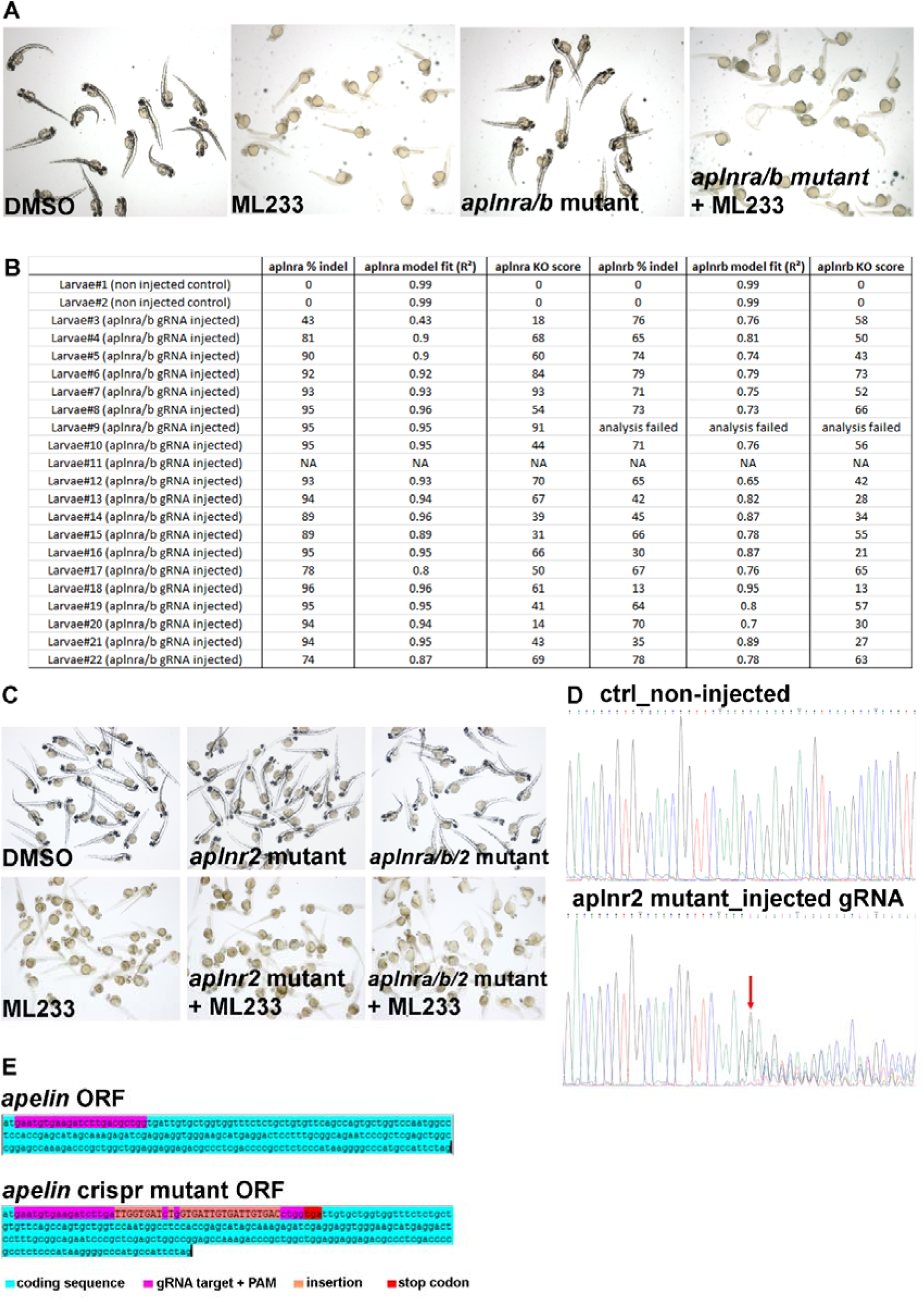
Apelin signaling is not involved in regulation of melanin production. **(A)** Skin pigmentation of DMSO, ML233, apelin receptor mutant, or ML233-treated apelin receptor mutant embryos ML233 (between 24 and 72 hpf) at 72 hpf. **(B)** Quantification of the percentage of indel in *aplnra* and *aplnrb* locus after CRISPR/Cas-9 mutagenesis (n=20). **(C)** Skin pigmentation of untreated and ML233-treated DMSO, *aplnr2* mutant, and *aplnra-aplnrb-aplnr2* mutant embryos (treatment was between 4 and 48 hpf) at 48 hpf. **(D)** Representative chromatograph sequences of control (top) and mutated (bottom) *aplnr2* coding sequences (Sanger sequencing of individual embryos). The red arrow shows the INDEL introduced in the locus targeted by the *aplnr2* gRNA. **(E)** Schematic representation and nucleotide sequence of the *apelin* gene coding sequence in control or CRISPR/Cas-9 mutant. Early stop codon (red) appears in the *apelin* mutant.

**Figure S4.**
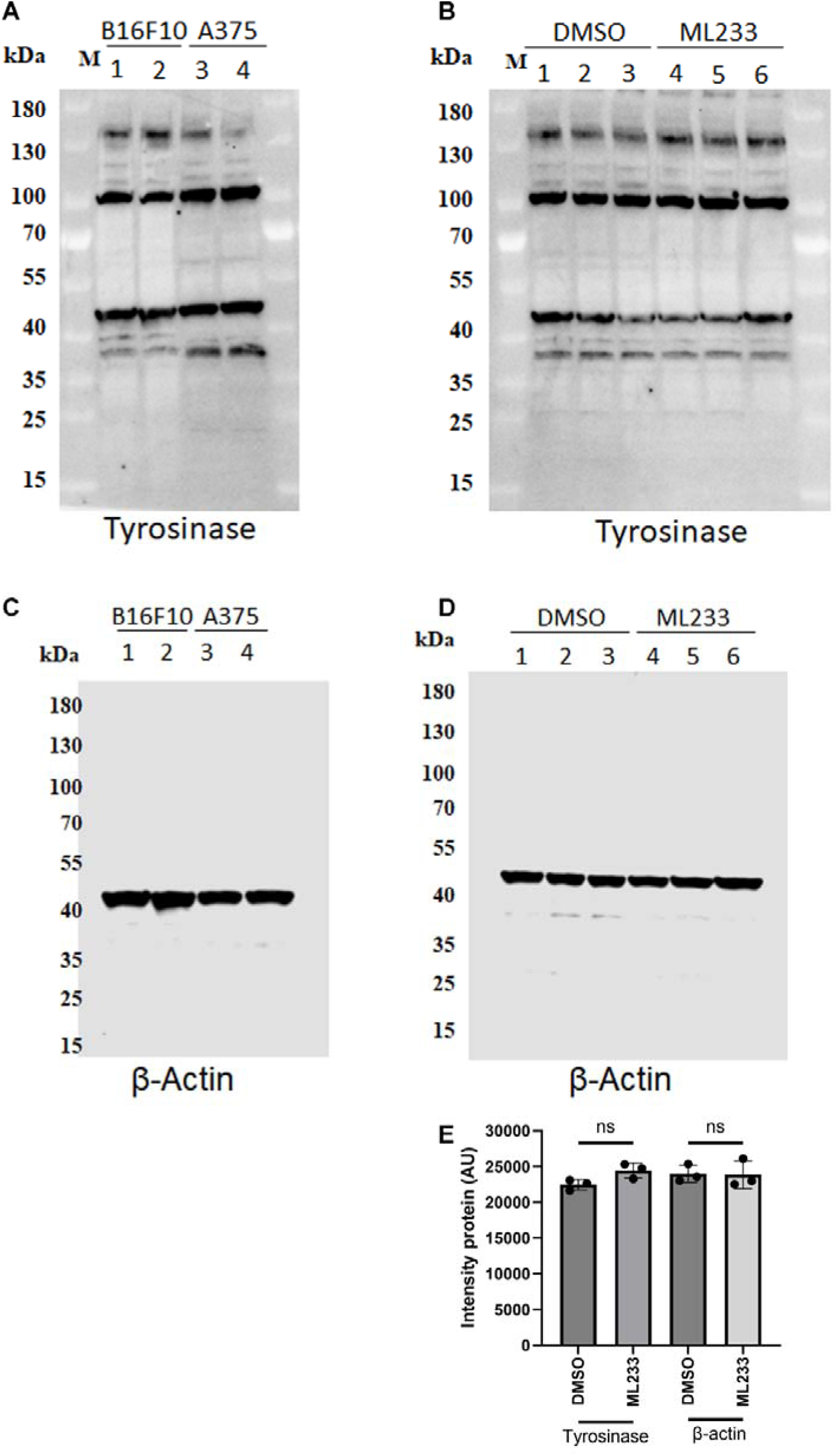
Tyrosinase protein expression is not affected by ML233 treatment. **(A)** Expression of tyrosinase protein was analyzed by western blot in murine (B16F10) or human (A375) melanoma cells (n=3). **(B)** Expression of tyrosinase protein was analyzed by western blot in murine (B16F10) melanoma cells after DMSO or ML233 treatment (n=3). **(C)** Expression of beta-actin protein was analyzed by western blot in murine (B16F10) or human (A375) melanoma cells (n=2). **(D)** Expression of beta-actin protein was analyzed by western blot in murine (B16F10) melanoma cells after DMSO or ML233 treatment (n=3). **(E)** Tyrosinase or beta-actin protein expression was quantified by intensity of protein expression (AU) after DMSO or ML233 treatment. Error bars represent s.d. Statistical significance is determined by t-test, two-tailed, unpaired.

**Figure S5.**
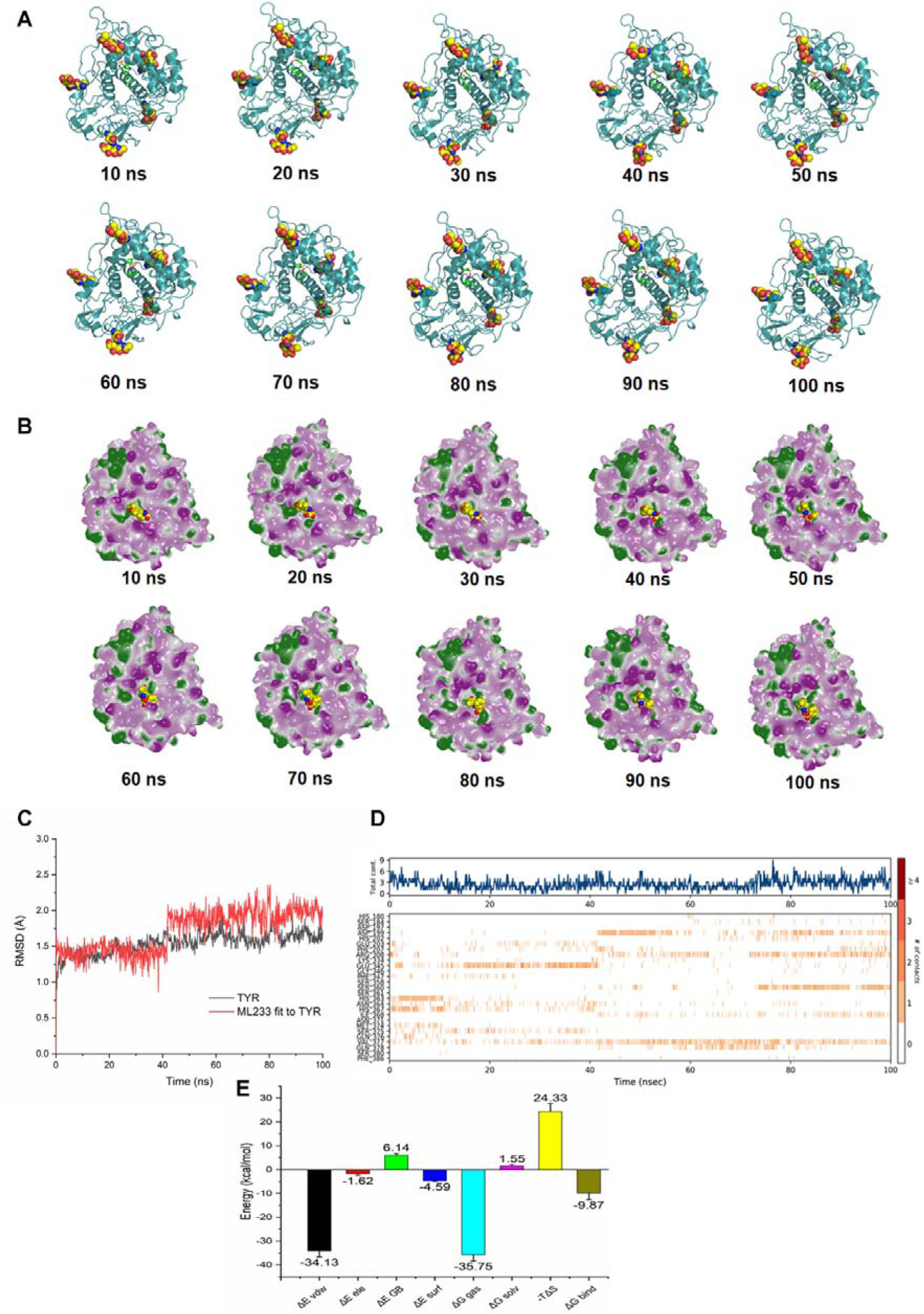
ML233 binds the human TYR protein with stability. **(A)** Representation of the molecular dynamics during 100 ns simulation. Proteins are shown using cyan ribbons, glycosylation sites are shown using spheres, and ML233 is shown using a structure of greensticks **(B)** Distribution of hydrophobic area and hydrophilic area of TYR protein during dynamic simulation. Hydrophobic area is shown using green, and hydrophilic area is shown using purple. ML233 is shown using yellow spheres. **(C)** Stability analysis between TYR protein and ML233 molecule during molecular dynamics. Black curve represents the RMSD of the TYR protein over time, and red curve represents the RMSD of ML233 over time. **(D)** Analysis of the critical amino acids interacting with ML233 and the number of contacts. **(E)** Quantification of the binding free energy during TYR and ML233 interaction.

## SUPPLEMENTARY MATERIALS

**Table S1.** GSEA (Gene Set Enrichment Analysis) between *aplnr* KO embryos and *aplnr* KO embryos treated with 0.5 µM of ML233 for 3 hours.

**Table S2.** GSEA (Gene Set Enrichment Analysis) between *aplnr* KO embryos and *aplnr* KO embryos treated with 0.5 µM of ML233 for 24 hours.

**Table S3.** GO (Gene Ontology) between *aplnr* KO embryos and *aplnr* KO embryos treated with 0.5 µM of ML233 for 3 hours.

**Table S4.** GO (Gene Ontology) between *aplnr* KO embryos and *aplnr* KO embryos treated with 0.5 µM of ML233 for 24 hours.

**Table S5.** KEGG analysis between *aplnr* KO embryos and *aplnr* KO embryos treated with 0.5 µM of ML233 for 3 hours.

**Table S6.** KEGG analysis between *aplnr* KO embryos and *aplnr* KO embryos treated with 0.5 µM of ML233 for 24 hours.

**Video S1.** Movie of the molecular dynamic during simulation of TYR-ML233 interaction.

