## Supplementary material for "The small molecule ML233 is a direct inhibitor of tyrosinase function": TableS1

Table 1

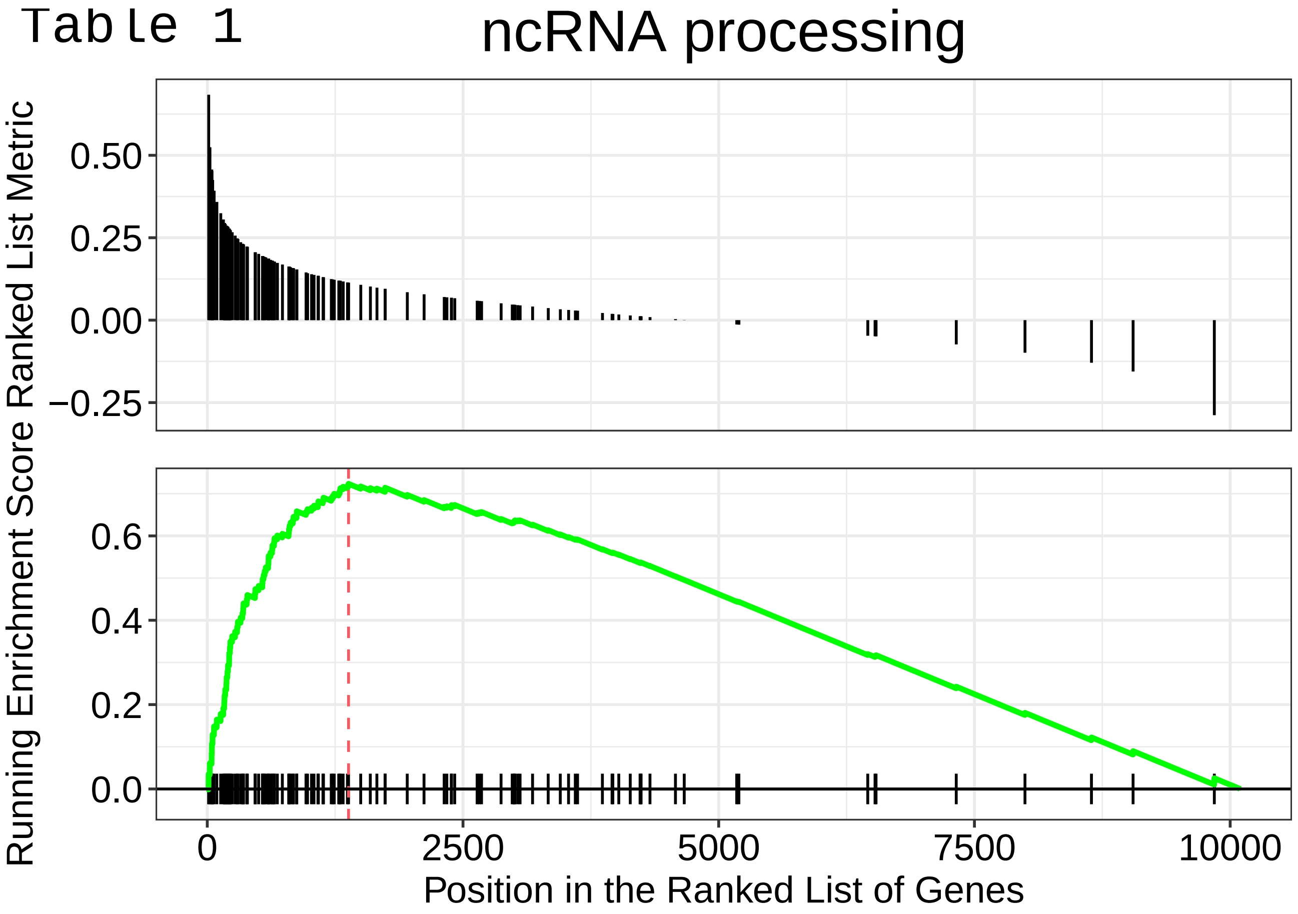
ncRNA processing
Running Enrichment Score Ranked List Metric
0.50
0.25
0.00
−0.25
0.6
0.4
0.2
0.0
0
2500
5000
7500
10000
Position in the Ranked List of Genes

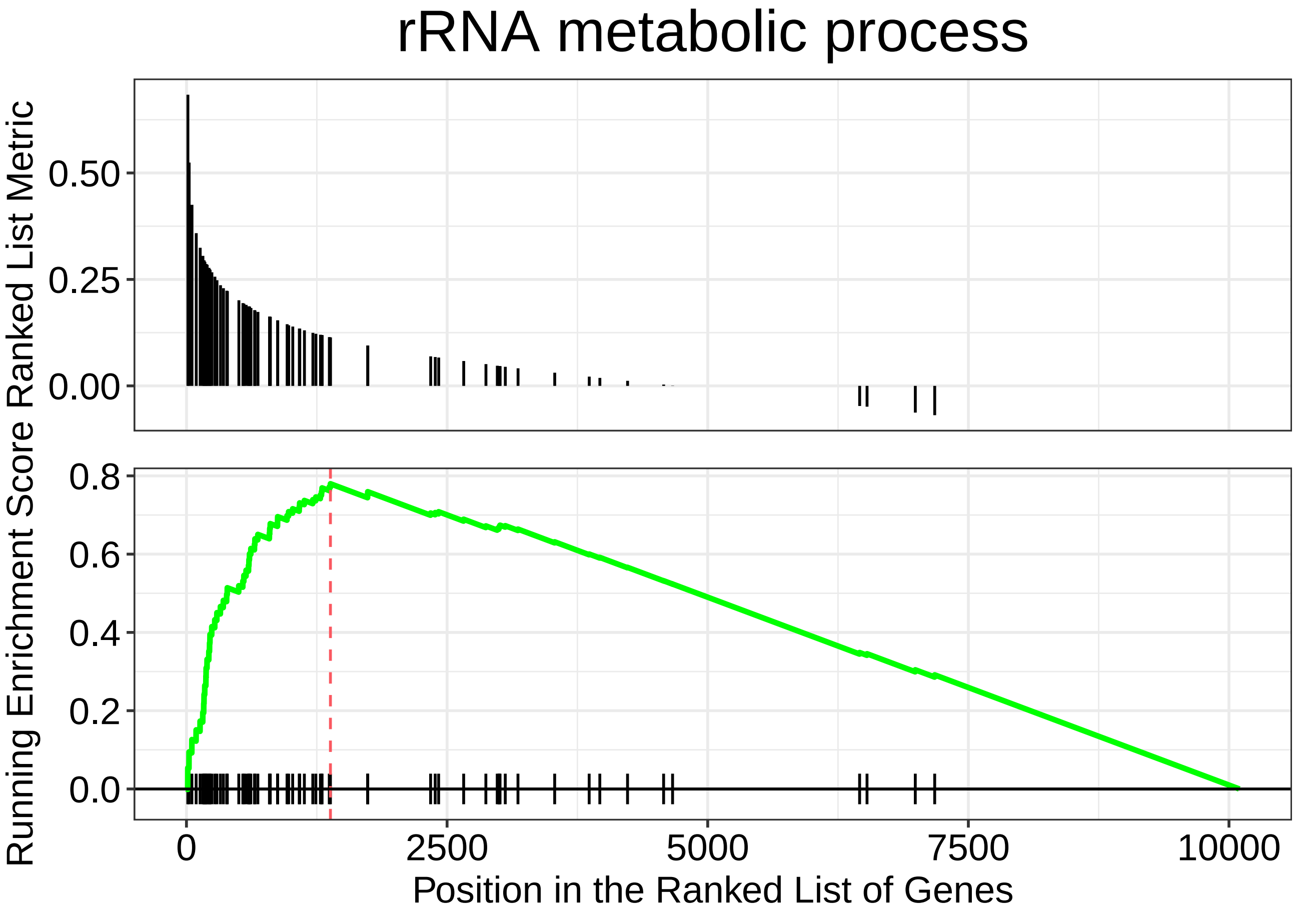
rRNA metabolic process
Ranked List Metric
0.50
0.25
0.00
Running Enrichment Score
0.8
0.6
0.4
0.2
0.0
0
2500
5000
7500
10000
Position in the Ranked List of Genes

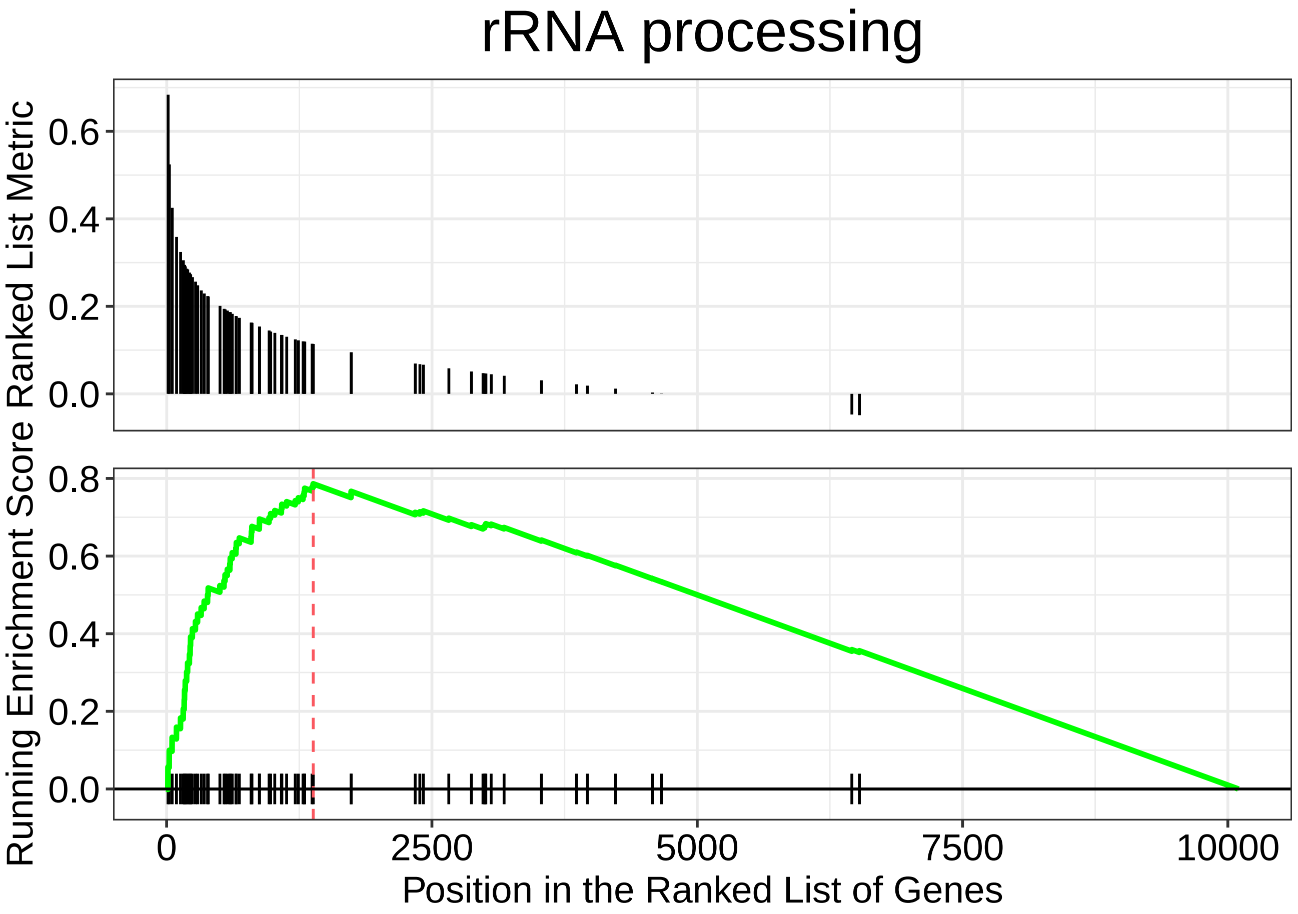

rRNA processing
Ranked List Metric
0.6
0.4
0.2
0.0
Running Enrichment Score
0.8
0.6
0.4
0.2
0.0
0
2500
5000
7500
10000
Position in the Ranked List of Genes

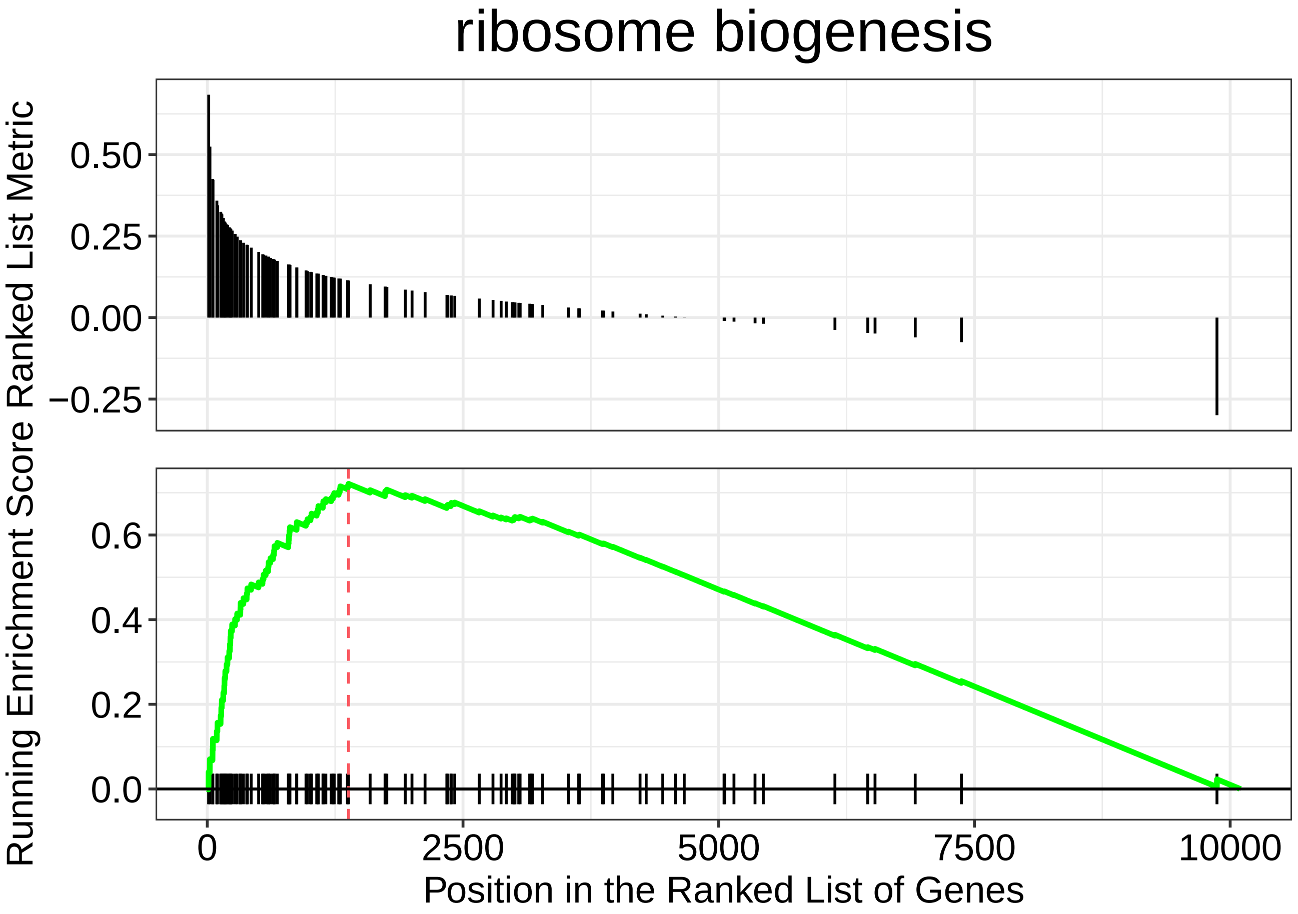

ribosome biogenesis
Running Enrichment Score Ranked List Metric
0.50
0.25
0.00
−0.25
0.6
0.4
0.2
0.0
0
2500
5000
7500
10000
Position in the Ranked List of Genes

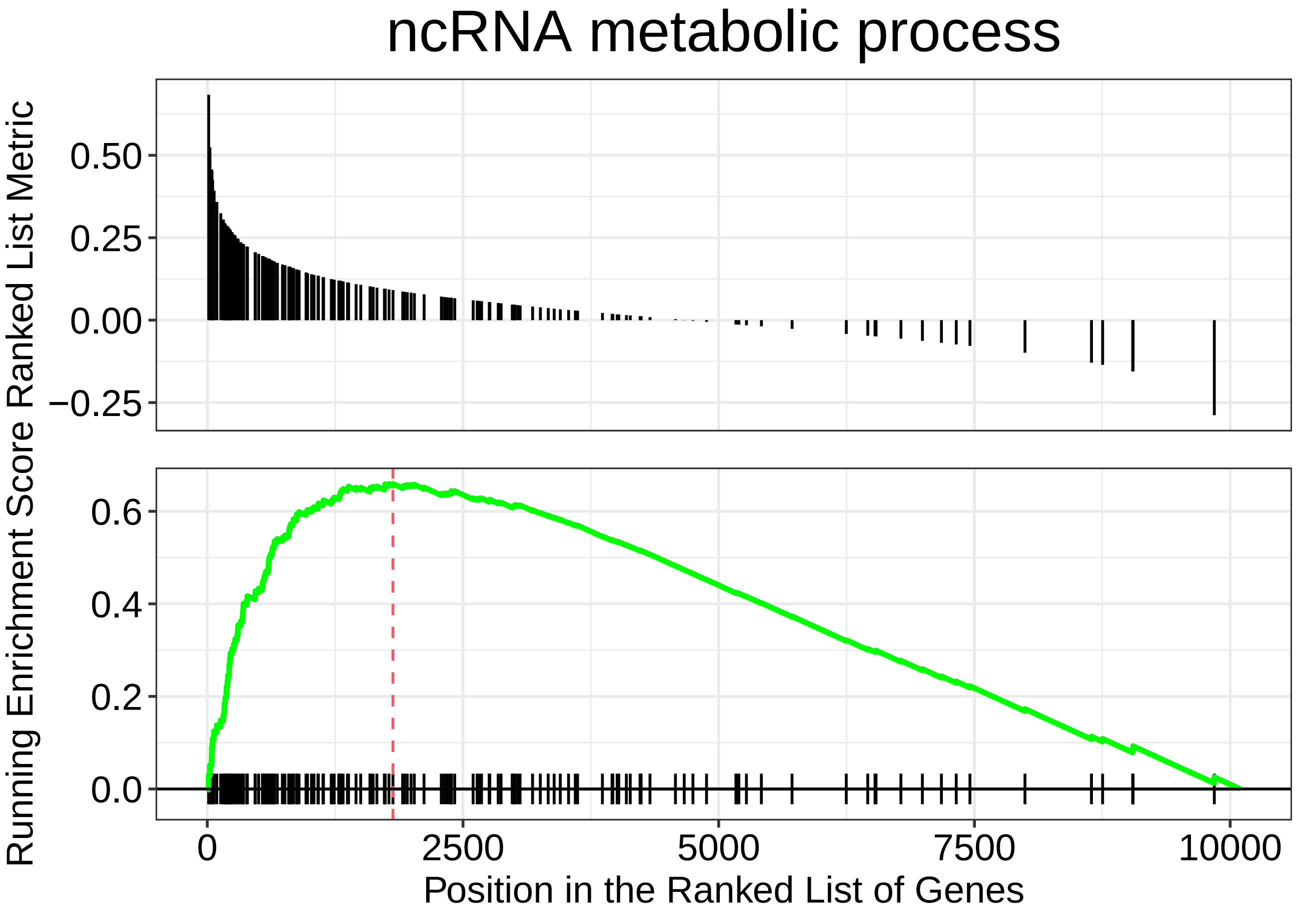

ncRNA metabolic process
Ranked List Metric
0.50
0.25
0.00
−0.25
Running Enrichment Score
0.6
0.4
0.2
0.0
0
2500
5000
7500
10000
Position in the Ranked List of Genes

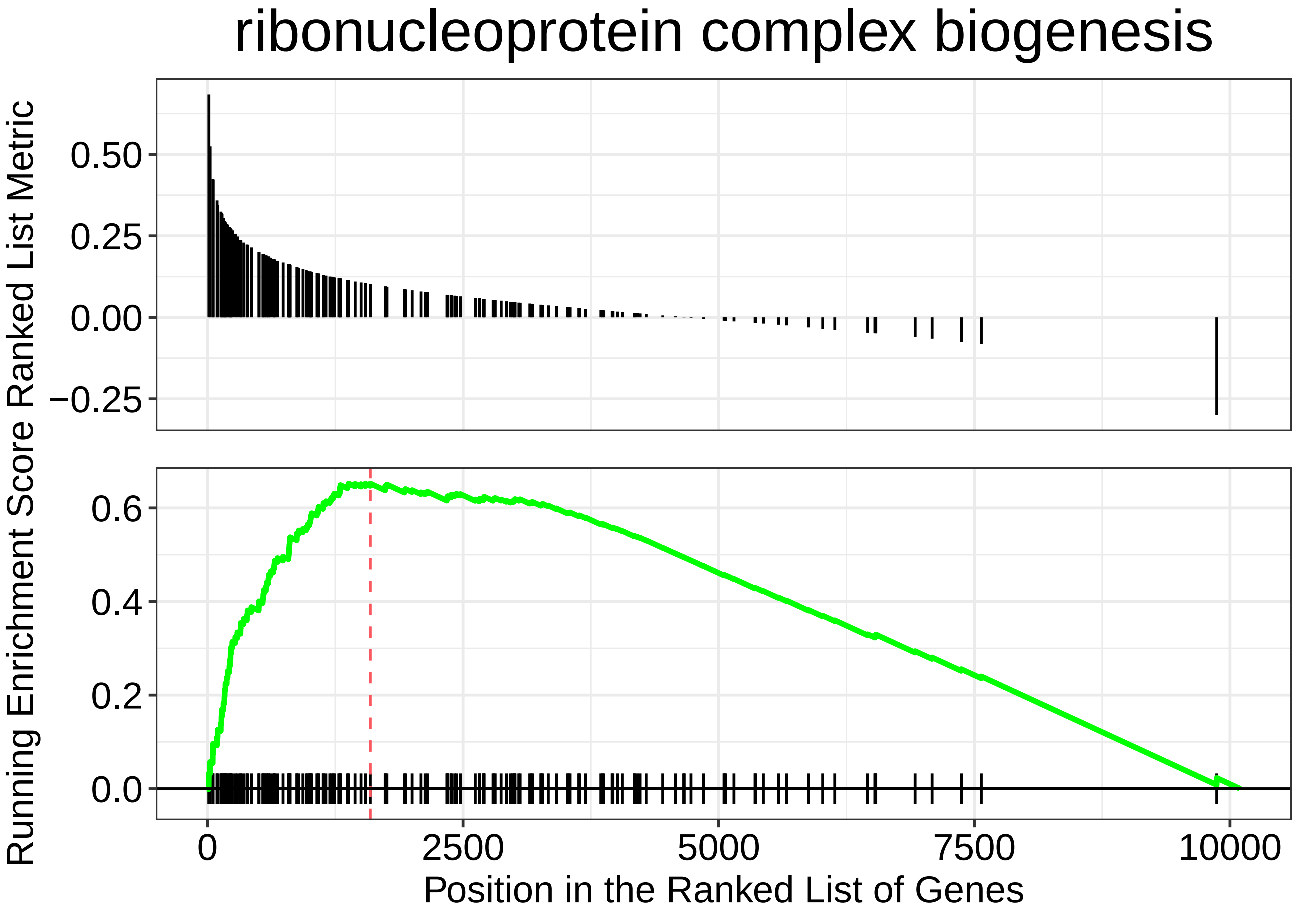
ribonucleoprotein complex biogenesis
Running Enrichment Score Ranked List Metric
0.50
0.25
0.00
−0.25
0.6
0.4
0.2
0.0
0
2500
5000
7500
10000
Position in the Ranked List of Genes

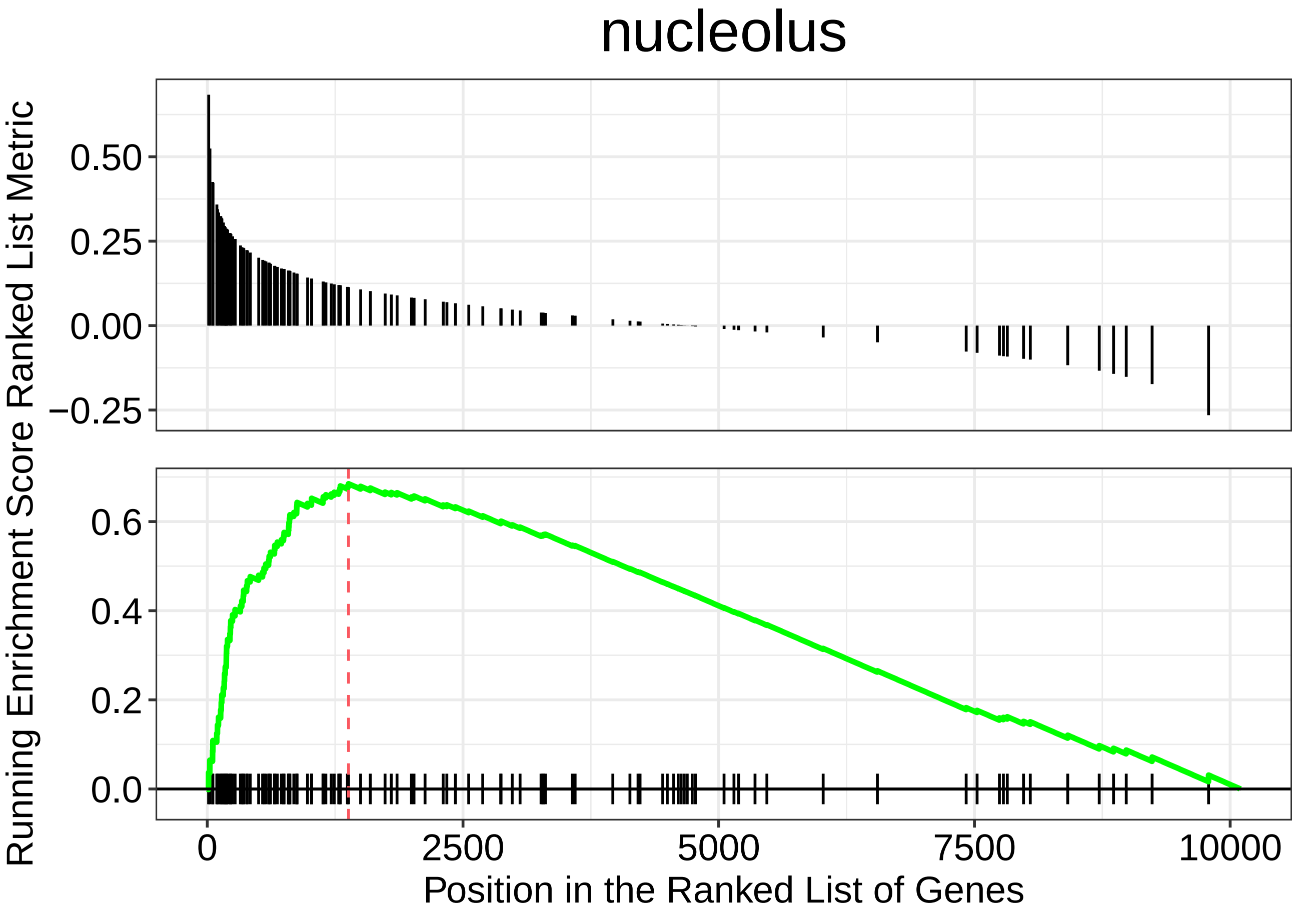

nucleolus
Running Enrichment Score Ranked List Metric
0.50
0.25
0.00
−0.25
0.6
0.4
0.2
0.0
0
2500
5000
7500
10000
Position in the Ranked List of Genes

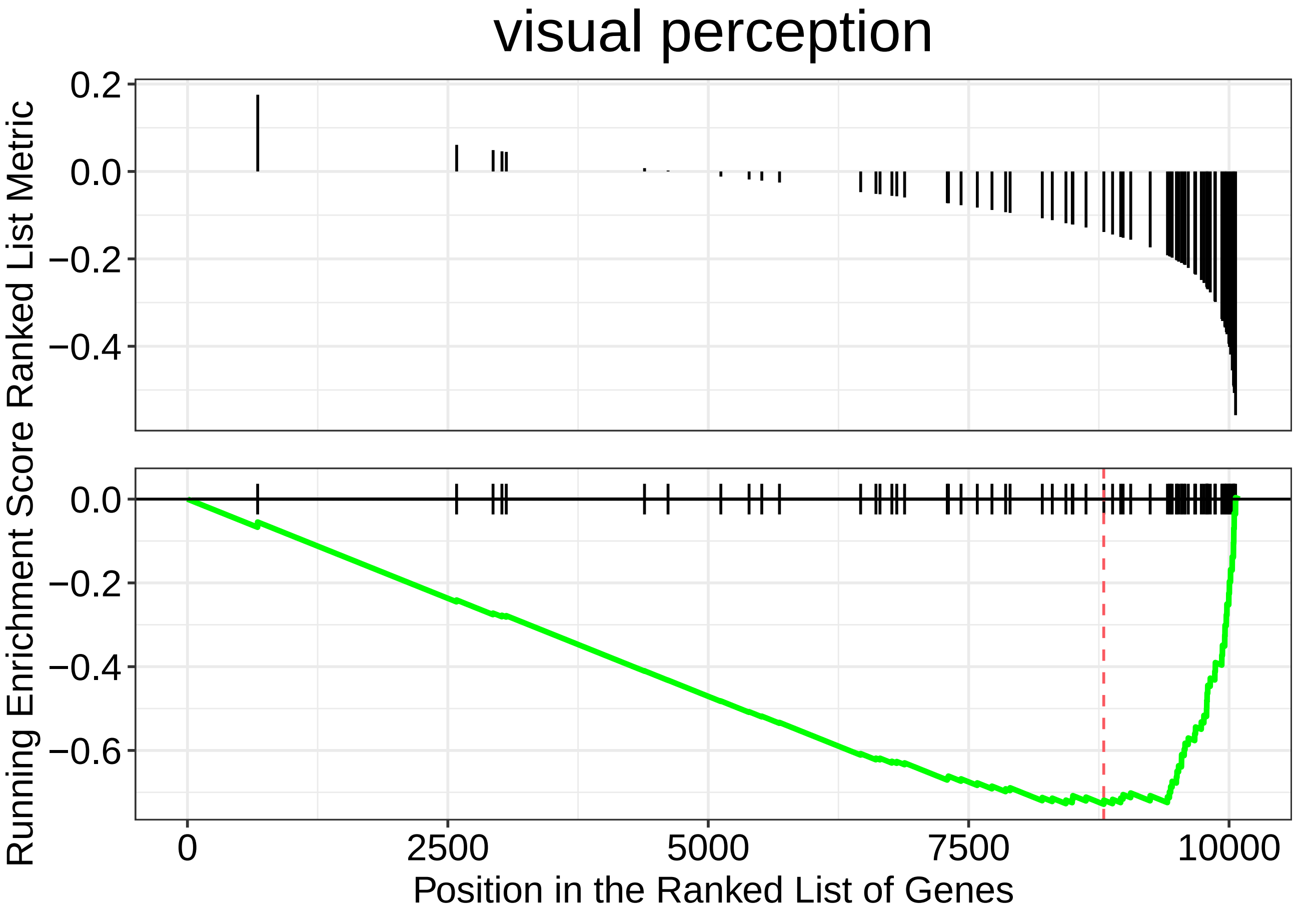

visual perception
Running Enrichment Score Ranked List Metric
0.2
0.0
−0.2
−0.4
0.0
−0.2
−0.4
−0.6
0
2500
5000
7500
10000
Position in the Ranked List of Genes

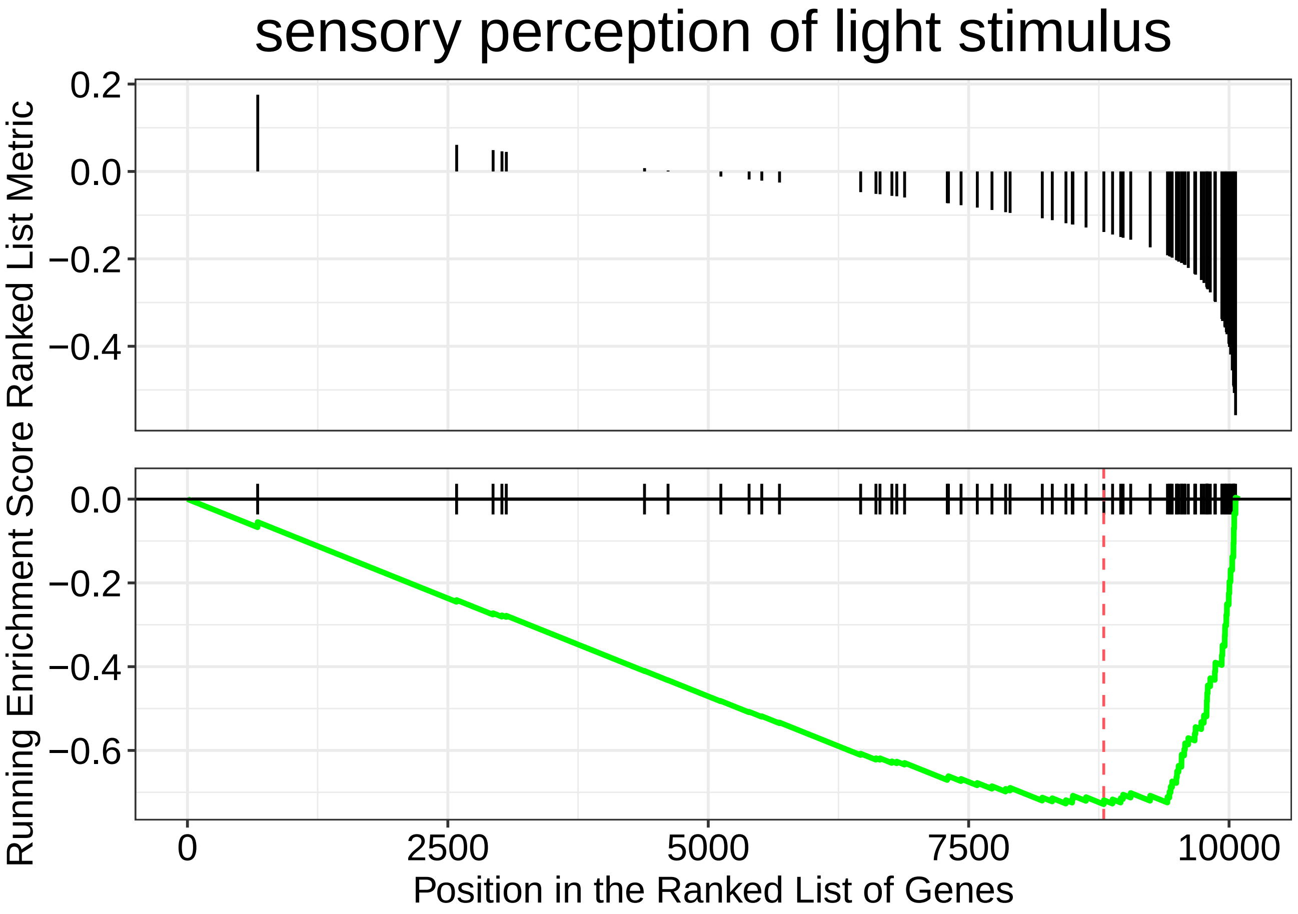
sensory perception of light stimulus
Ranked List Metric
Running Enrichment Score
0.2
0.0
−0.2
−0.4
−0.6
0
2500
5000
7500
10000
Position in the Ranked List of Genes

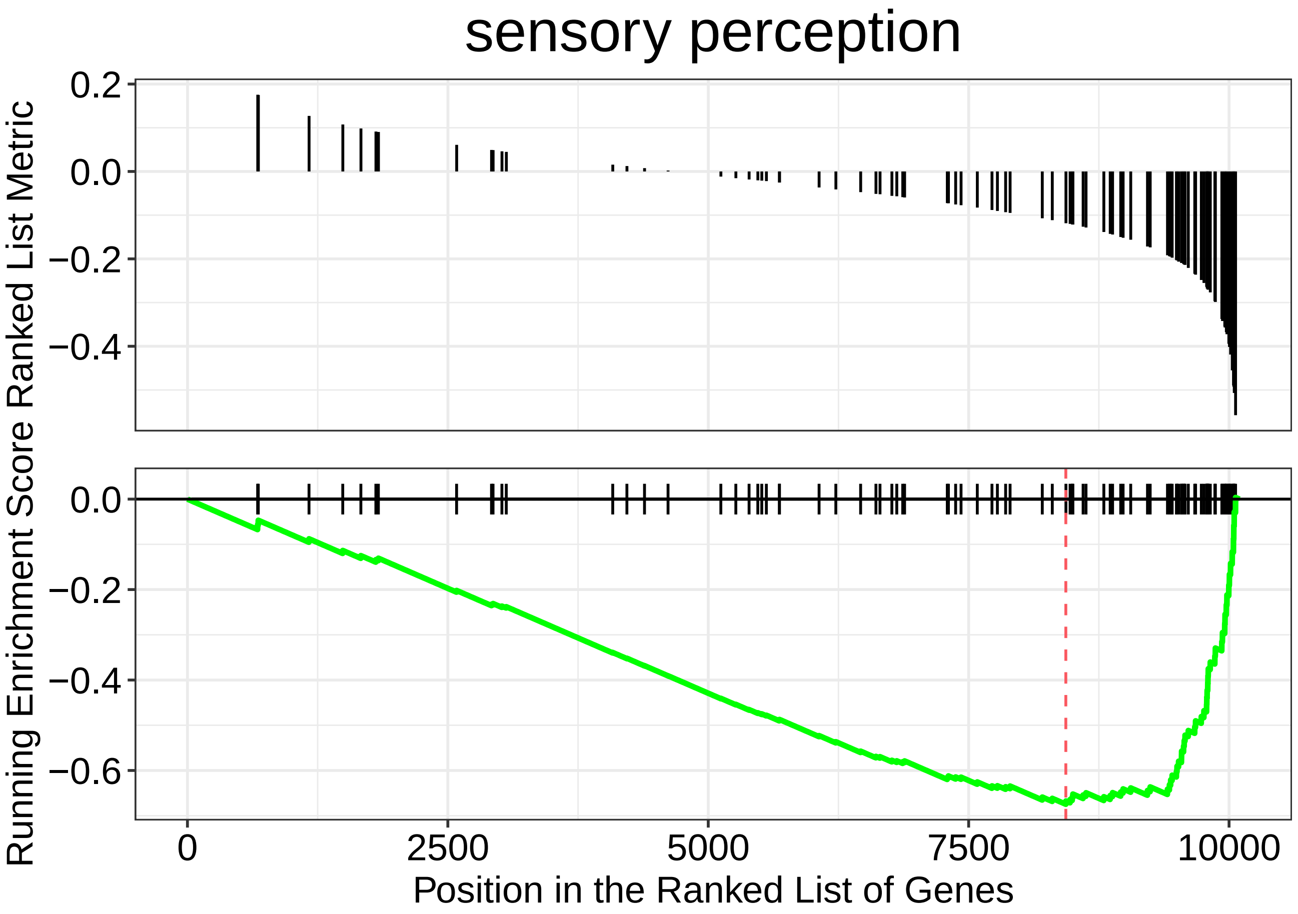
sensory perception
0.2
0.0
−0.2
−0.4
Running Enrichment Score Ranked List Metric
0.0
−0.2
−0.4
−0.6
0
2500
5000
7500
10000
Position in the Ranked List of Genes

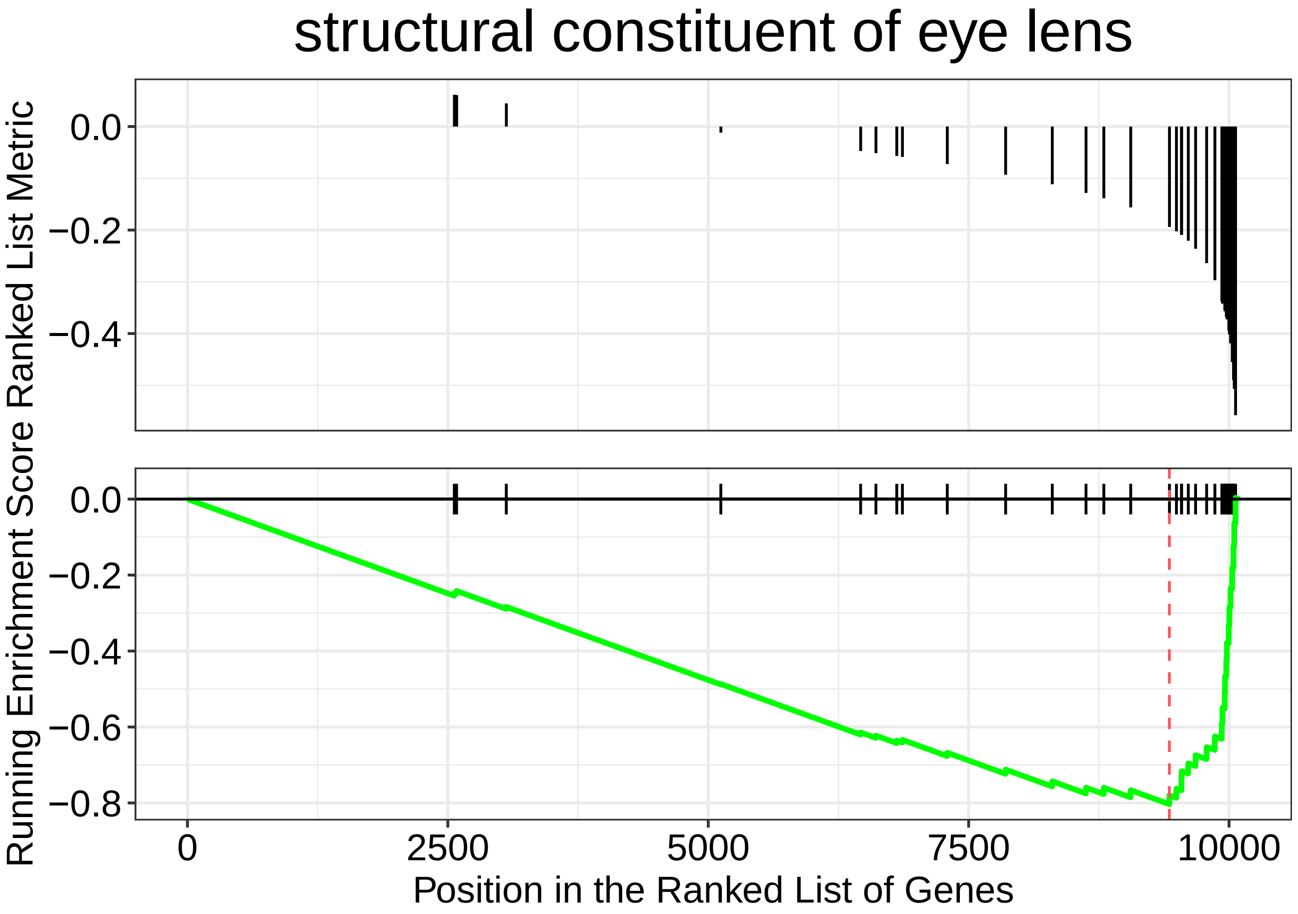

structural constituent of eye lens
Ranked List Metric
0.0
−0.2
−0.4
Running Enrichment Score
0.0
−0.2
−0.4
−0.6
−0.8
0
2500
5000
7500
10000
Position in the Ranked List of Genes

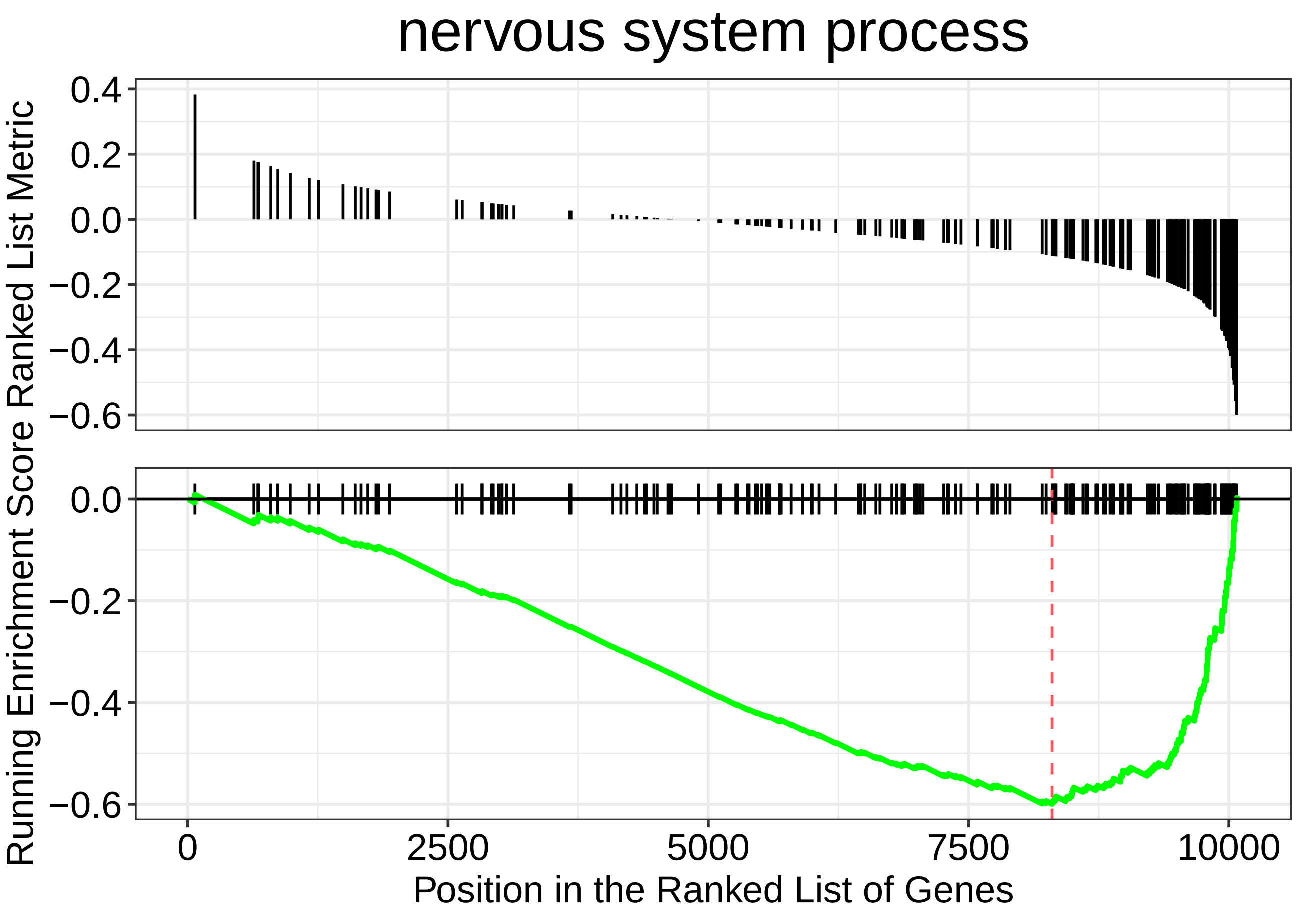
nervous system process
Running Enrichment Score Ranked List Metric
0.4
0.2
0.0
−0.2
−0.4
−0.6
0.0
−0.2
−0.4
−0.6
0
2500
5000
7500
10000
Position in the Ranked List of Genes

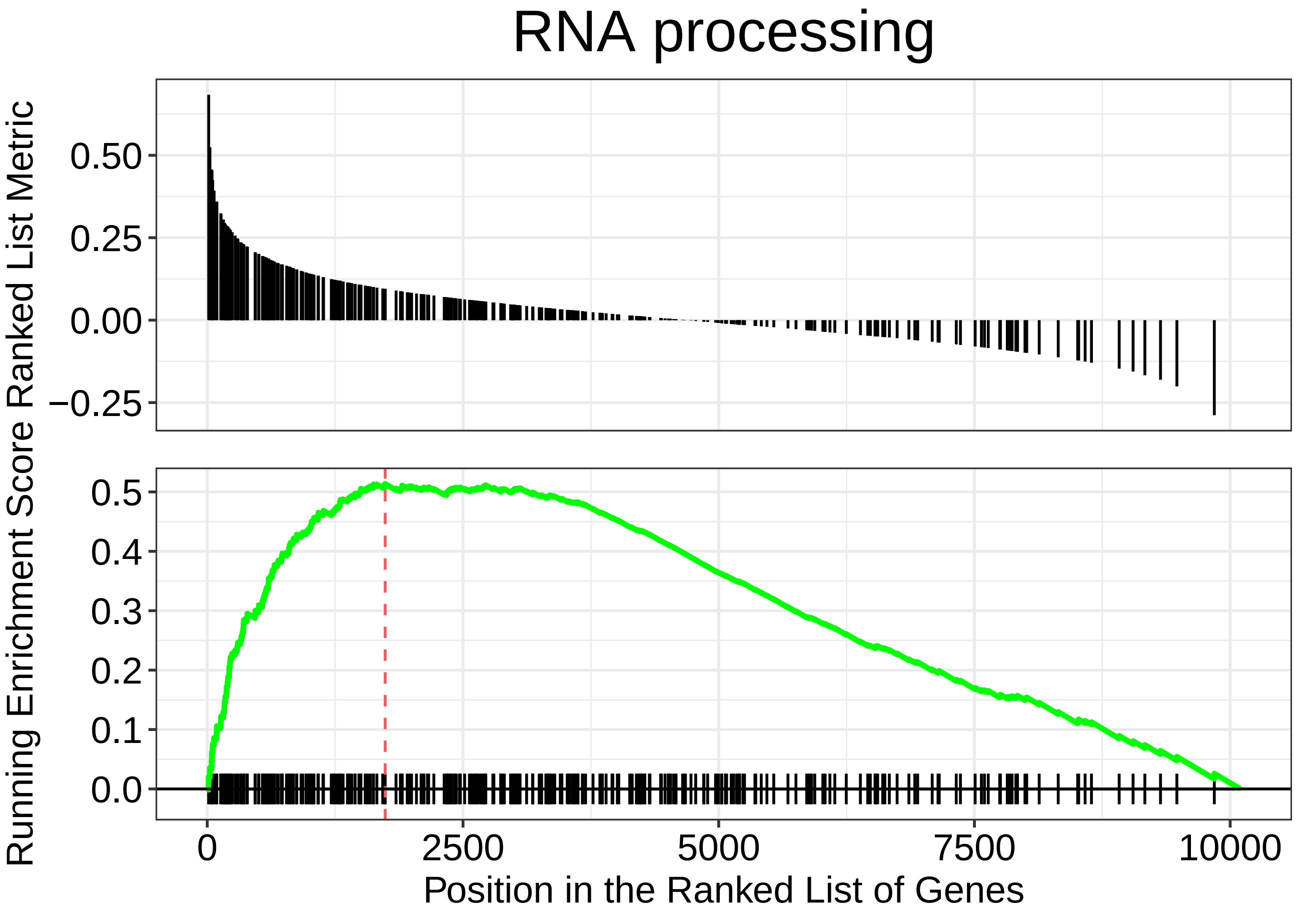
RNA processing
Running Enrichment Score Ranked List Metric
0.50
0.25
0.00
−0.25
0.5
0.4
0.3
0.2
0.1
0.0
0
2500
5000
7500
10000
Position in the Ranked List of Genes

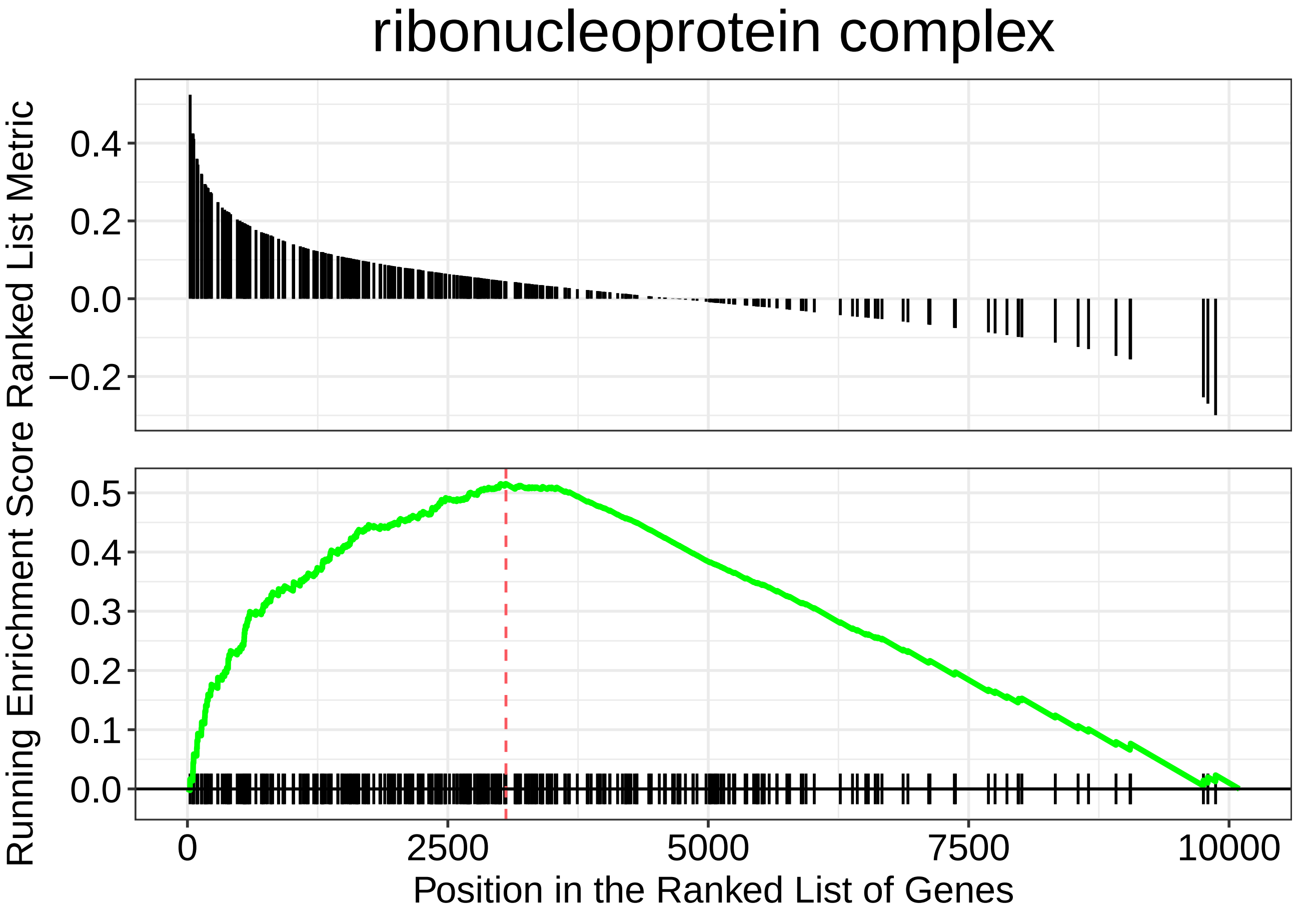

ribonucleoprotein complex
Ranked List Metric
0.4
0.2
0.0
−0.2
Running Enrichment Score
0.5
0.4
0.3
0.2
0.1
0.0
0
2500
5000
7500
10000
Position in the Ranked List of Genes

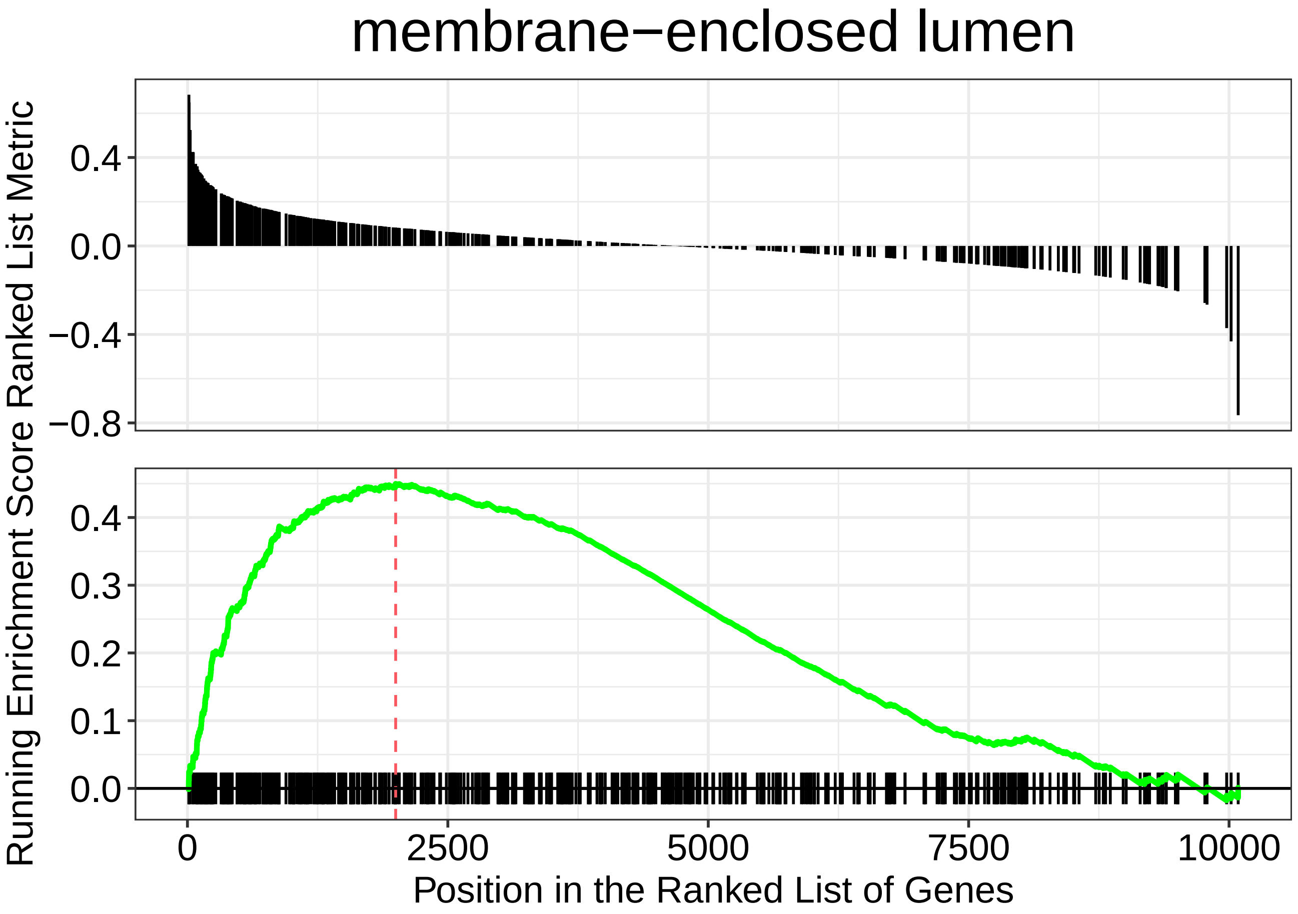
membrane−enclosed lumen
Ranked List Metric
0.4
0.0
−0.4
−0.8
Running Enrichment Score
0.4
0.3
0.2
0.1
0.0
0
2500
5000
7500
10000
Position in the Ranked List of Genes

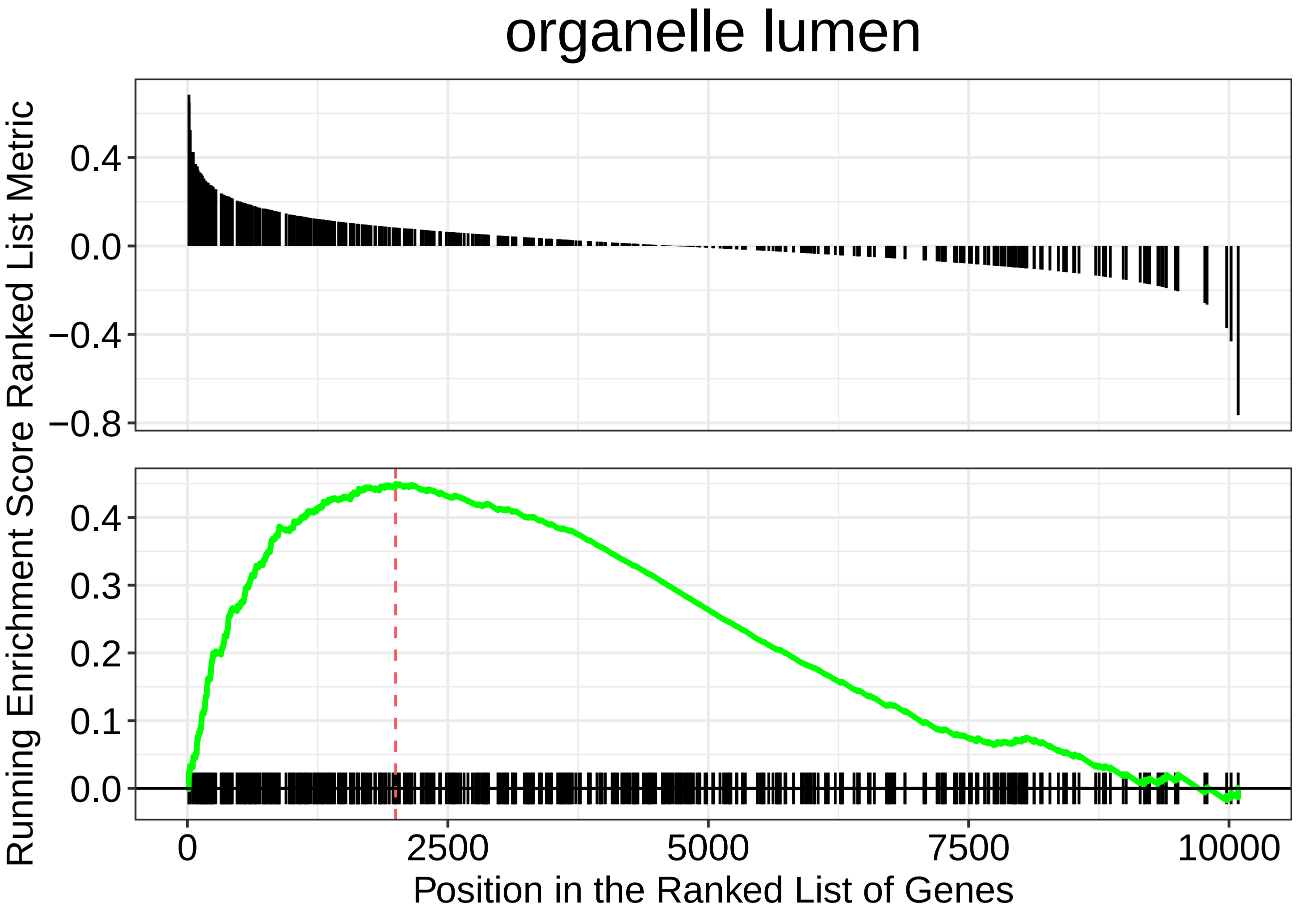

organelle lumen
Running Enrichment Score Ranked List Metric
0.4
0.0
−0.4
−0.8
0.4
0.3
0.2
0.1
0.0
0
2500
5000
7500
10000
Position in the Ranked List of Genes

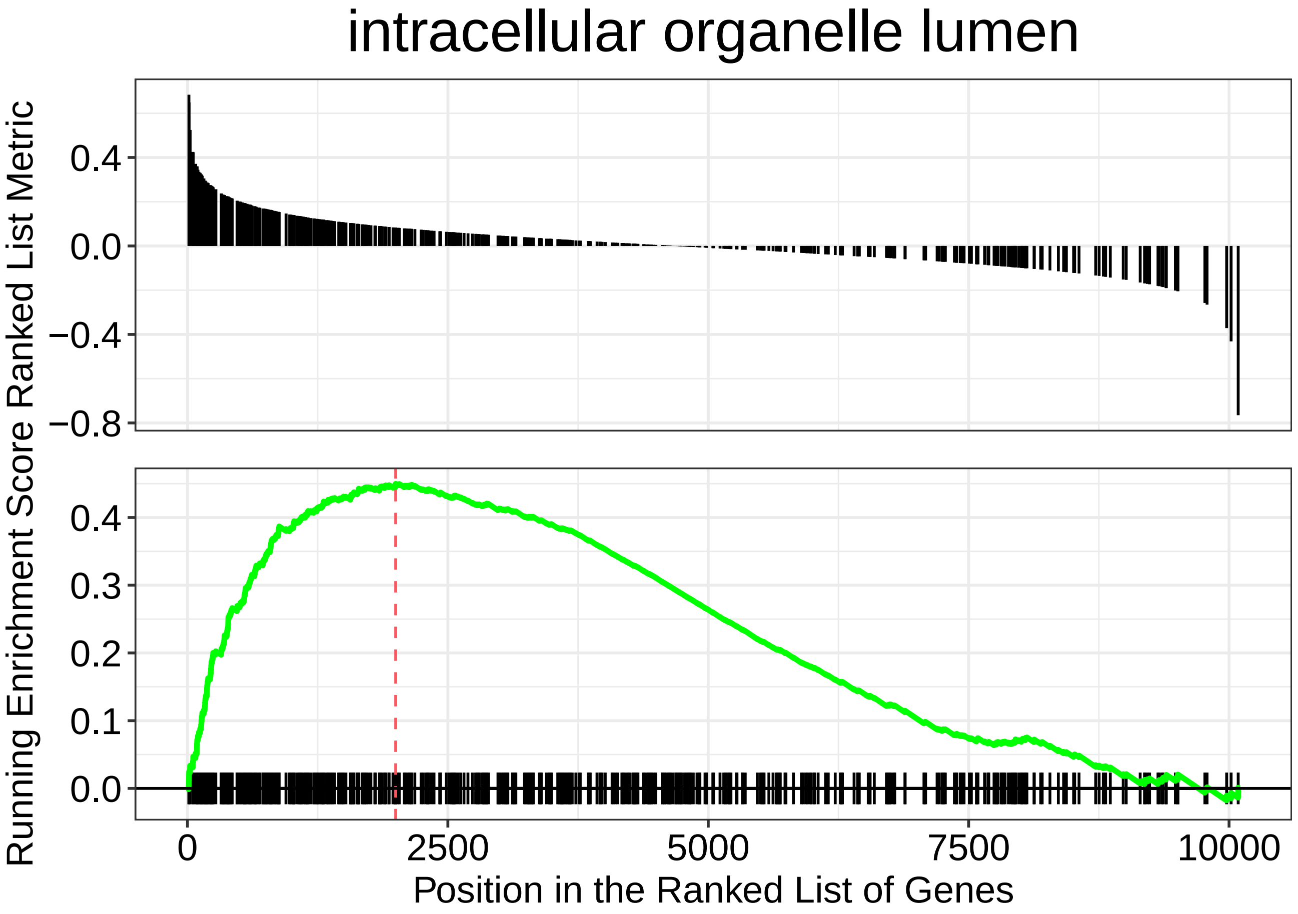
intracellular organelle lumen
Running Enrichment Score Ranked List Metric
0.4
0.0
−0.4
−0.8
0.4
0.3
0.2
0.1
0.0
0
2500
5000
7500
10000
Position in the Ranked List of Genes

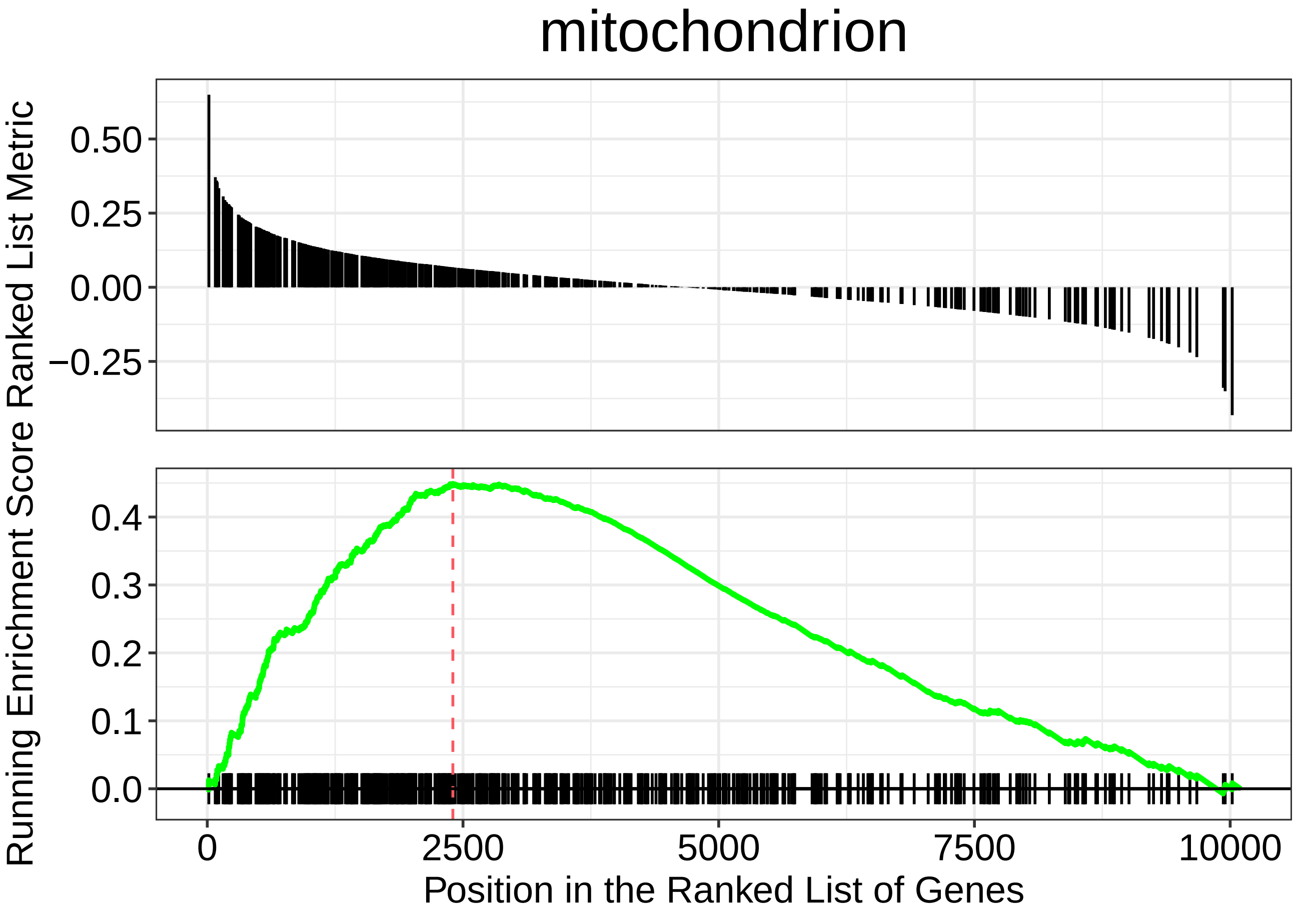
mitochondrion
Running Enrichment Score Ranked List Metric
0.50
0.25
0.00
−0.25
0.4
0.3
0.2
0.1
0.0
0
2500
5000
7500
10000
Position in the Ranked List of Genes

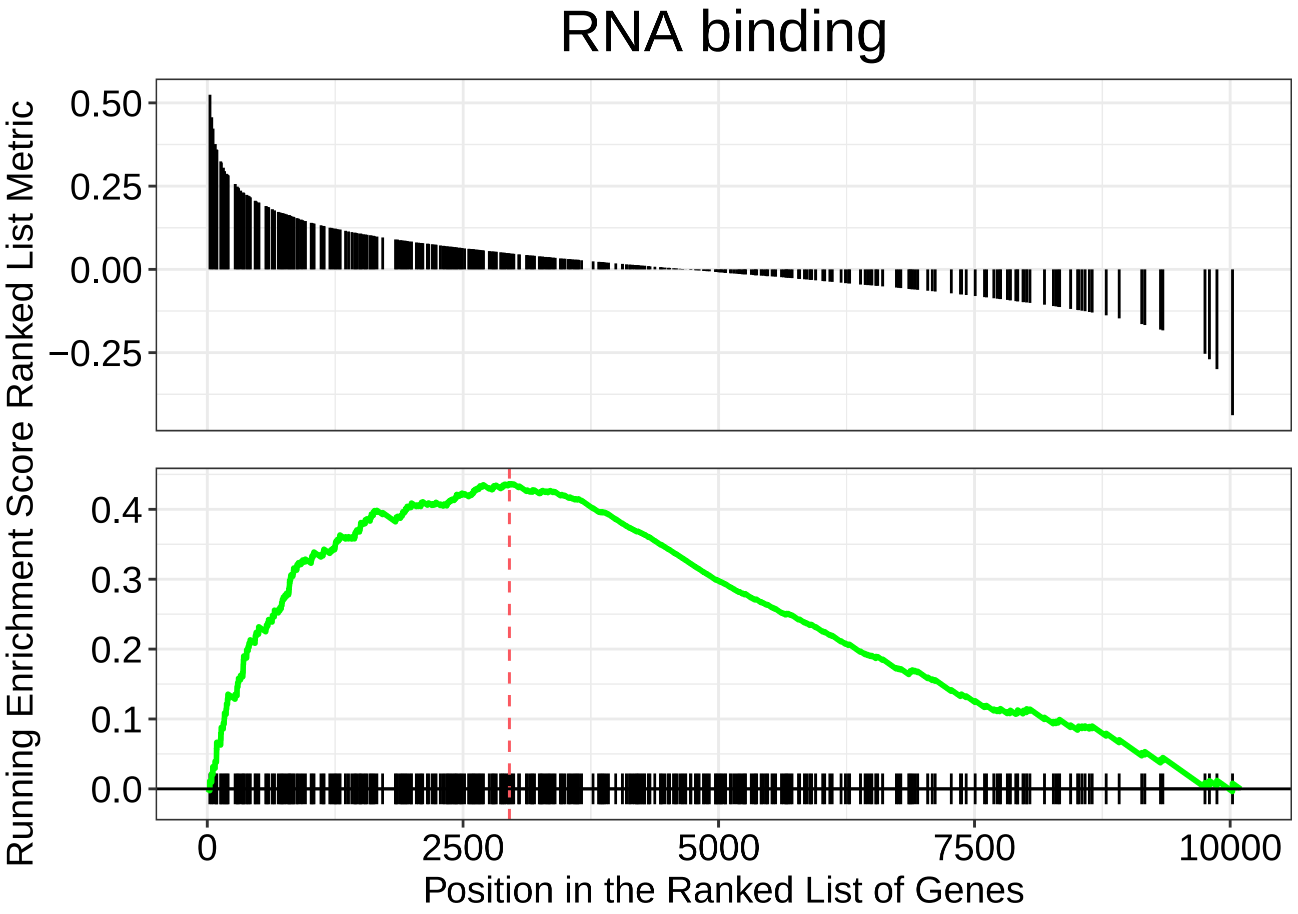
RNA binding
Running Enrichment Score Ranked List Metric
0.50
0.25
0.00
−0.25
0.4
0.3
0.2
0.1
0.0
0
2500
5000
7500
10000
Position in the Ranked List of Genes

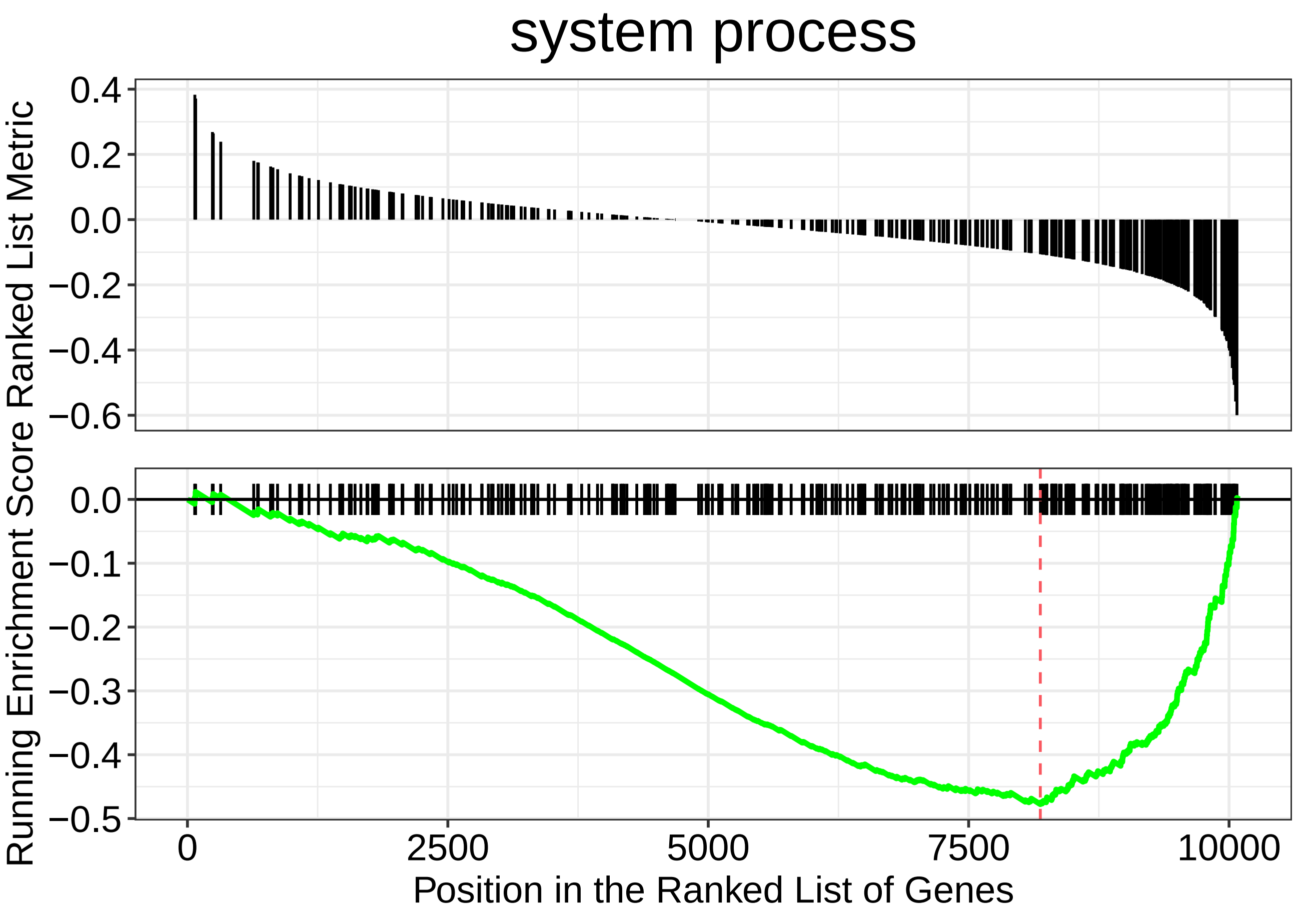

system process
Running Enrichment Score Ranked List Metric
0.4
0.2
0.0
−0.2
−0.4
−0.6
0.0
−0.1
−0.2
−0.3
−0.4
−0.5
0
2500
5000
7500
10000
Position in the Ranked List of Genes

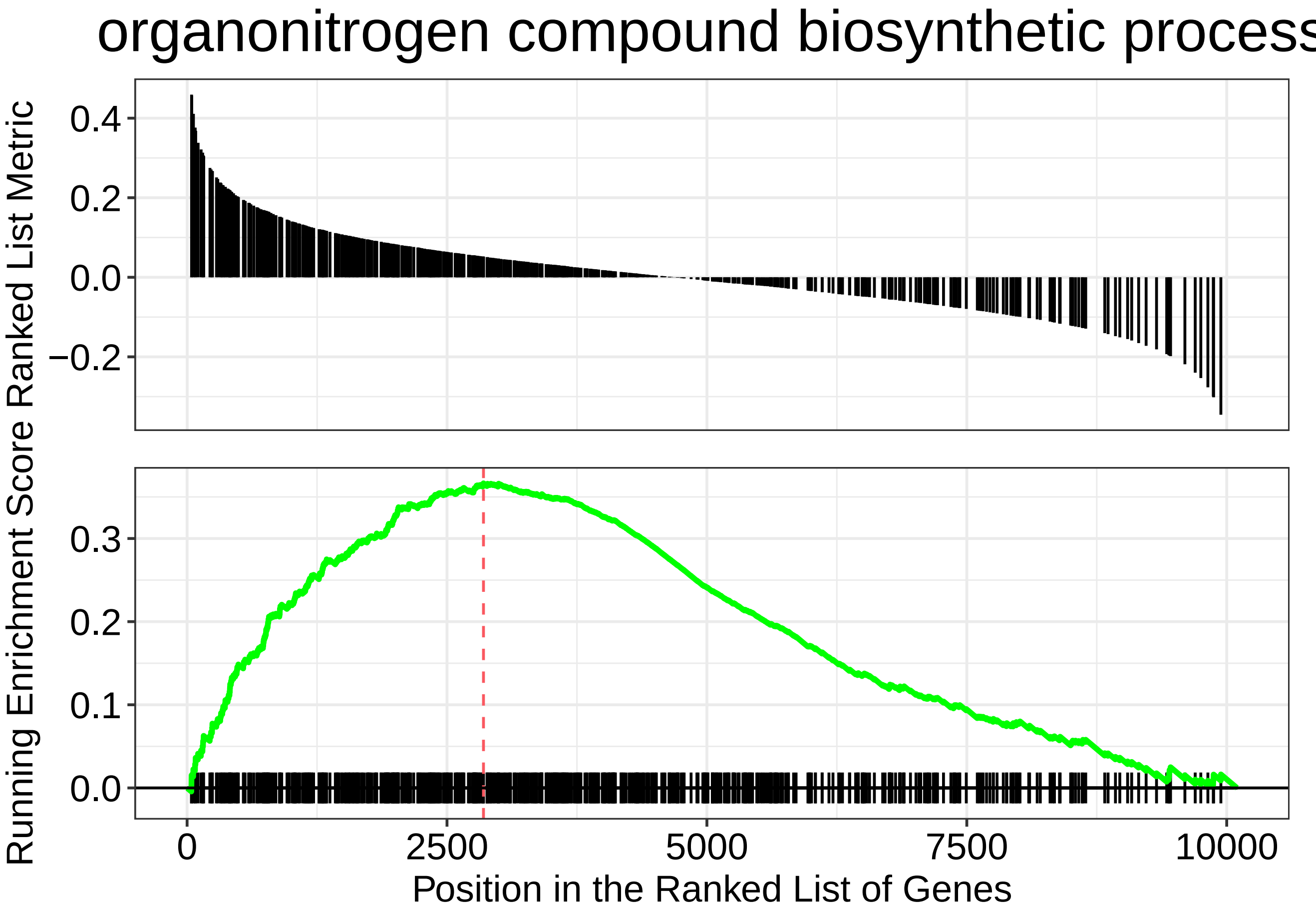
organonitrogen compound biosynthetic process
Ranked List Metric
0.4
0.2
0.0
−0.2
Running Enrichment Score
0.3
0.2
0.1
0.0
0
2500
5000
7500
10000
Position in the Ranked List of Genes

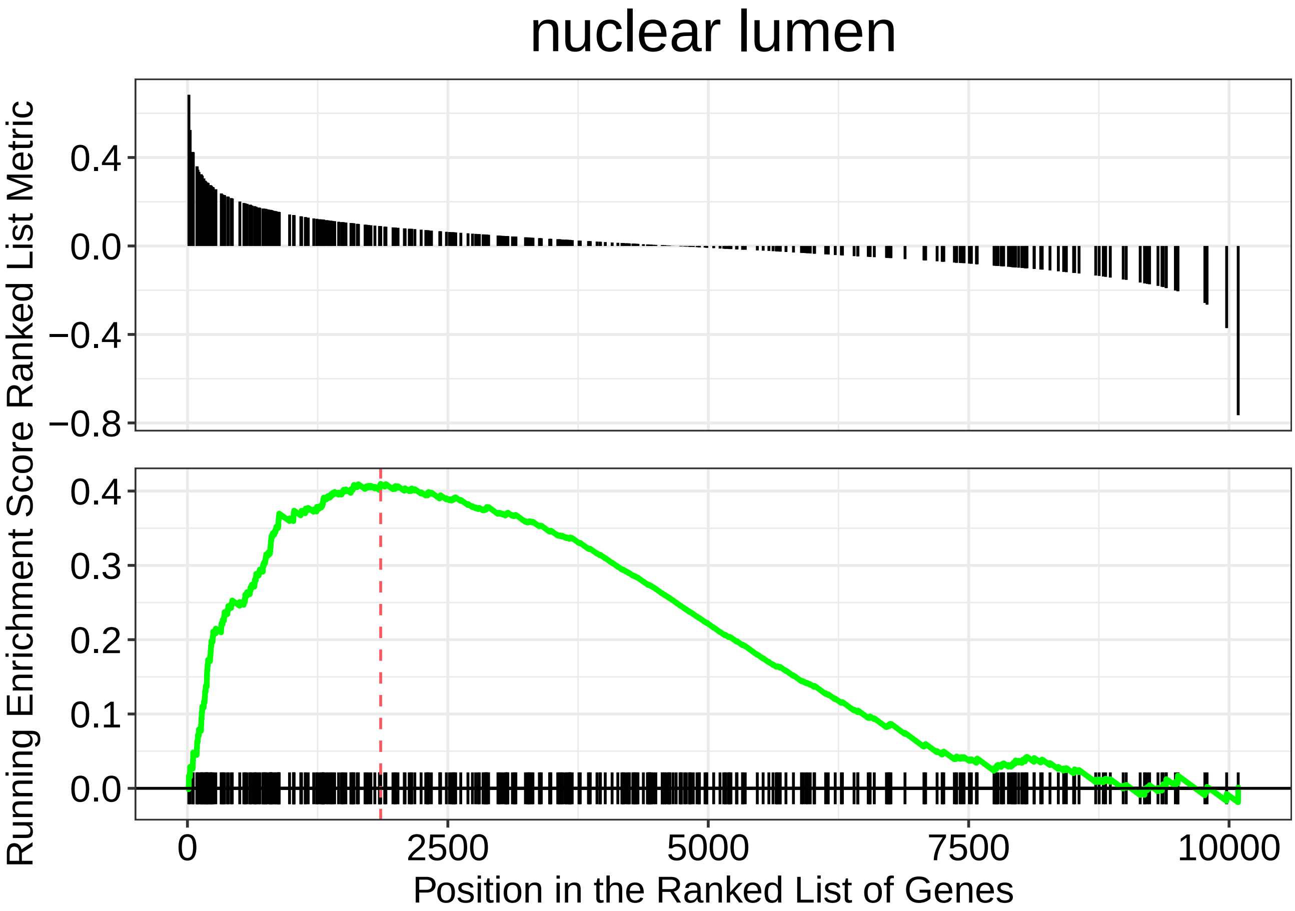
nuclear lumen
Running Enrichment Score Ranked List Metric
0.4
0.0
−0.4
−0.8
0.4
0.3
0.2
0.1
0.0
0
2500
5000
7500
10000
Position in the Ranked List of Genes

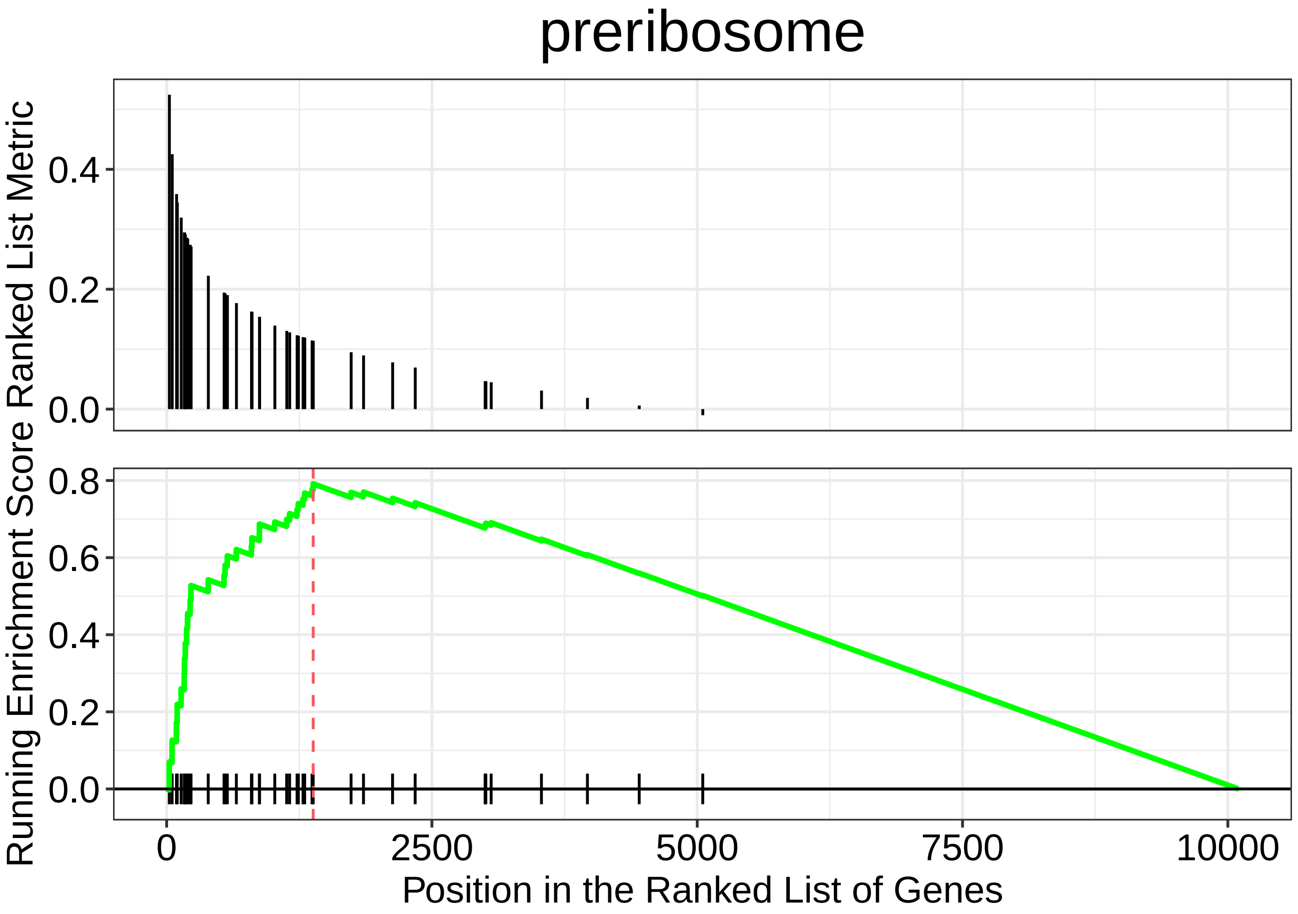

preriboseome

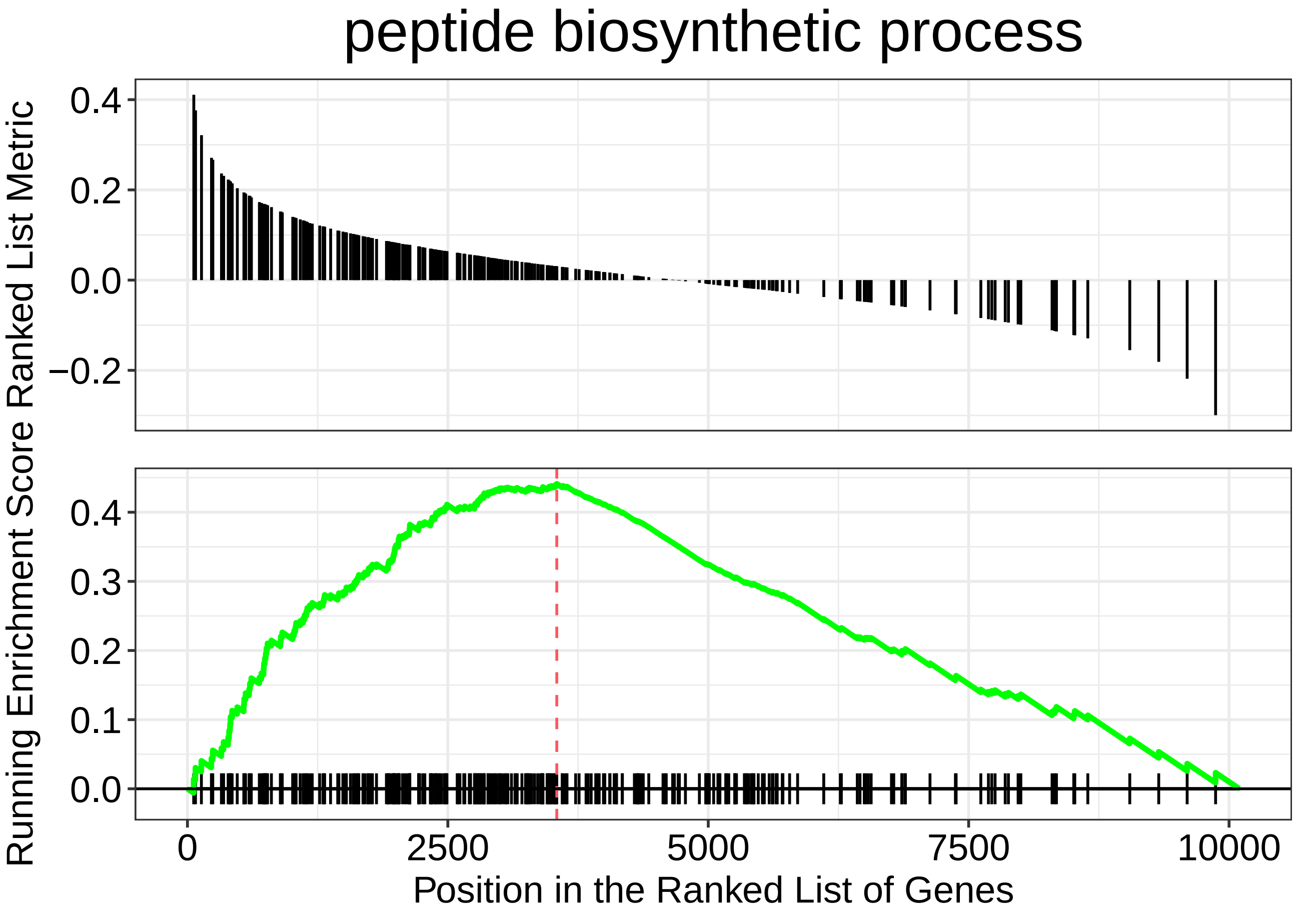

peptide biosynthetic process
Running Enrichment Score Ranked List Metric
0.4
0.2
0.0
−0.2
0.4
0.3
0.2
0.1
0.0
0
2500
5000
7500
10000
Position in the Ranked List of Genes

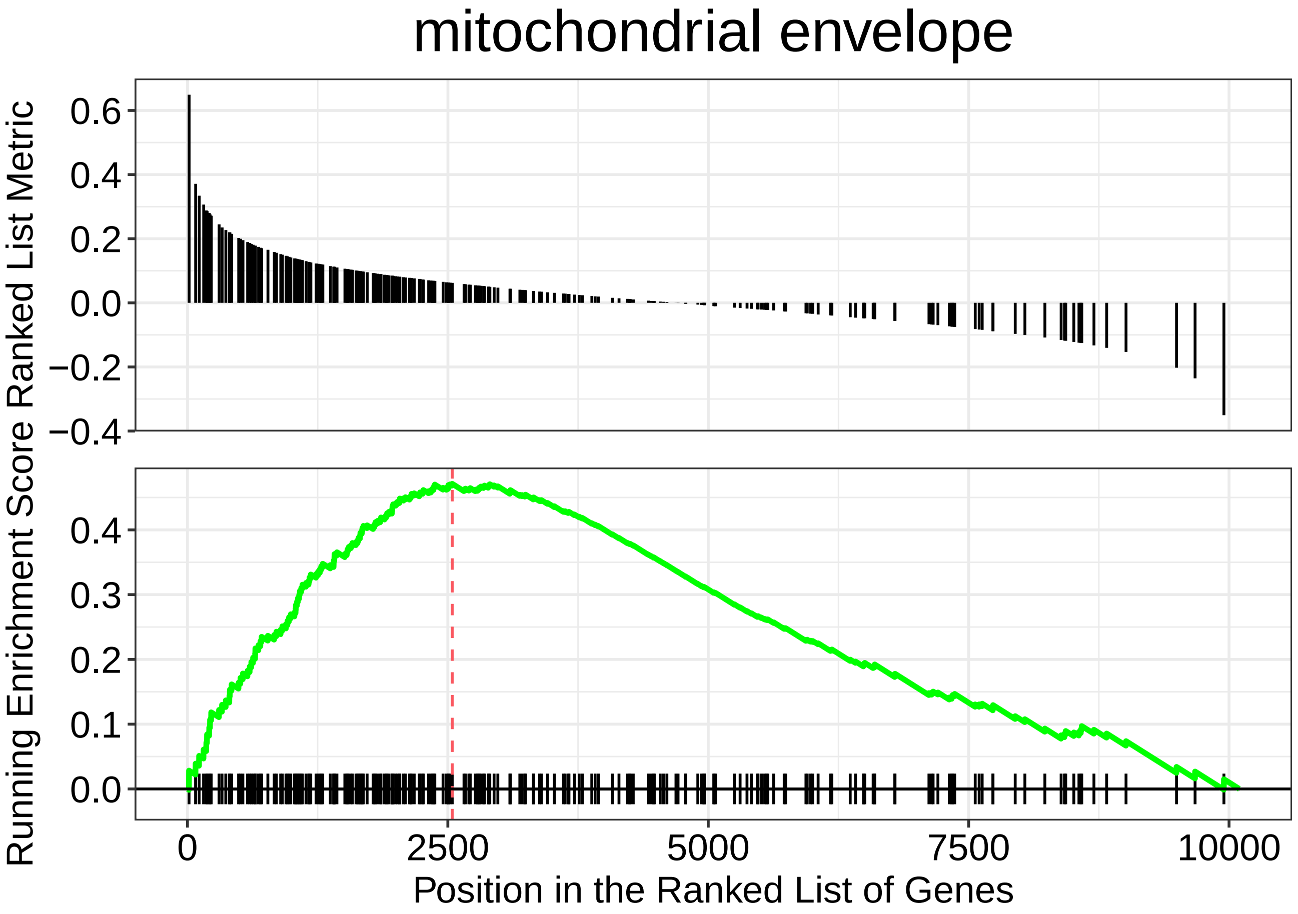
mitochondrial envelope
Ranked List Metric
0.6
0.4
0.2
0.0
−0.2
−0.4
Running Enrichment Score
0.4
0.3
0.2
0.1
0.0
0
2500
5000
7500
10000
Position in the Ranked List of Genes
