## Supplementary material for "The small molecule ML233 is a direct inhibitor of tyrosinase function": TableS2

TABLE 2

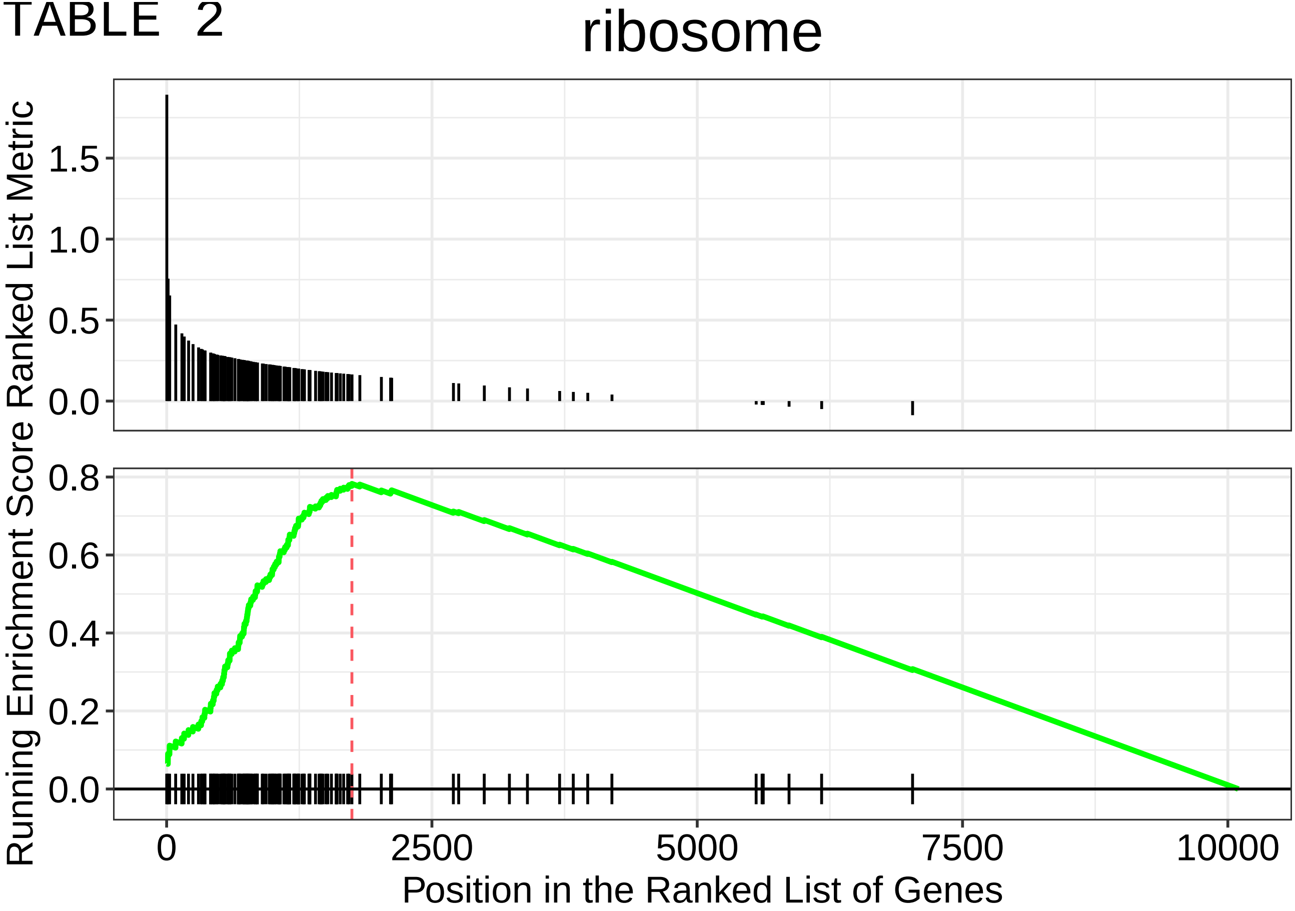
ribosome
Running Enrichment Score Ranked List Metric
1.5
1.0
0.5
0.0
0.8
0.6
0.4
0.2
0.0
0
2500
5000
7500
10000
Position in the Ranked List of Genes

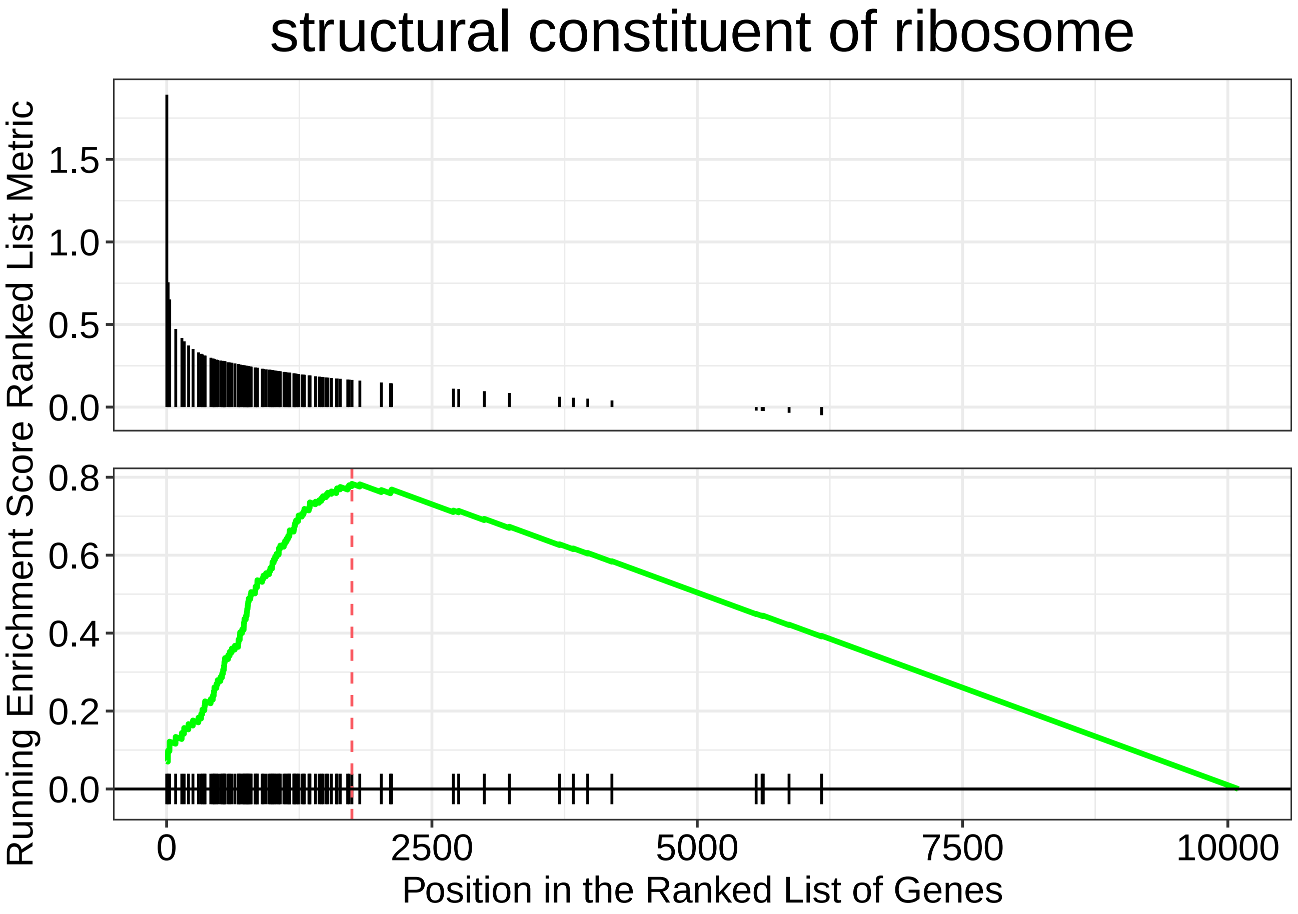

structural constituent of ribosome
Ranked List Metric
1.5
1.0
0.5
0.0
Running Enrichment Score
0.8
0.6
0.4
0.2
0.0
0
2500
5000
7500
10000
Position in the Ranked List of Genes

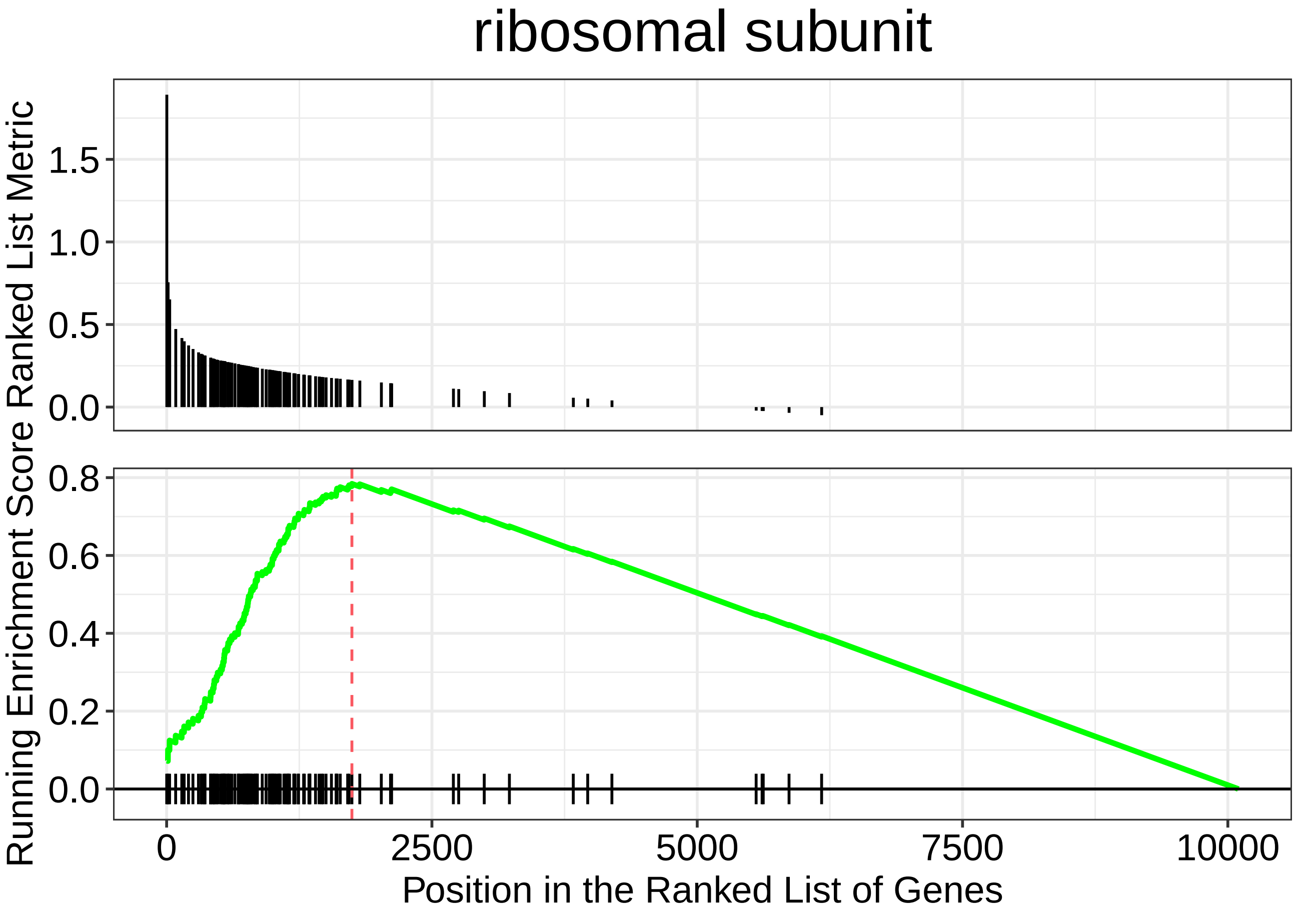

ribosomal subunit
Ranked List Metric
1.5
1.0
0.5
0.0
Running Enrichment Score
0.8
0.6
0.4
0.2
0.0
0
2500
5000
7500
10000
Position in the Ranked List of Genes

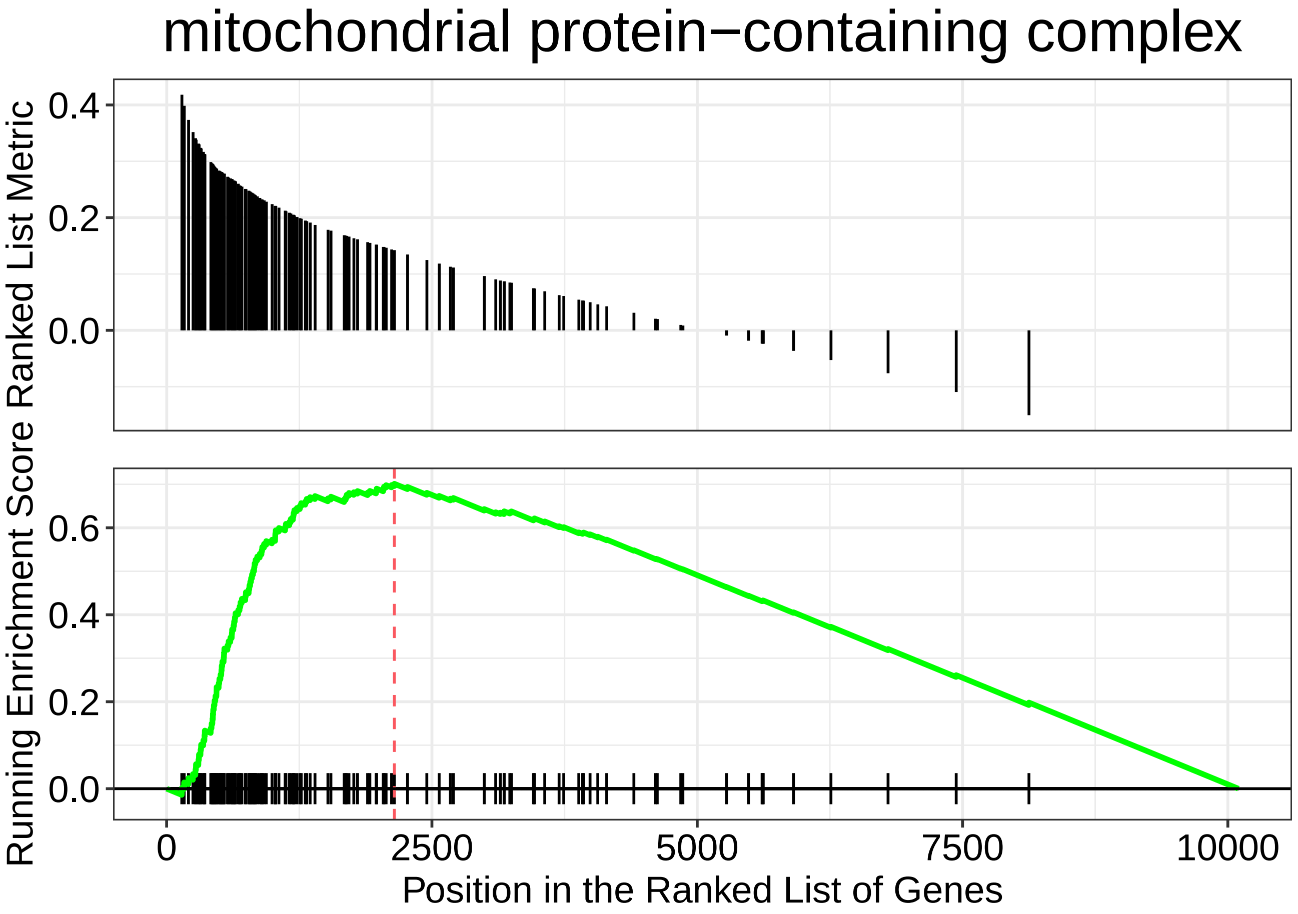

mitochondrial protein−containing complex
Running Enrichment Score Ranked List Metric
0.4
0.2
0.0
0.6
0.4
0.2
0.0
0
2500
5000
7500
10000
Position in the Ranked List of Genes

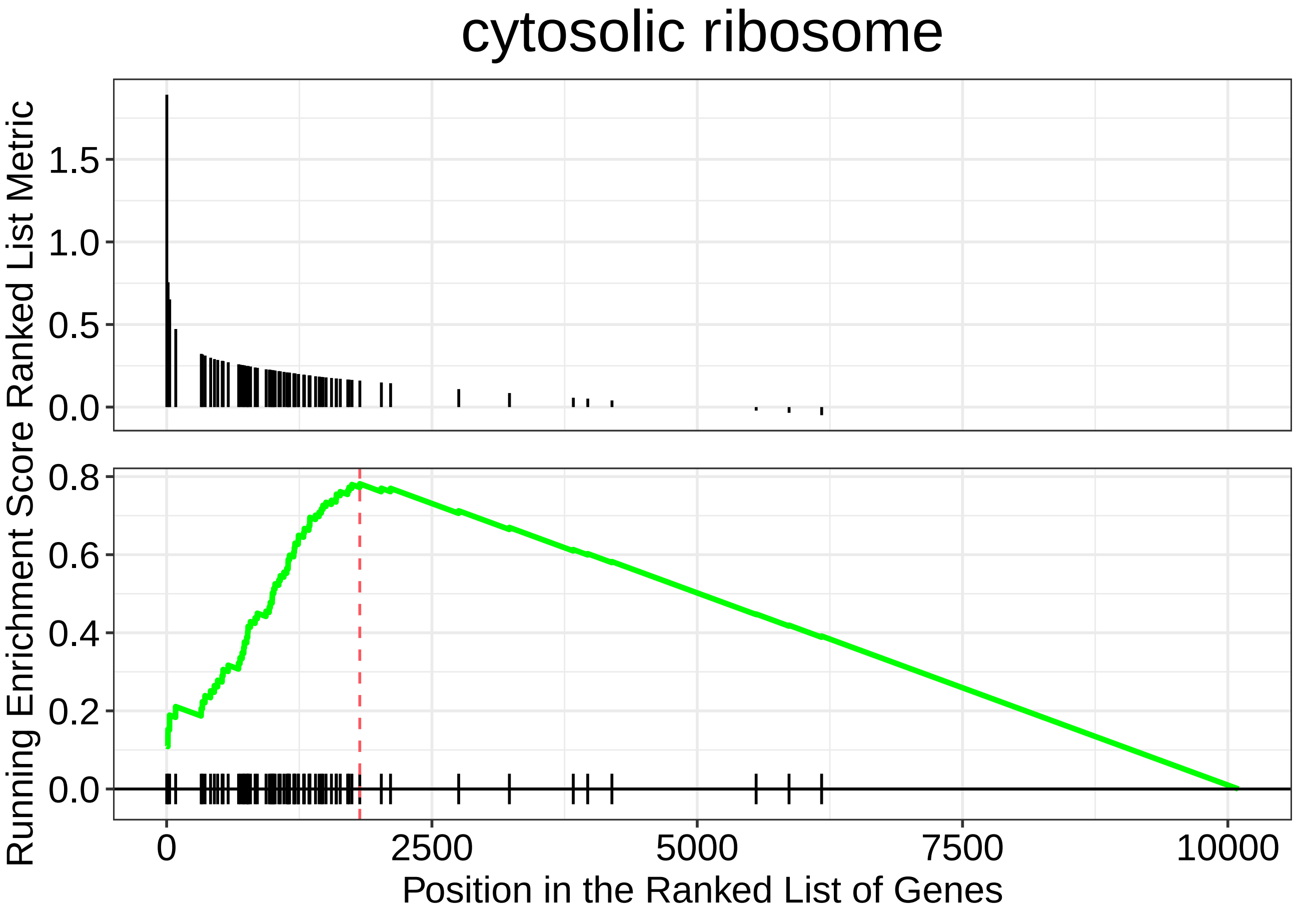
cytosolic ribosome
Ranked List Metric
1.5
1.0
0.5
0.0
Running Enrichment Score
0.8
0.6
0.4
0.2
0.0
0
2500
5000
7500
10000
Position in the Ranked List of Genes

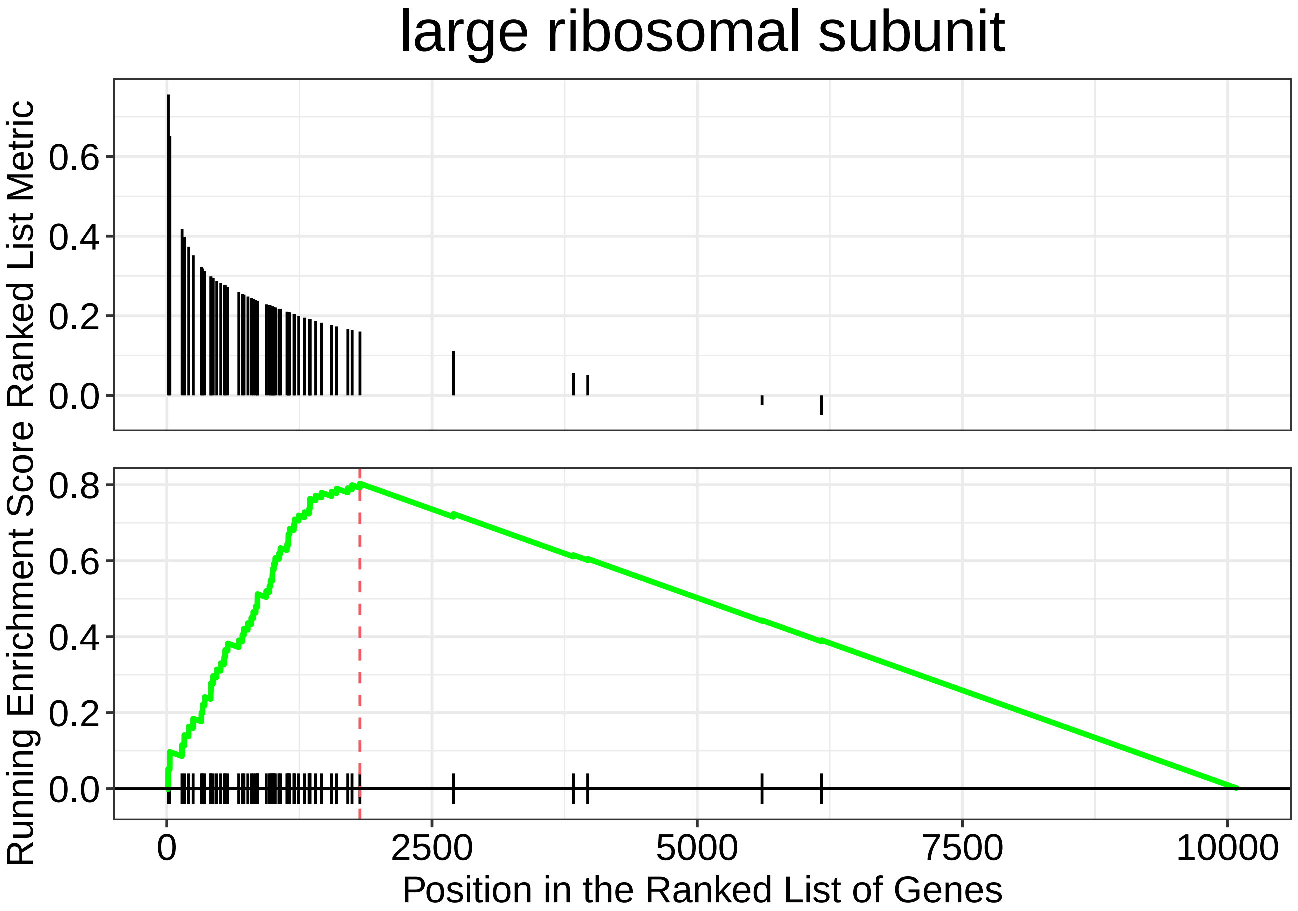
large ribosomal subunit
Ranked List Metric
Running Enrichment Score
0.0
0.2
0.4
0.6
0.8
0
2500
5000
7500
10000
Position in the Ranked List of Genes

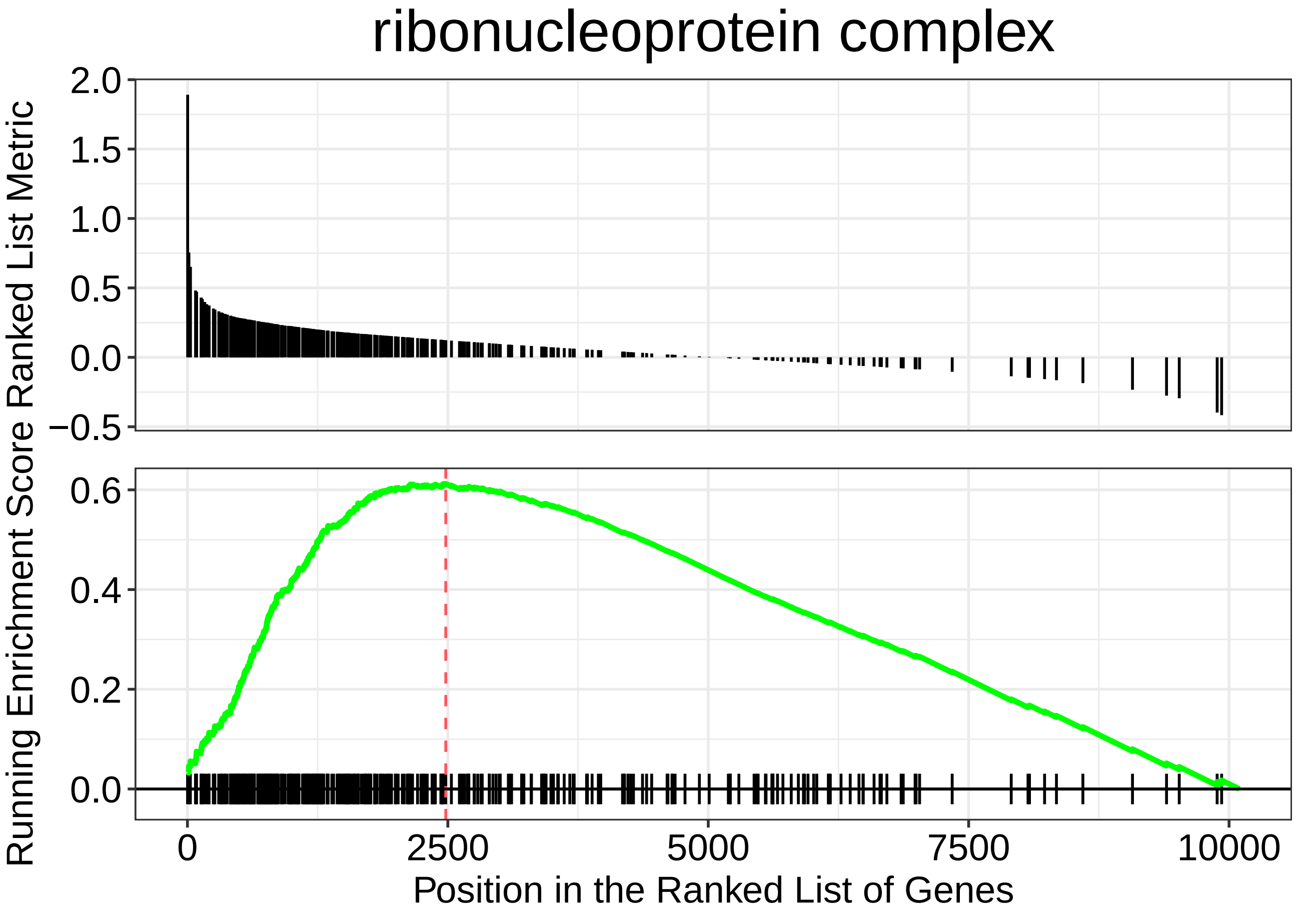
ribonucleoprotein complex
Ranked List Metric
2.0
1.5
1.0
0.5
0.0
−0.5
Running Enrichment Score
0.6
0.4
0.2
0.0
0
2500
5000
7500
10000
Position in the Ranked List of Genes

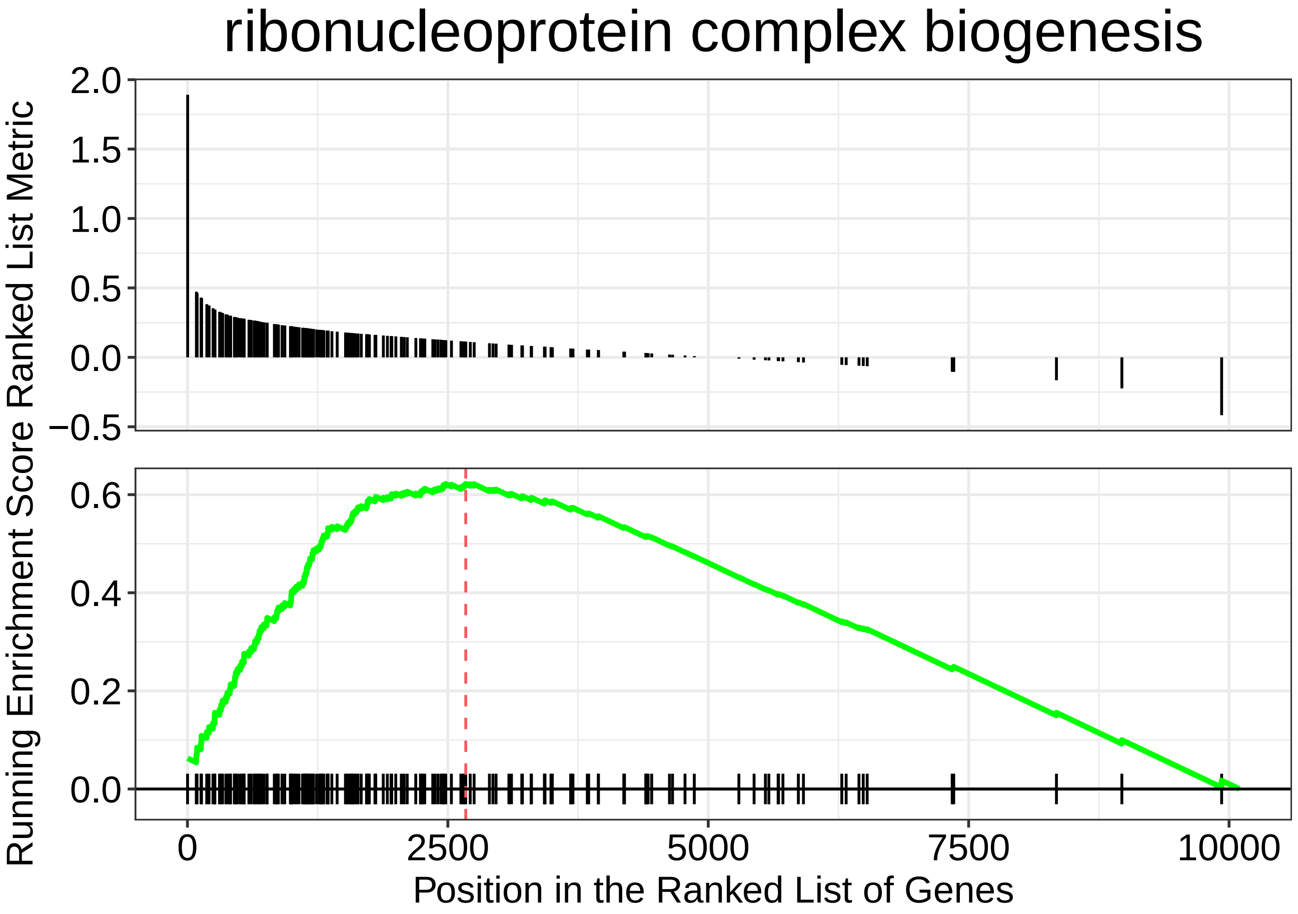

ribonucleoprotein complex biogenesis
Ranked List Metric
2.0
1.5
1.0
0.5
0.0
−0.5
Running Enrichment Score
0.6
0.4
0.2
0.0
0
2500
5000
7500
10000
Position in the Ranked List of Genes

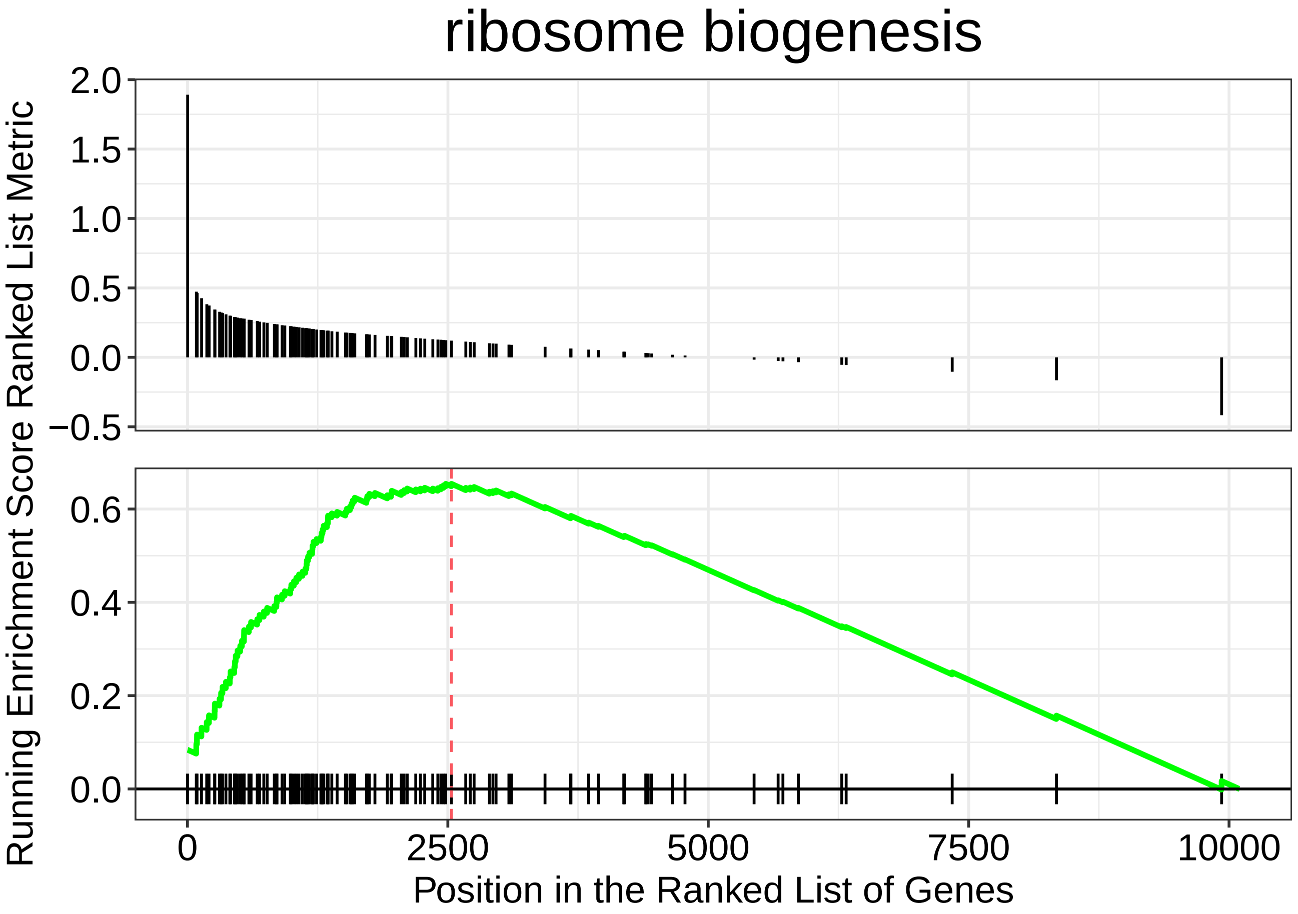

ribosome biogenesis
Ranked List Metric
2.0
1.5
1.0
0.5
0.0
−0.5
Running Enrichment Score
0.6
0.4
0.2
0.0
0
2500
5000
7500
10000
Position in the Ranked List of Genes

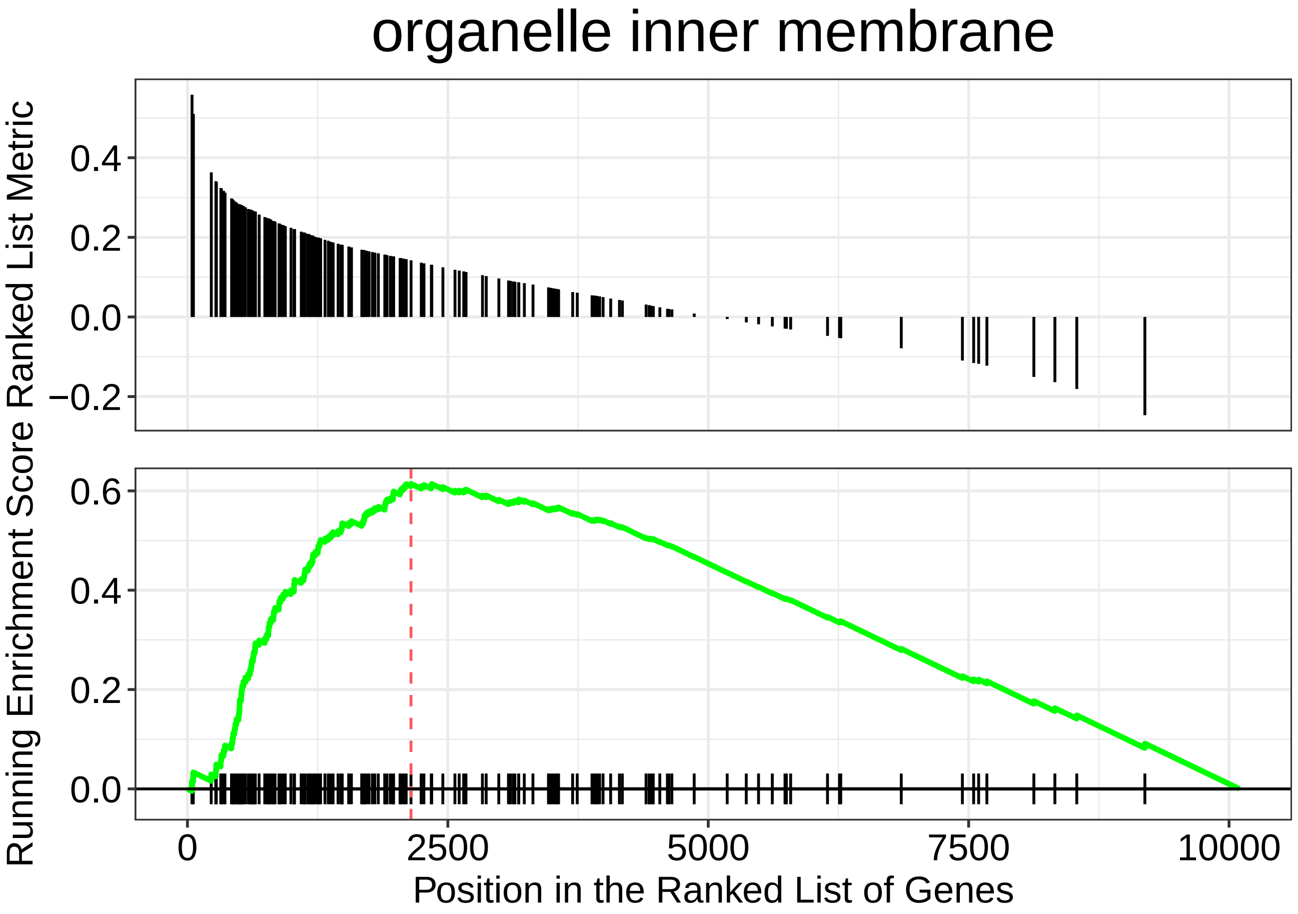
organelle inner membrane
Running Enrichment Score Ranked List Metric
0.4
0.2
0.0
−0.2
0.6
0.4
0.2
0.0
0
2500
5000
7500
10000
Position in the Ranked List of Genes

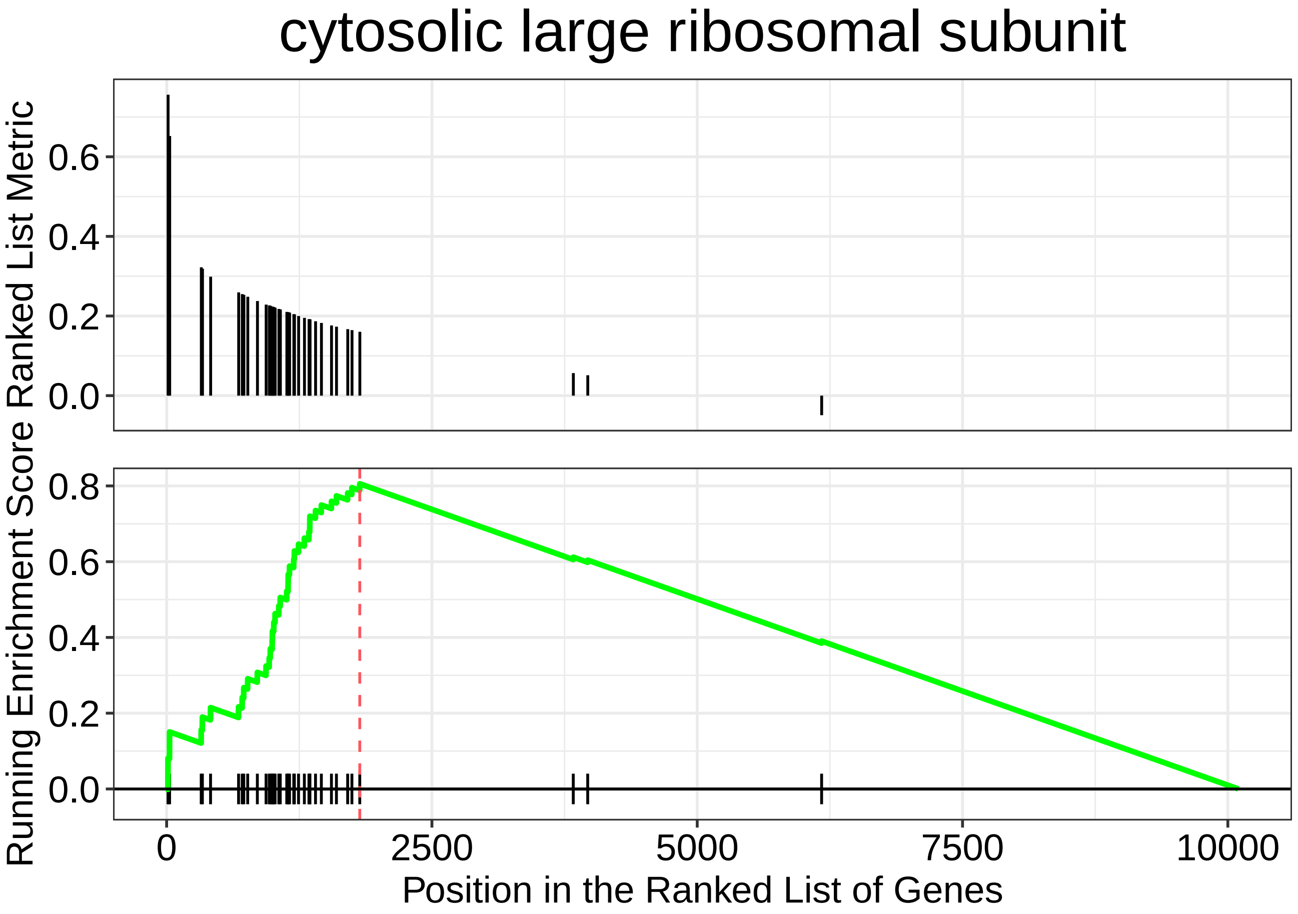
cytosolic large ribosomal subunit
Ranked List Metric
0.6
0.4
0.2
0.0
Running Enrichment Score
0.8
0.6
0.4
0.2
0.0
0
2500
5000
7500
10000
Position in the Ranked List of Genes

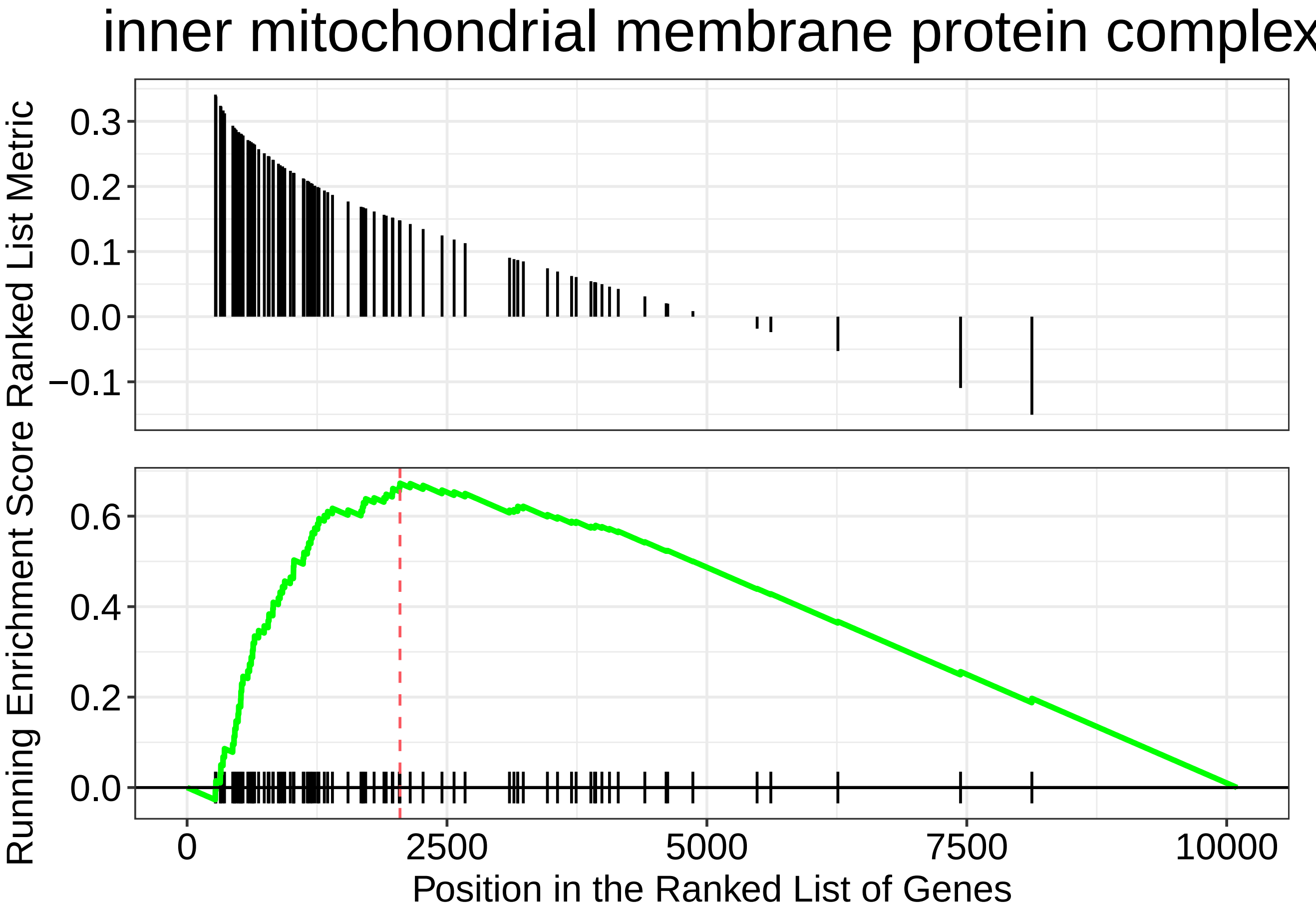

inner mitochondrial membrane protein complex
Running Enrichment Score Ranked List Metric
0.3
0.2
0.1
0.0
−0.1
0.6
0.4
0.2
0.0
0
2500
5000
7500
10000
Position in the Ranked List of Genes

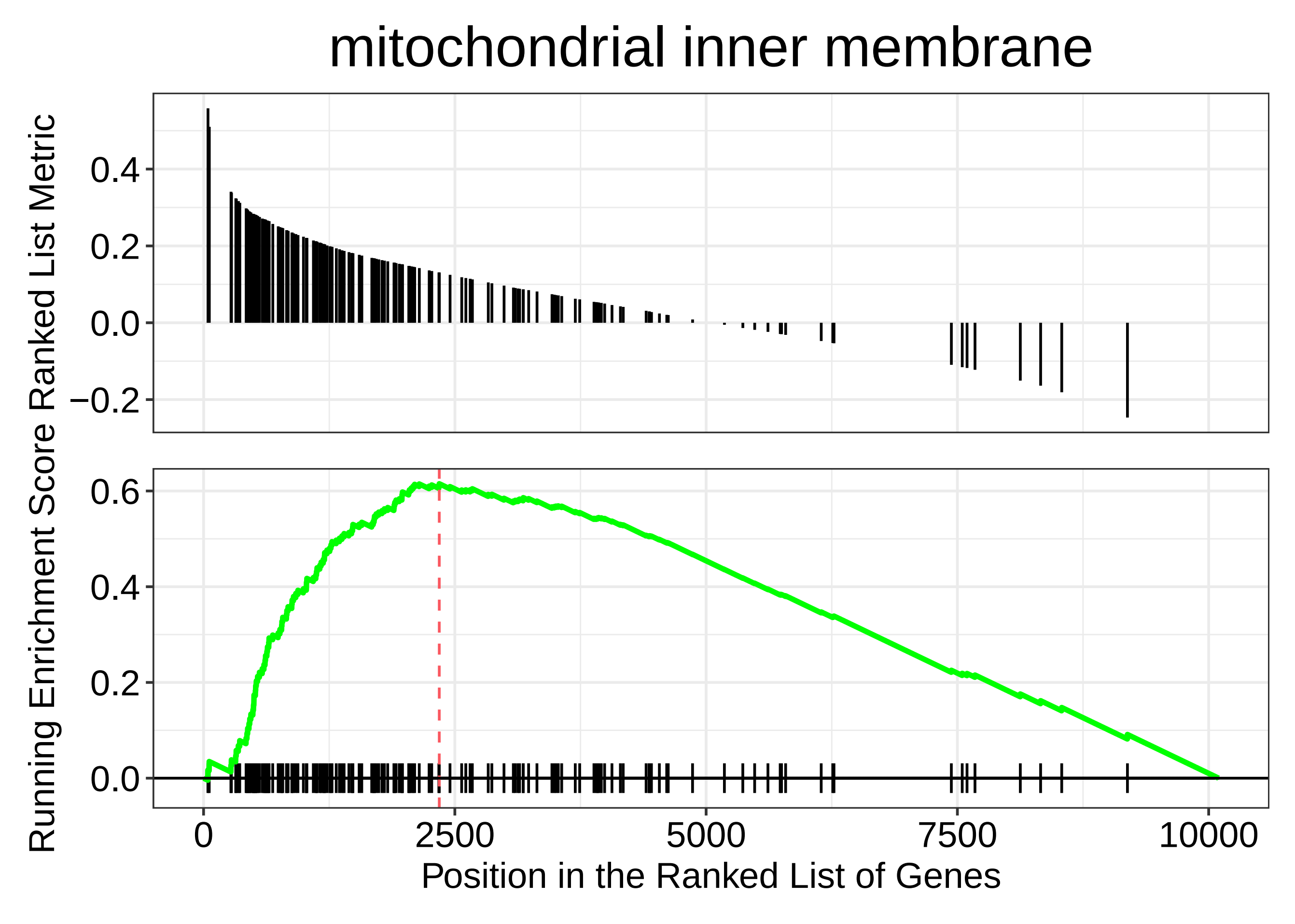

mitochondrial inner membrane
Running Enrichment Score Ranked List Metric
0.4
0.2
0.0
−0.2
0.6
0.4
0.2
0.0
0
2500
5000
7500
10000
Position in the Ranked List of Genes

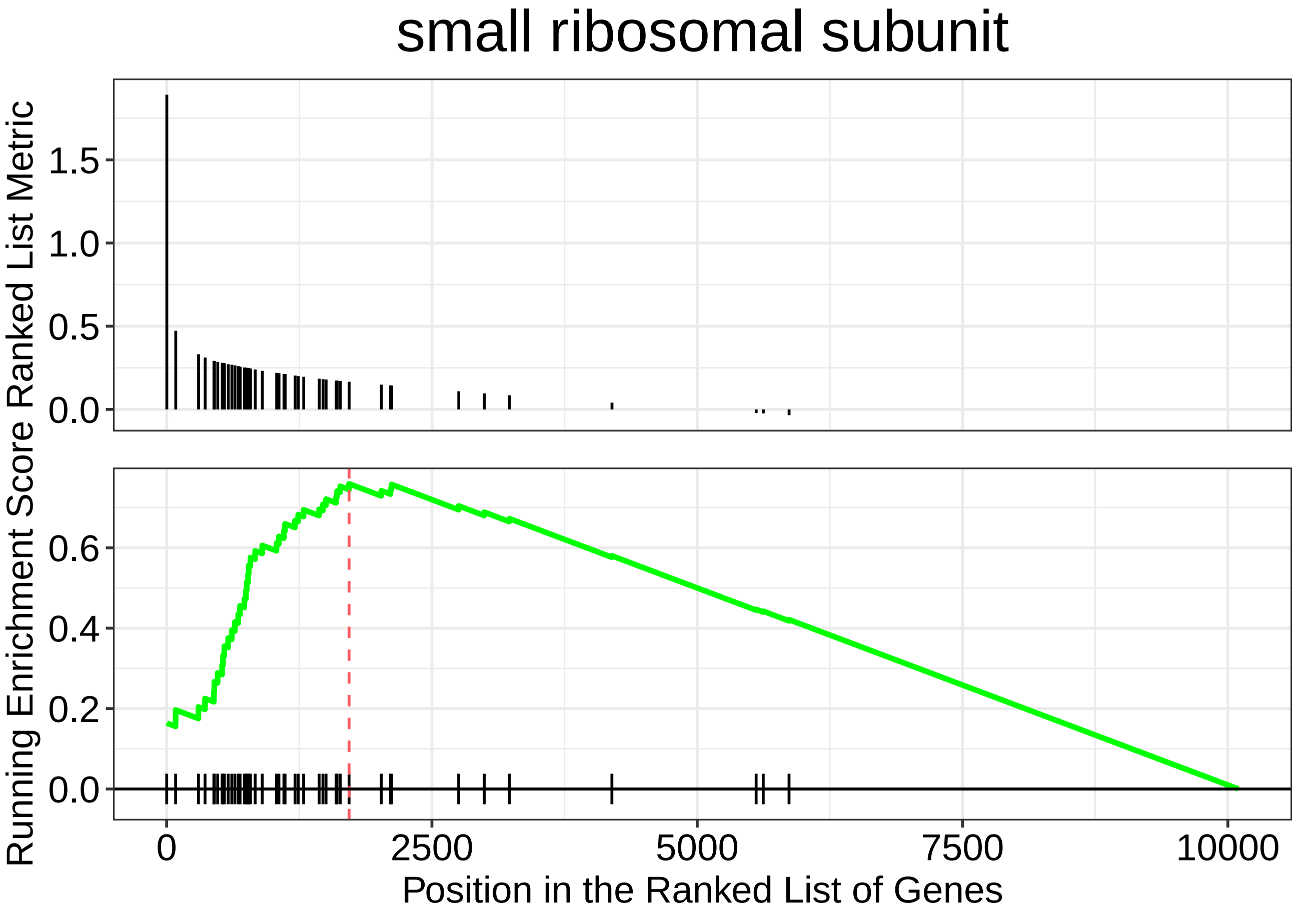

small ribosomal subunit
Ranked List Metric
1.5
1.0
0.5
0.0
Running Enrichment Score
0.6
0.4
0.2
0.0
0
2500
5000
7500
10000
Position in the Ranked List of Genes

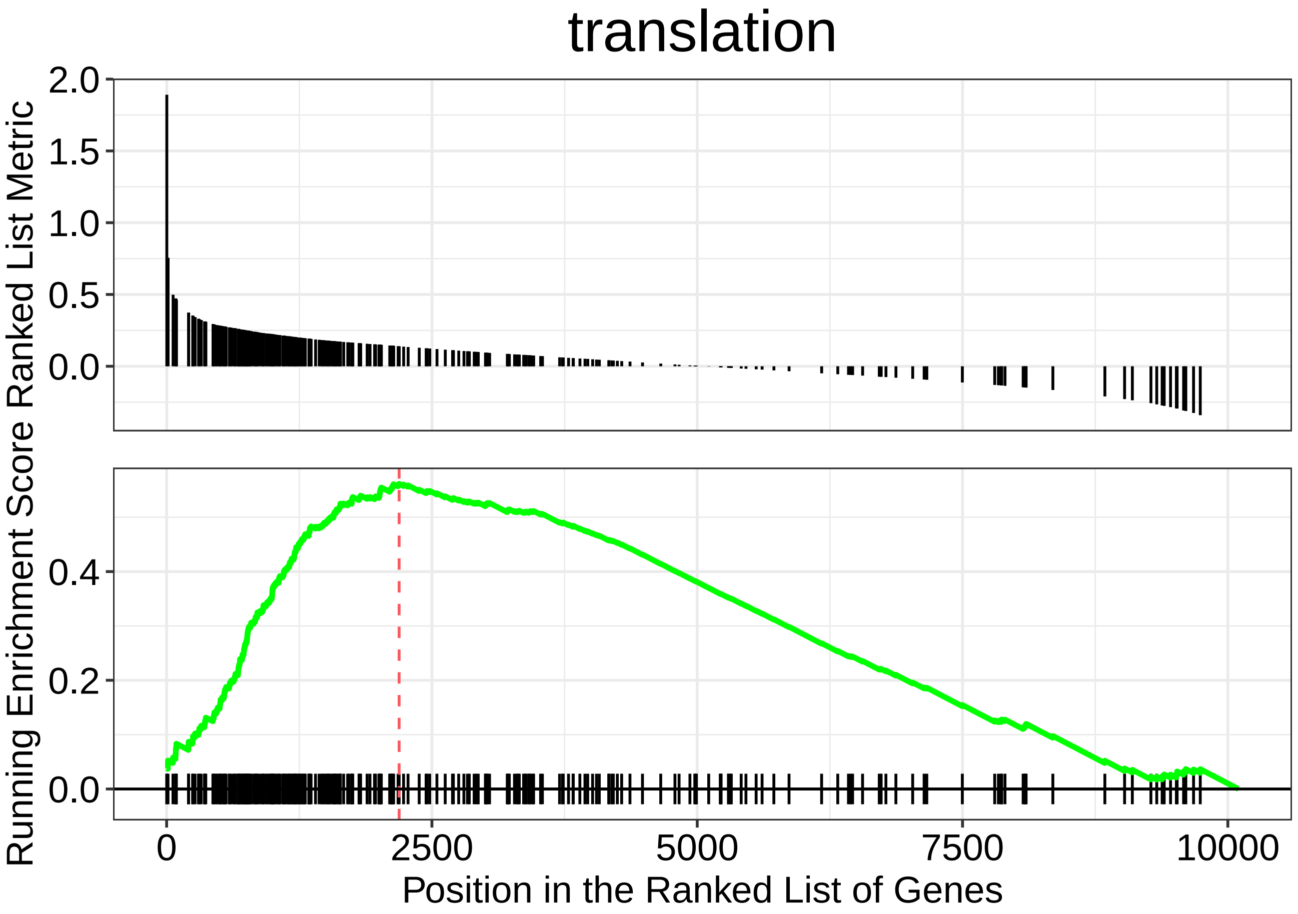

translation
Ranked List Metric
2.0
1.5
1.0
0.5
0.0
Running Enrichment Score
0.4
0.2
0.0
0
2500
5000
7500
10000
Position in the Ranked List of Genes

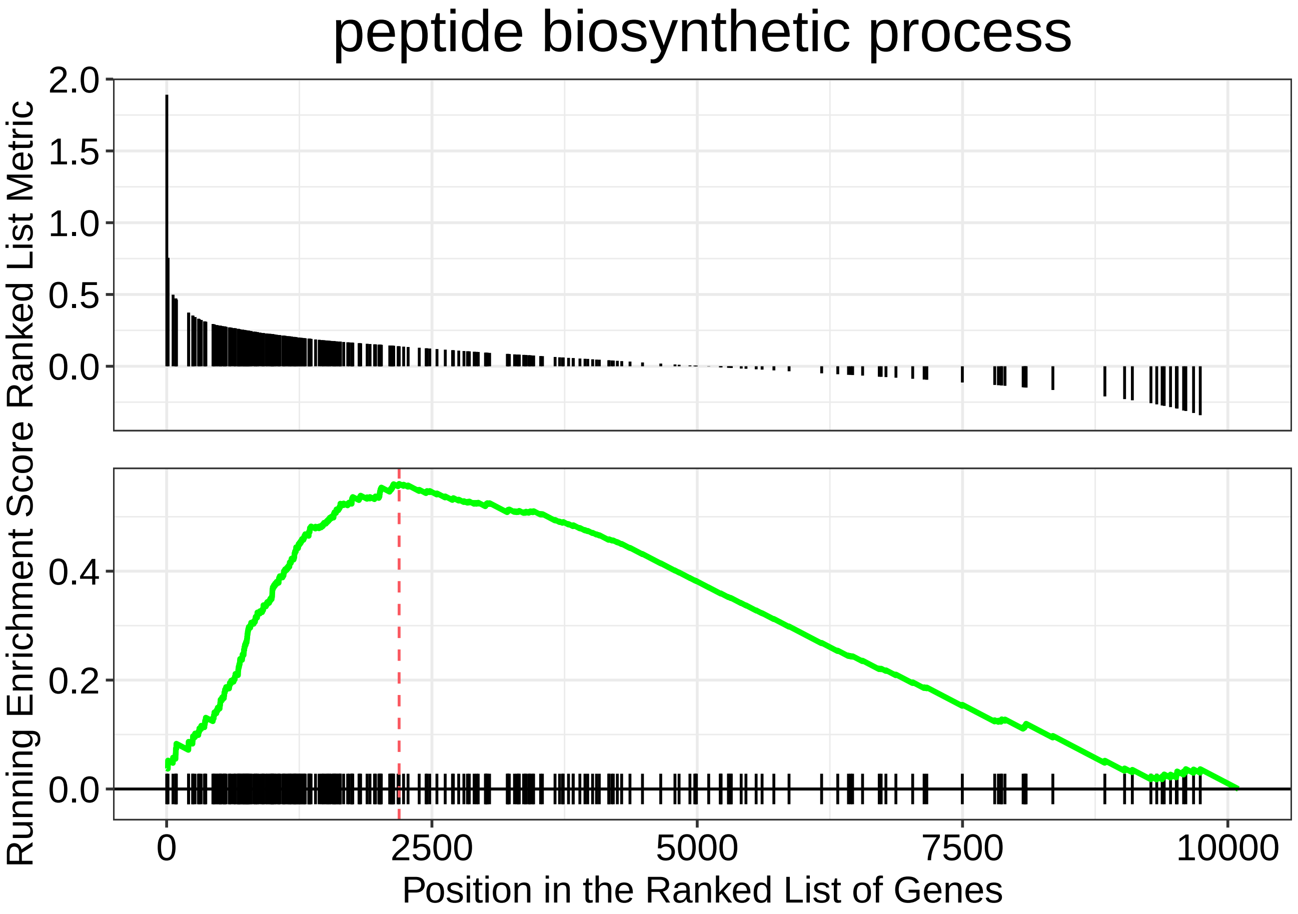

peptide biosynthetic process
Running Enrichment Score Ranked List Metric
2.0
1.5
1.0
0.5
0.0
0.4
0.2
0.0
0
2500
5000
7500
10000
Position in the Ranked List of Genes

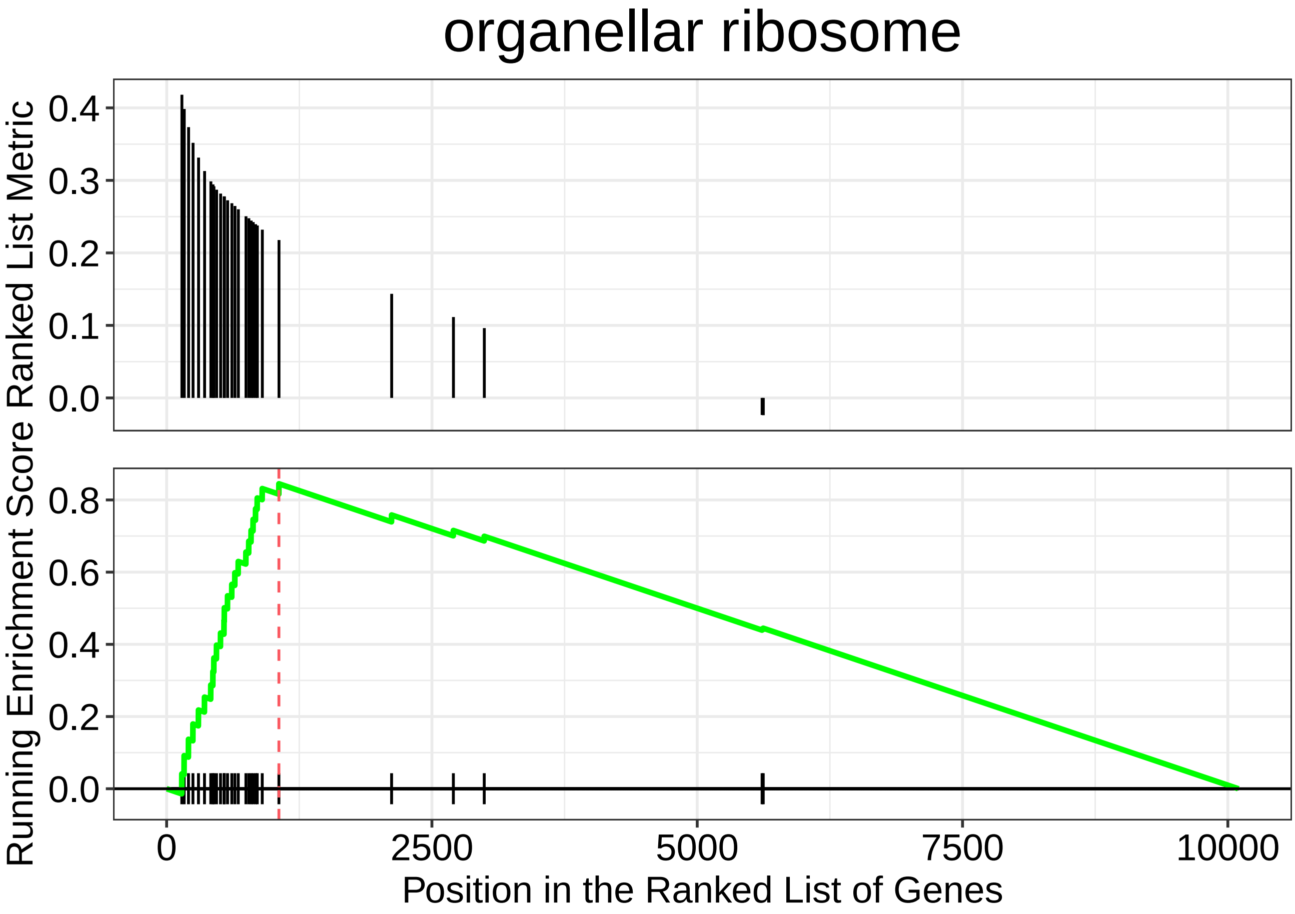
organellar ribosome
Running Enrichment Score Ranked List Metric
0.4
0.3
0.2
0.1
0.0
0.8
0.6
0.4
0.2
0.0
0
2500
5000
7500
10000
Position in the Ranked List of Genes

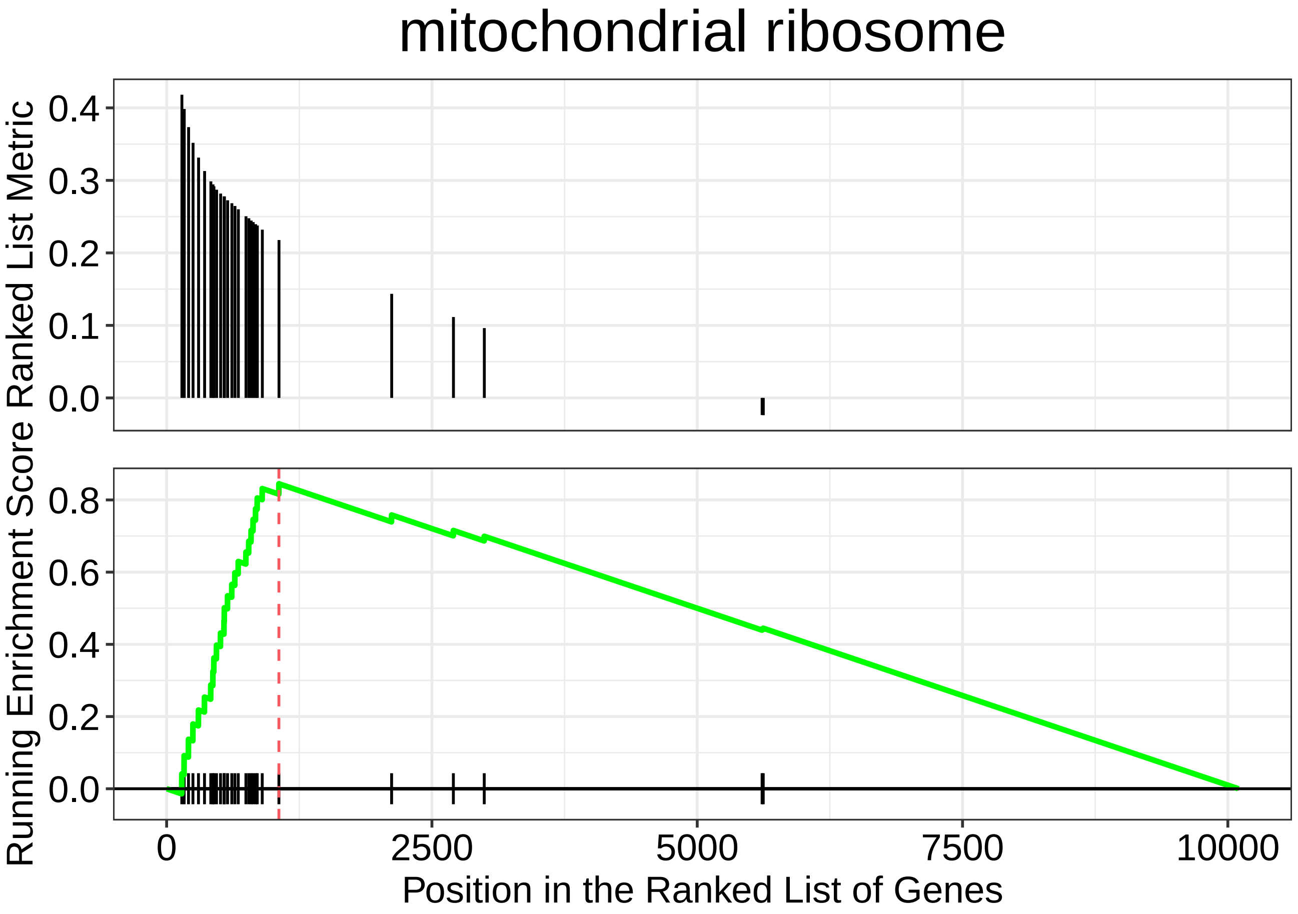

mitochondrial ribosome
Ranked List Metric
0.4
0.3
0.2
0.1
0.0
Running Enrichment Score
0.8
0.6
0.4
0.2
0.0
0
2500
5000
7500
10000
Position in the Ranked List of Genes

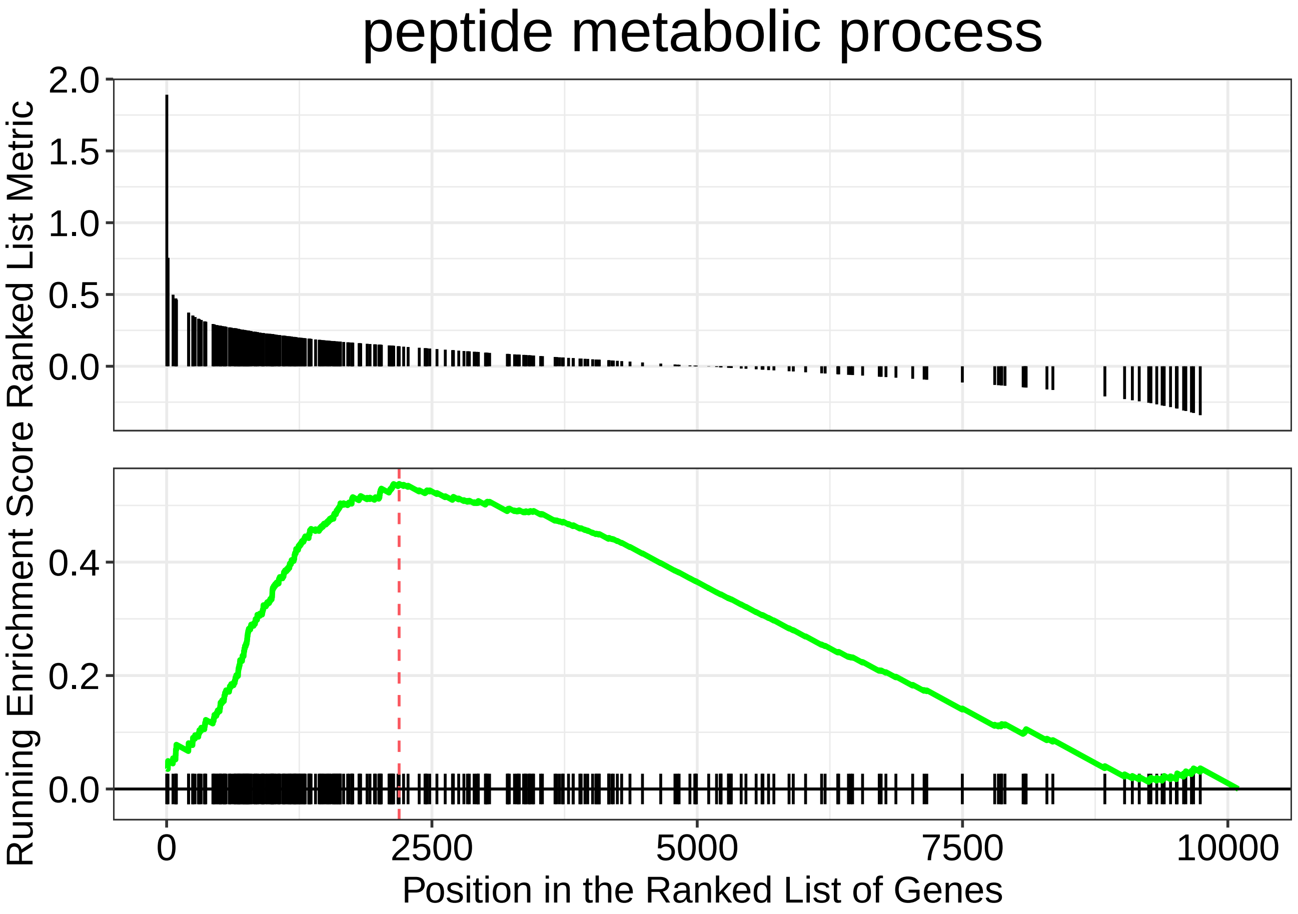
peptide metabolic process
Running Enrichment Score Ranked List Metric
2.0
1.5
1.0
0.5
0.0
0.4
0.2
0.0
0
2500
5000
7500
10000
Position in the Ranked List of Genes

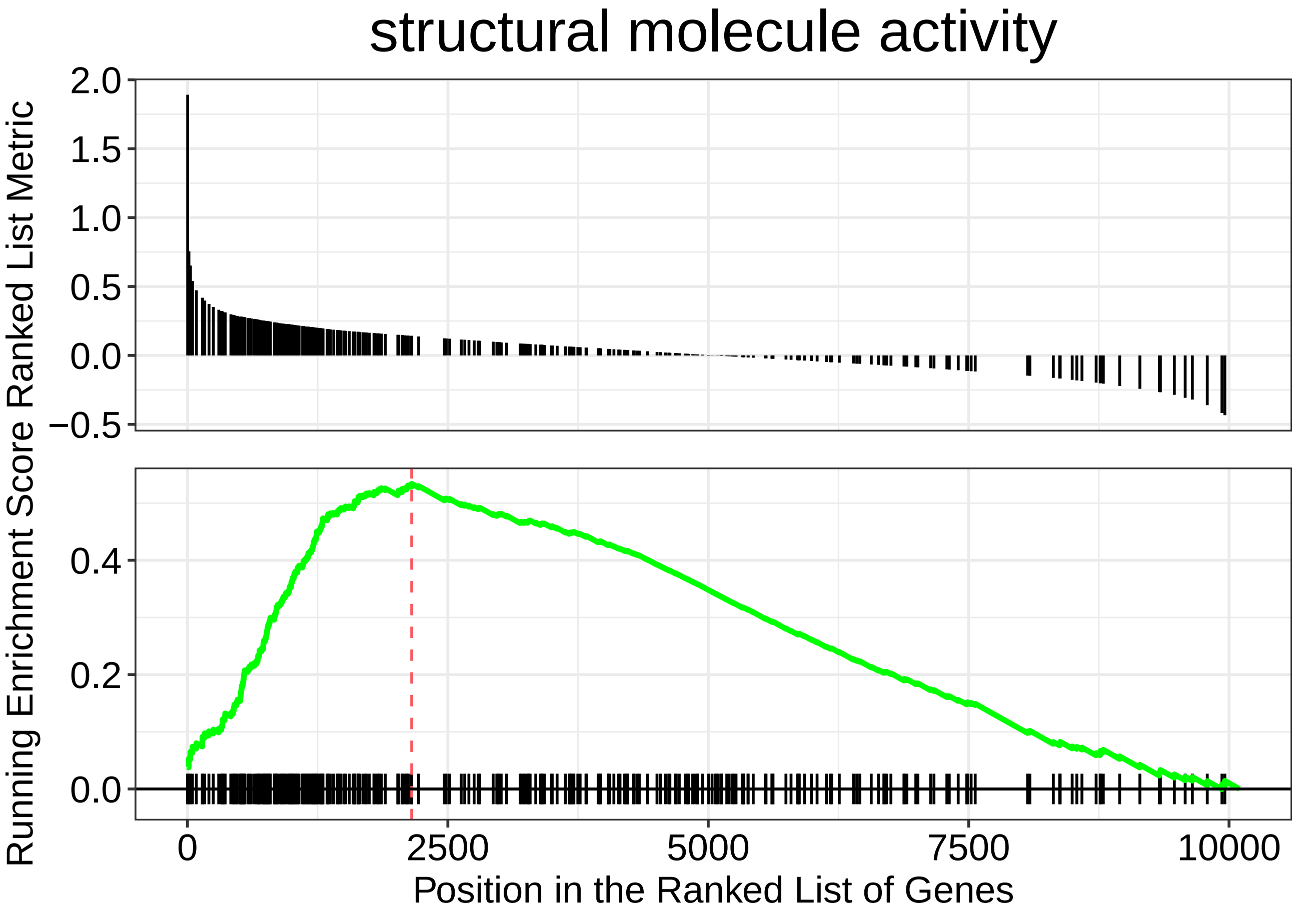

structural molecule activity
Running Enrichment Score Ranked List Metric
2.0
1.5
1.0
0.5
0.0
−0.5
0.4
0.2
0.0
0
2500
5000
7500
10000
Position in the Ranked List of Genes

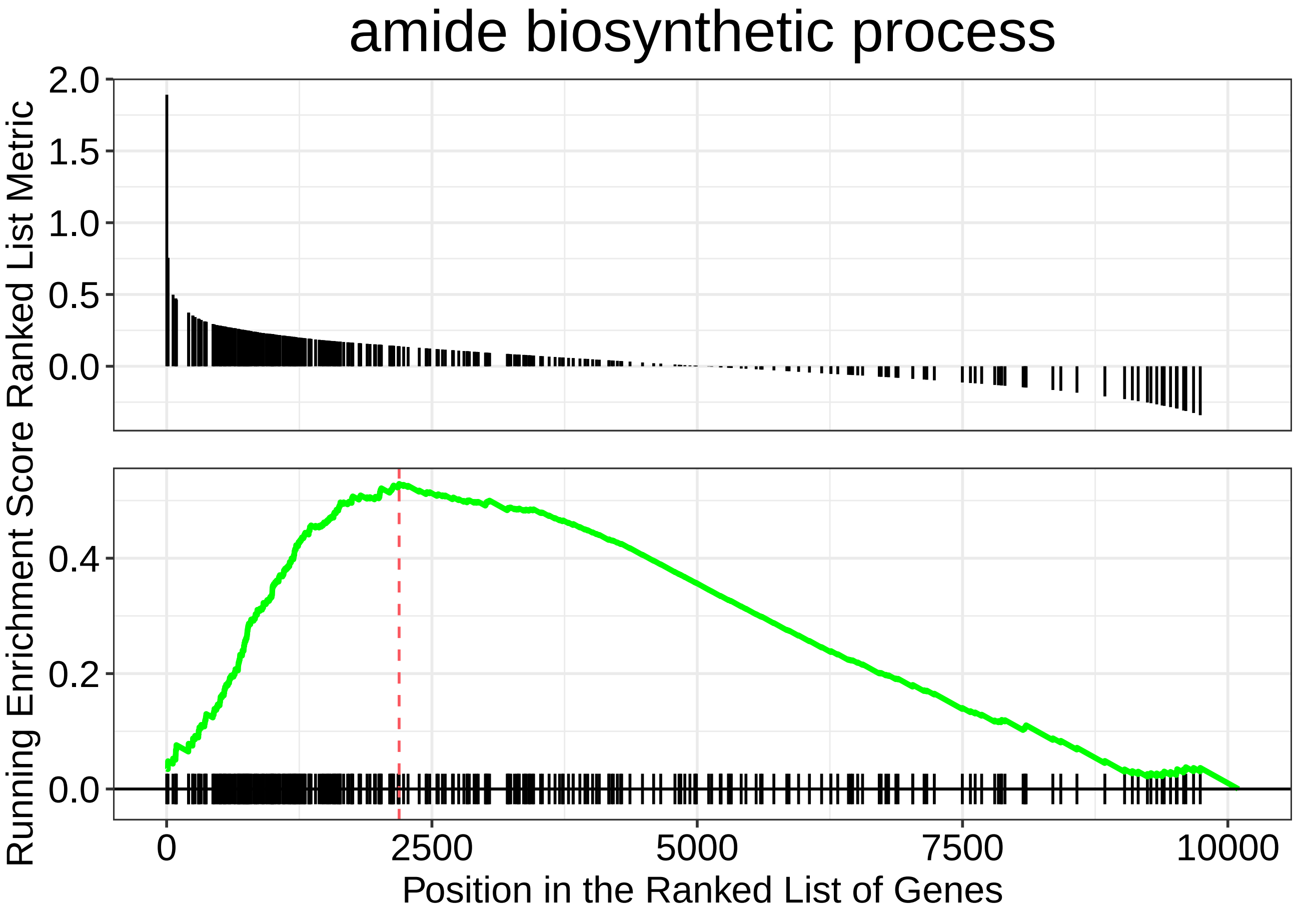

amide biosynthetic process
Ranked List Metric
2.0
1.5
1.0
0.5
0.0
Running Enrichment Score
0.4
0.2
0.0
0
2500
5000
7500
10000
Position in the Ranked List of Genes

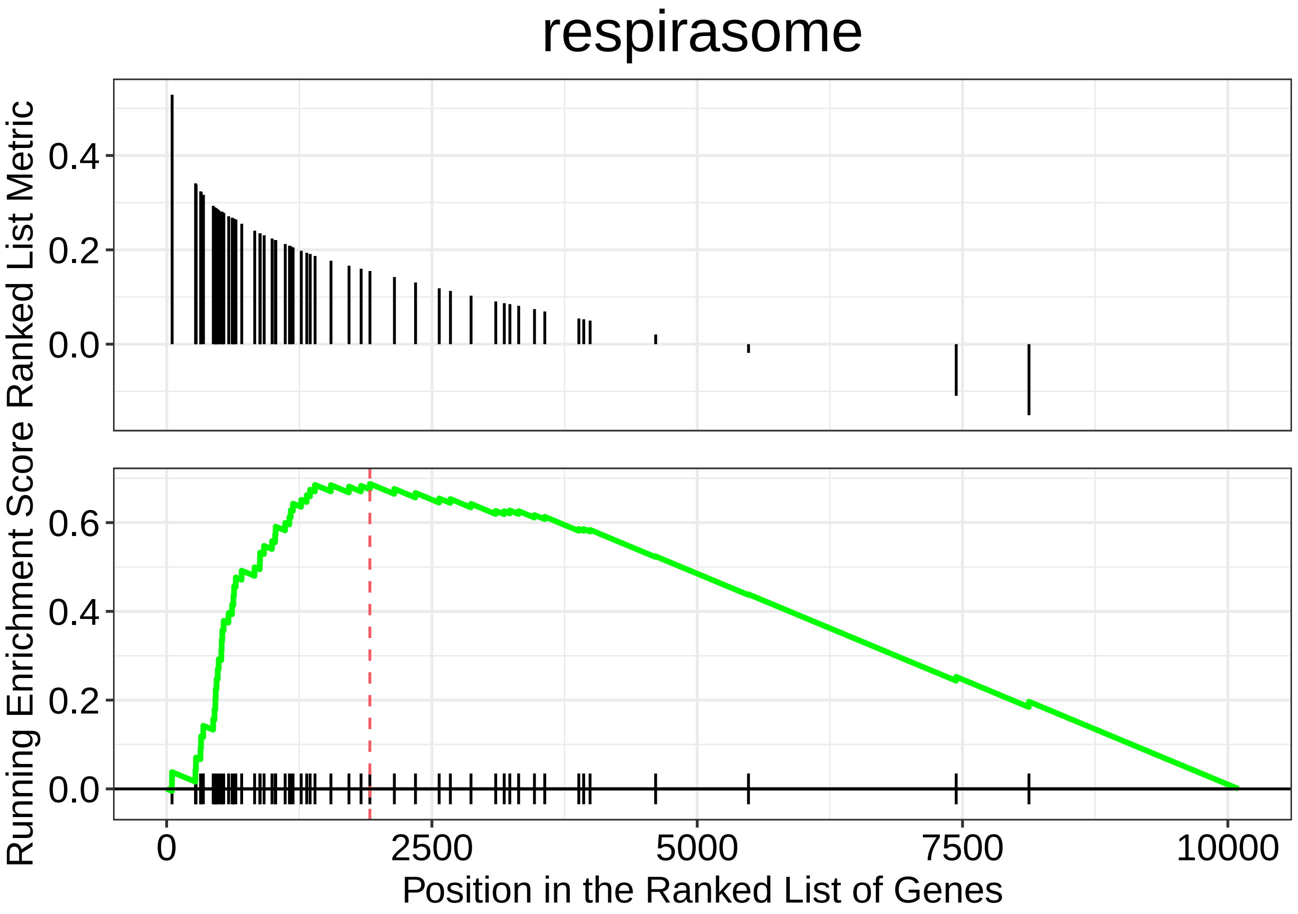

respirasome
Running Enrichment Score Ranked List Metric
0.4
0.2
0.0
0.6
0.4
0.2
0.0
0
2500
5000
7500
10000
Position in the Ranked List of Genes

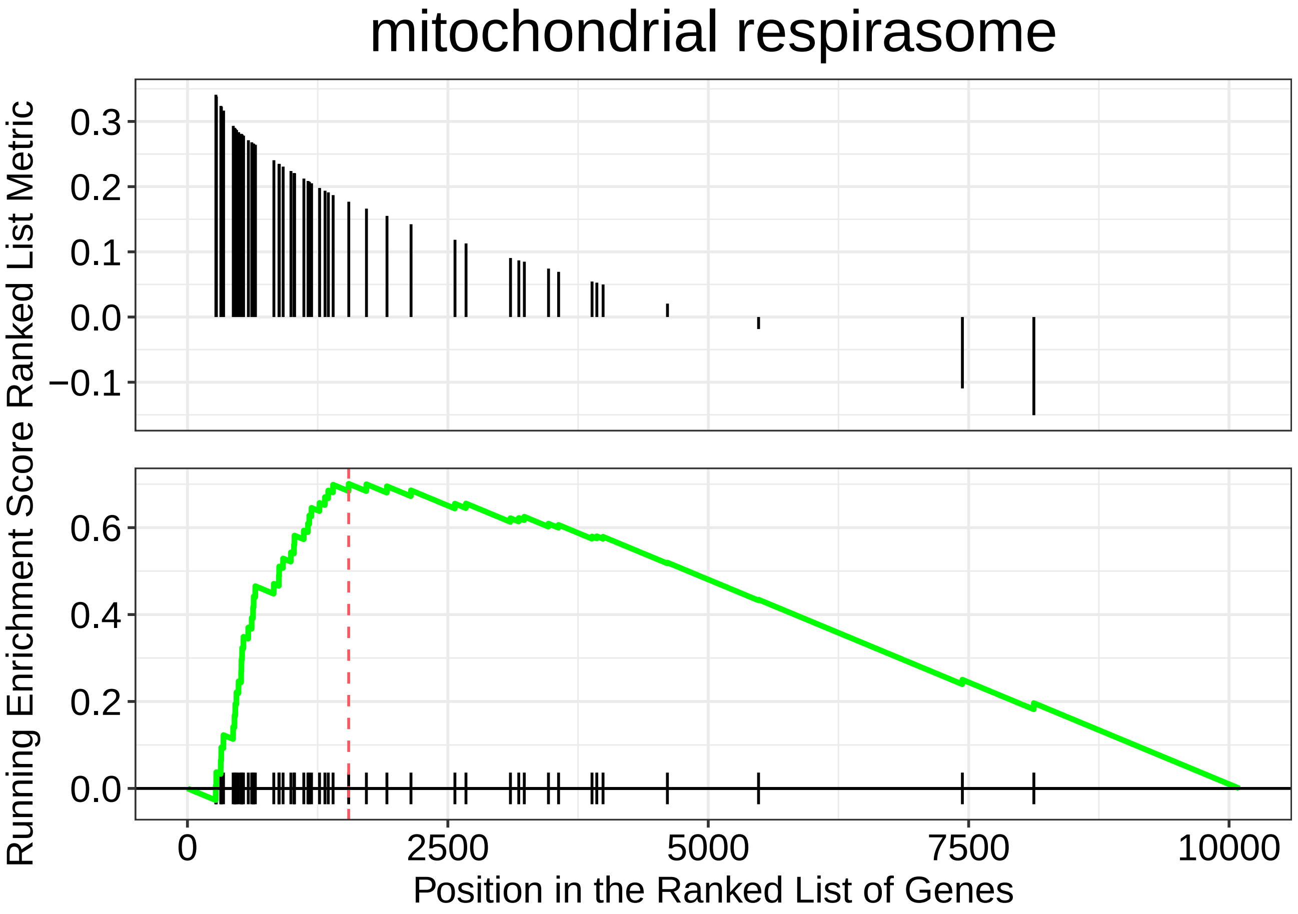

mitochondrial respirasome
Running Enrichment Score Ranked List Metric
0.3
0.2
0.1
0.0
−0.1
0.6
0.4
0.2
0.0
0
2500
5000
7500
10000
Position in the Ranked List of Genes

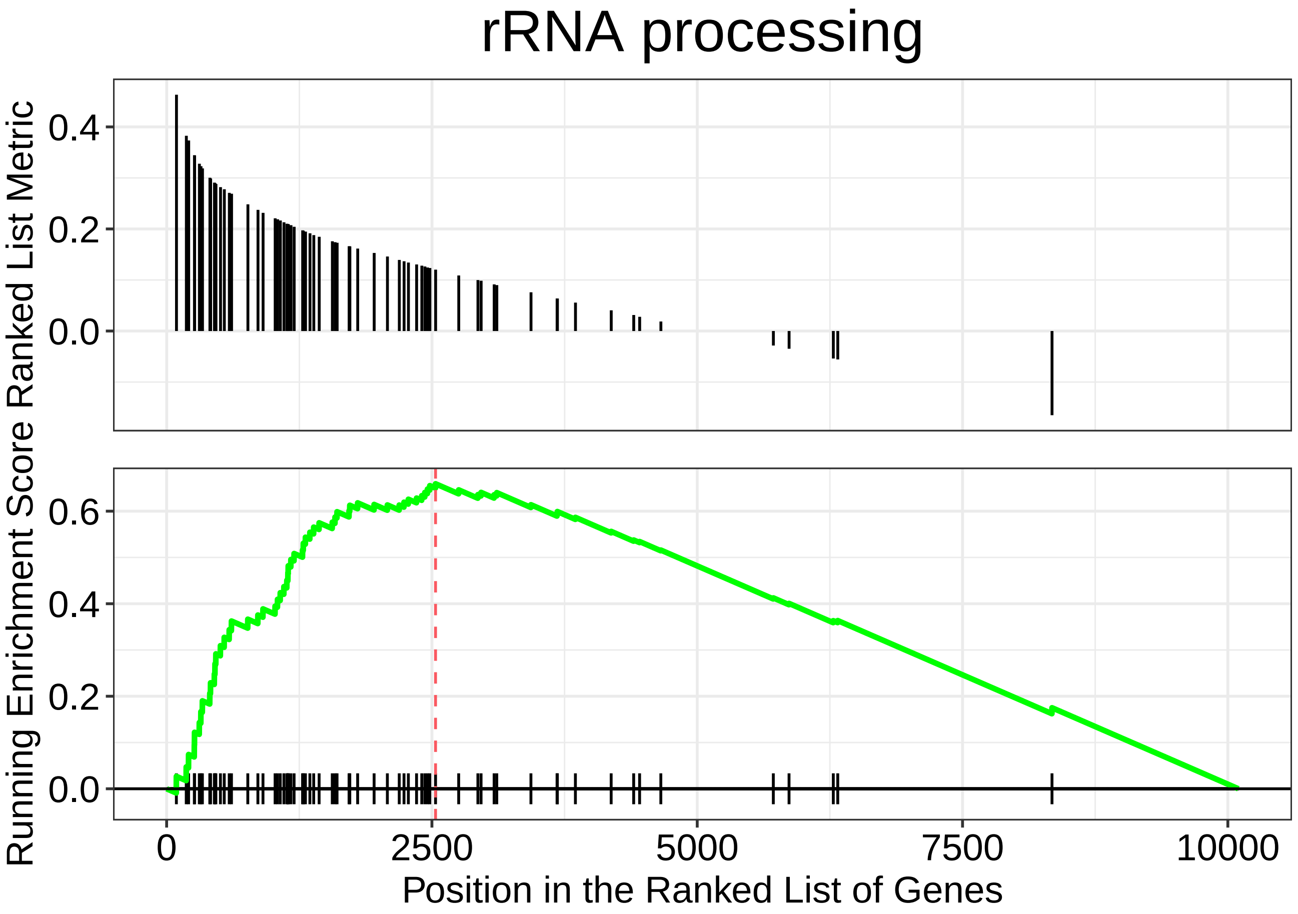
rRNA processing
Running Enrichment Score Ranked List Metric
0.4
0.2
0.0
0.6
0.4
0.2
0.0
0
2500
5000
7500
10000
Position in the Ranked List of Genes

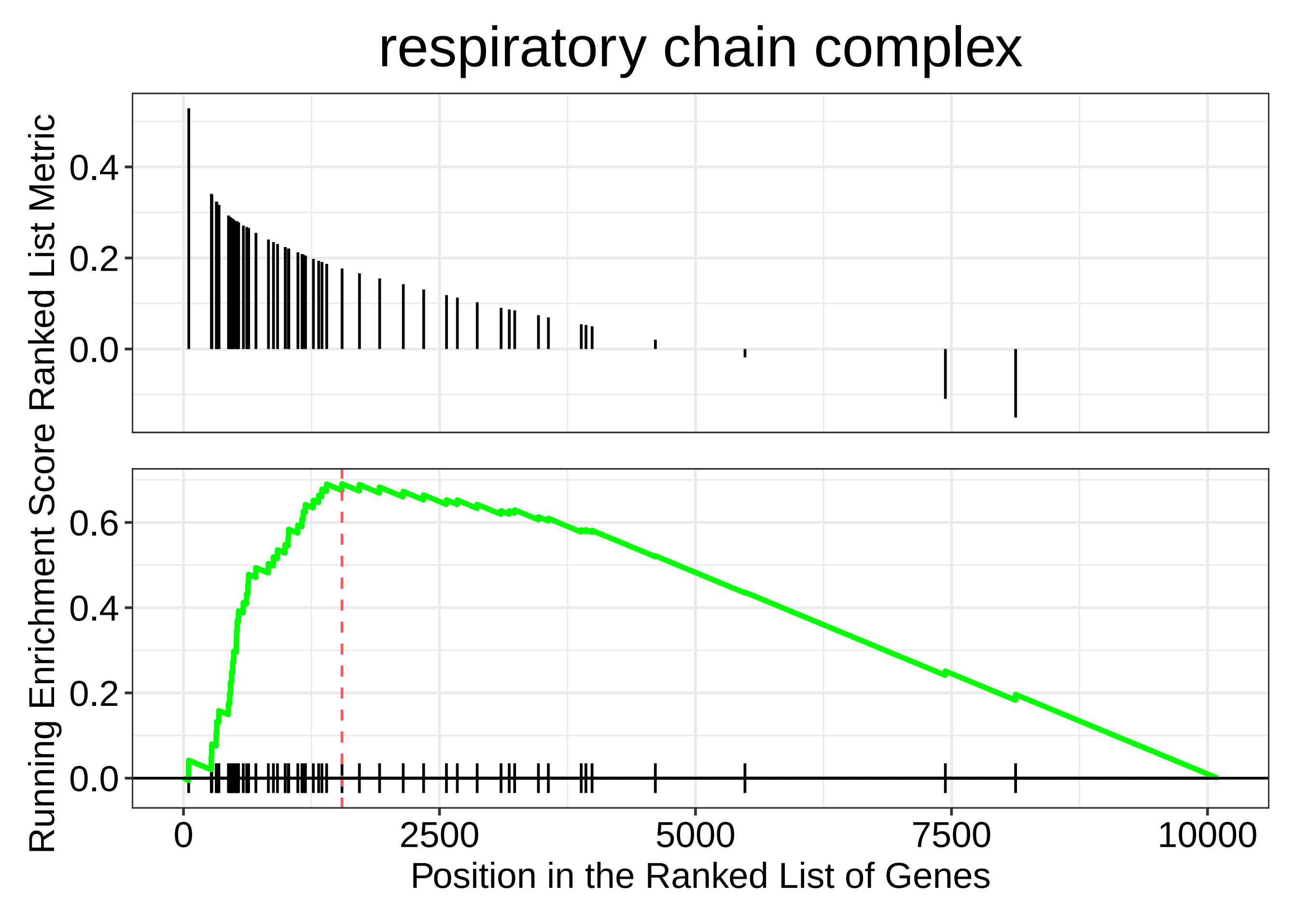

respiratory chain complex
Ranked List Metric
0.4
0.2
0.0
Running Enrichment Score
0.6
0.4
0.2
0.0
0
2500
5000
7500
10000
Position in the Ranked List of Genes
