## Supplementary figures and images for "The small molecule ML233 is a direct inhibitor of tyrosinase function"

### TableS3

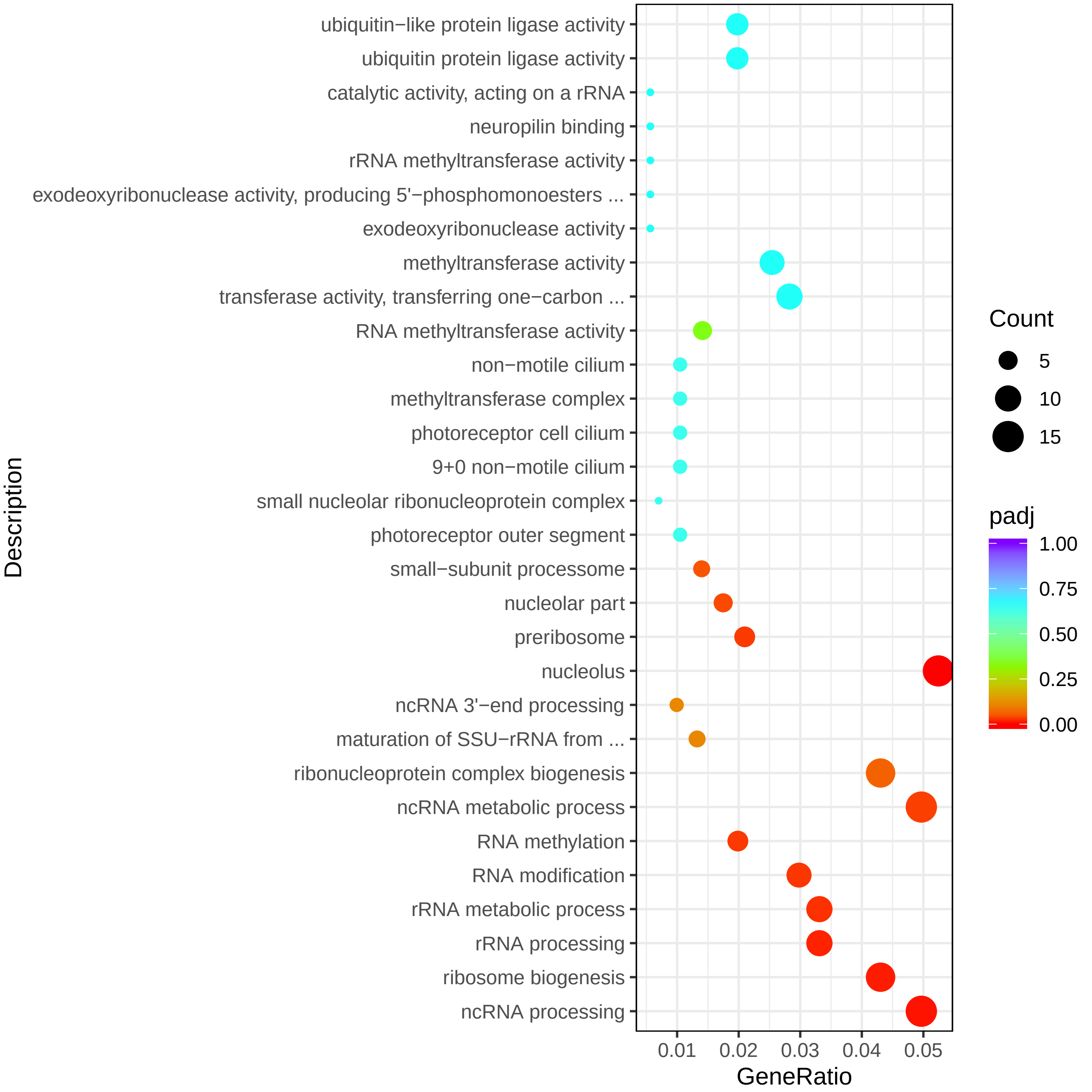

### TableS4

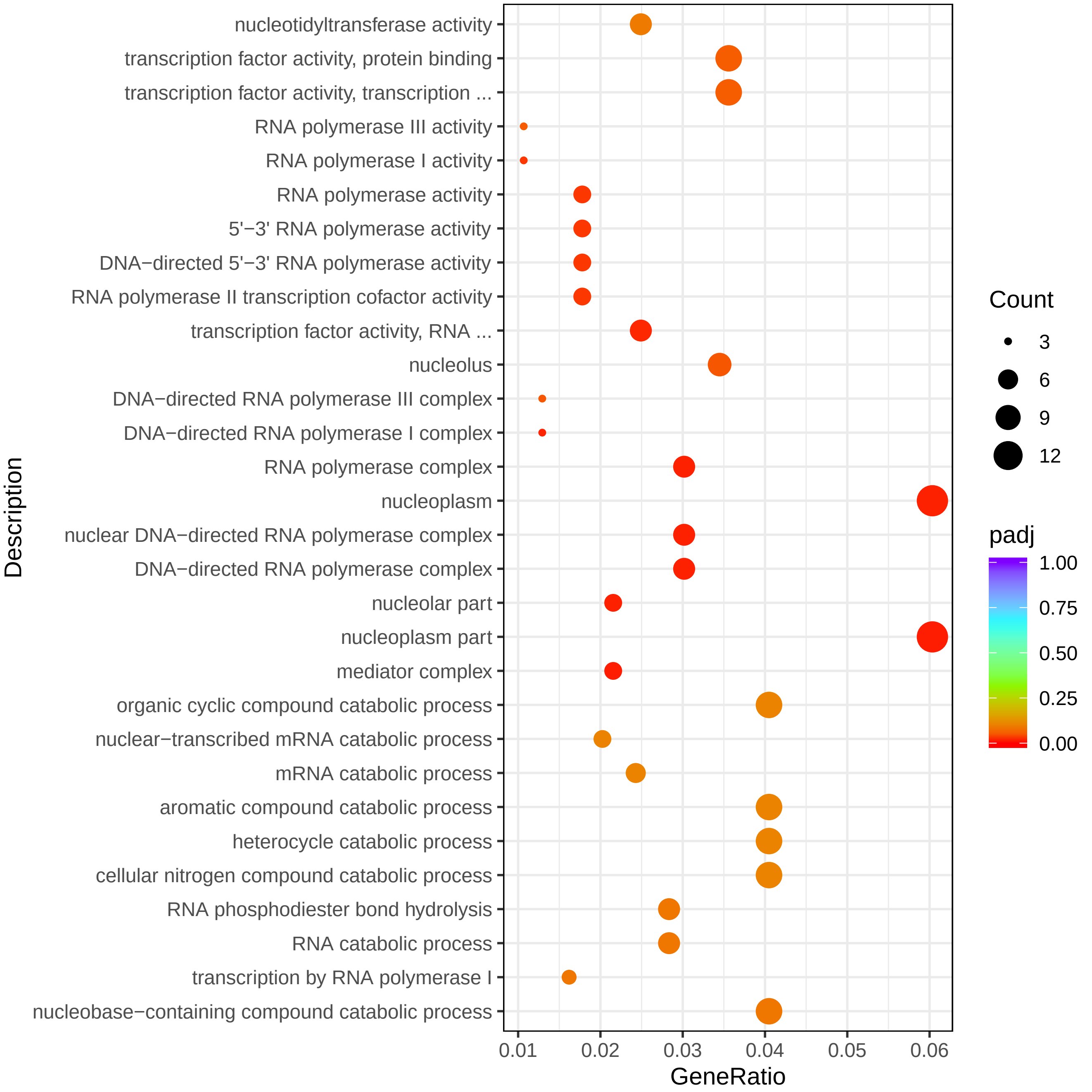

### TableS5

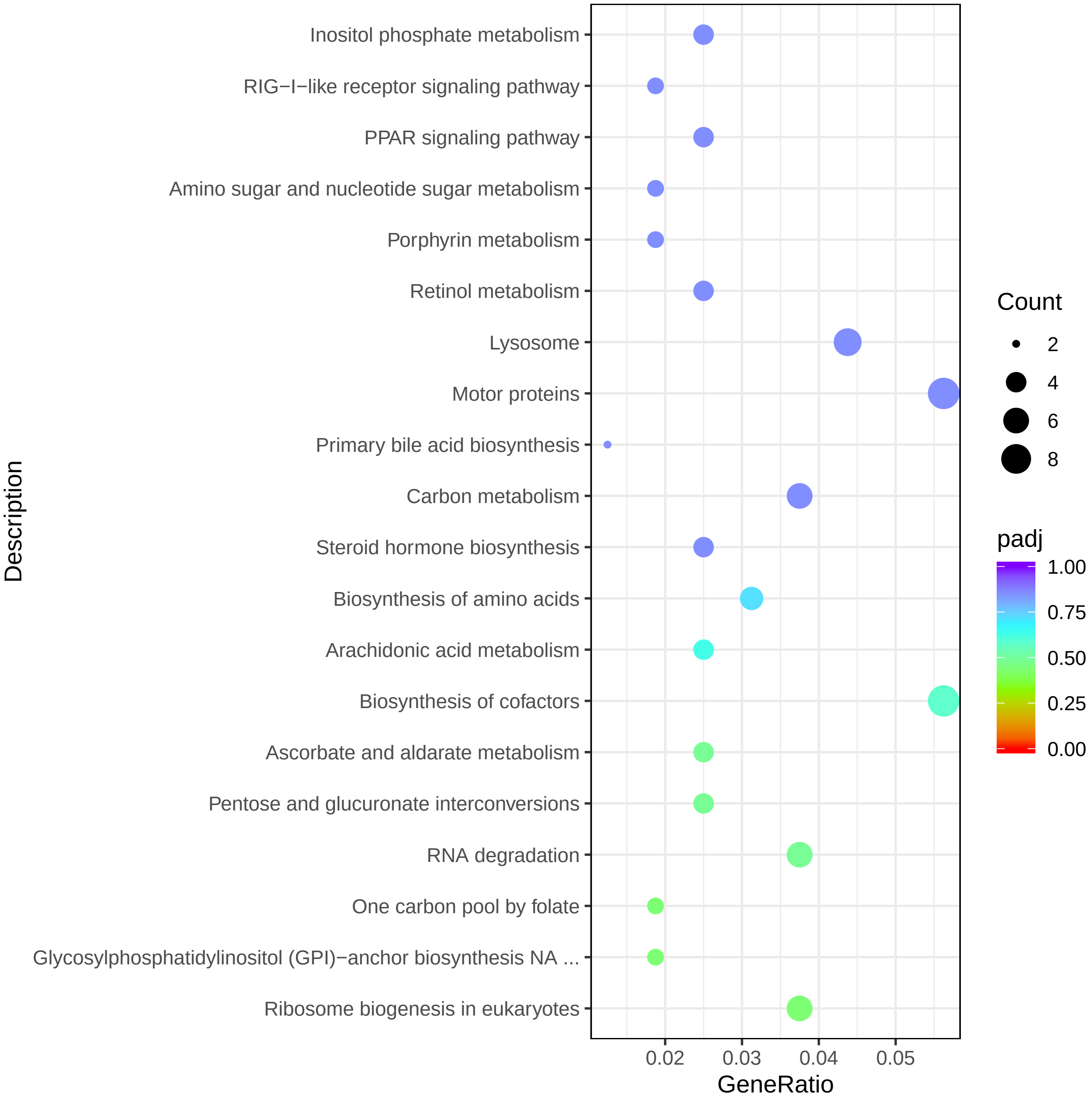

### TableS6

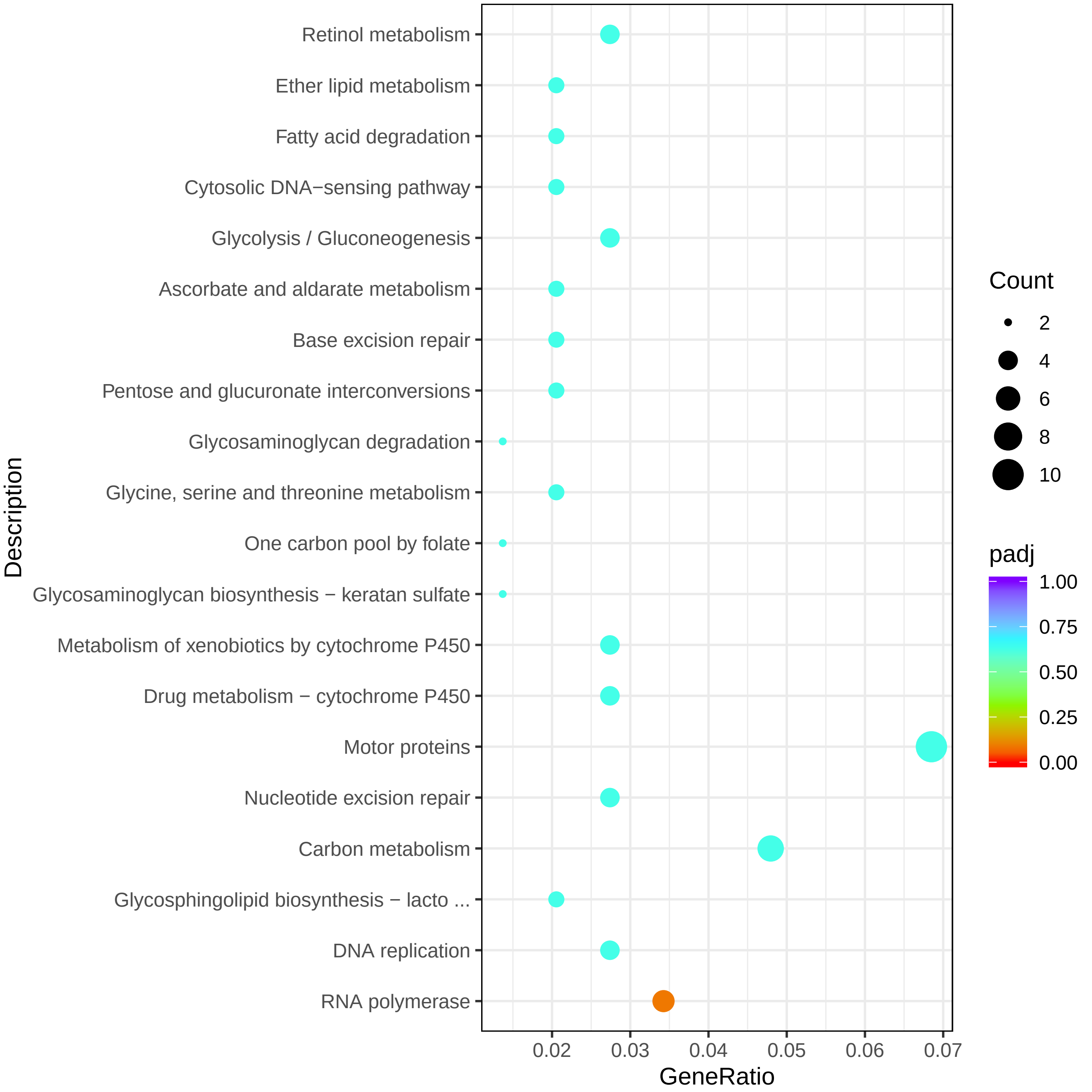
